# The Stochastic System Identification Toolkit (SSIT) to model, fit, predict, and design experiments

**DOI:** 10.64898/2026.02.20.707039

**Authors:** Alex Popinga, Jack Forman, Dmitri Svetlov, Huy Vo, Brian Munsky

## Abstract

Biological data is prone to both intrinsic and extrinsic noise and variability between experimental replicas. That same stochasticity and heterogeneity can carry information about underlying biochemical mechanisms but, if not incorporated in modeling and probabilistic inference, can also bias parameter estimates and misguide predictions and, subsequently, experiment design. Mechanistic inference typically requires lengthy simulations (e.g., the Stochastic Simulation Algorithm (SSA)); approximations to chemical master equation (CME) solutions that lack rigorous error tracking; or deterministic averaging that lacks the complexity necessary to reflect the data. We introduce the Stochastic System Identification Toolkit (SSIT) - a fast, flexible, and open-source software package available on GitHub that makes use of MATLAB’s efficient and diverse computational architecture. The SSIT is designed for building, simulating, and solving chemical reaction models using ODEs, moments, SSA, Finite State Projection truncations of the CME, or hybrid methods; sensitivity analysis and Fisher information quantification; parameter fitting using likelihood- or Bayesian-based methods; handling of experimental noise and measurement errors using probabilistic distortion operators; and sequential experiment design that empowers users to save time and resources while gaining the most information possible out of their data. The SSIT also offers advanced modeling tools, including model reduction methods for increased efficiency and joint fitting of models and datasets with overlapping reactions or parameters. To facilitate the ease and speed of use, the SSIT provides a graphical user interface and ready-made, adaptable pipelines that can be run in the background from commandline or high-performance computing clusters. We demonstrate features of the SSIT on two experimental datasets: the first consists of published mRNA count data that reflect *Saccharomyces cerevisiae* yeast cell response to osmotic shock using single-cell single-molecule fluorescence in situ hybridization; the second consists of single-cell RNA sequencing measurements of 151 activating genes in breast cancer cells following treatment with dexamethasone.

**Author summary:** We present the Stochastic System Identification Toolkit (SSIT) to model, fit, and predict any data that can be interpreted as changing populations or counts through time, including but not limited to single-cell experiments, economics, epidemiology, ecology, sociology, agriculture, and biotechnology. The SSIT was constructed particularly for stochastic modeling, which is important for systems whose states may experience significant fluctuations from mean behavior, thus affecting the inference of the underlying rate parameters and predictions of subsequent behavior. The SSIT provides statistical inference tools for parameter estimation; sensitivity analysis and information calculation; handling of distortions to probability distributions caused by experimental or measurement processes (e.g., dropout in single-cell RNA sequence data and total fluorescence intensities versus spot counting/puncta analysis); and quantitative design of experiments. The SSIT also offers a variety of complex modeling tools, including model reduction methods and fitting of combined models/datasets that share some behavior but remain distinct (e.g., different genes responding a single stimulus). The SSIT generates pipelines for easy, efficient analyses to run in the MATLAB environment, in the background on commandline, or on high-performance computing clusters, thus facilitating users to make informed, time- and cost-effective decisions about their next set of experiments.

## 1 Introduction

The Stochastic System Identification Toolkit (SSIT) addresses the need for a comprehensive software toolkit to analyze and predict noisy and heterogeneous data. Dynamical systems that feature small effective population sizes; population structures that may be multi-level or compartmentalized; slow or varying rates of change; or any other characteristic that can cause substantially divergent individual behavior may require a probabilistic approach to understand and predict its changes through time. Biological systems are especially prone to these effects, particularly at the cellular and molecular levels: gene expression can be intrinsically stochastic, with additional variability arising from cell-to-cell fluctuations in shared factors (often framed as intrinsic vs. extrinsic noise) [**?, ?, ?, ?**]. In contexts such as gene expression, intracellular signaling, and virus–host interactions, the copy numbers of key molecular species are often low (or frequently zero), and the timing of events such as transcription, translation, or degradation depends on factors hidden to the observer [**?, ?**]. Such variability can be informative about the processes that define the system and ignoring it can lead to biased inference results and predictions [**?**]. Additionally, when noise is introduced by particular experimental, measurement, or data processing decisions, special care should be applied to understand and contend with any misleading effects on analyses.

### 1.1 Chemical reaction networks

Any system that can be described by populations that change in size (i.e., counts) in discrete steps that occur at random times can be modeled as a chemical reaction network, which naturally defines a continuous-time Markov jump process on the nonnegative integer lattice. The Chemical Master Equation (CME) is the corresponding forward equation that governs the time evolution of the probability of each count state [**?**], (see an illustration of the CME in Fig. 2).

**Fig 1.**
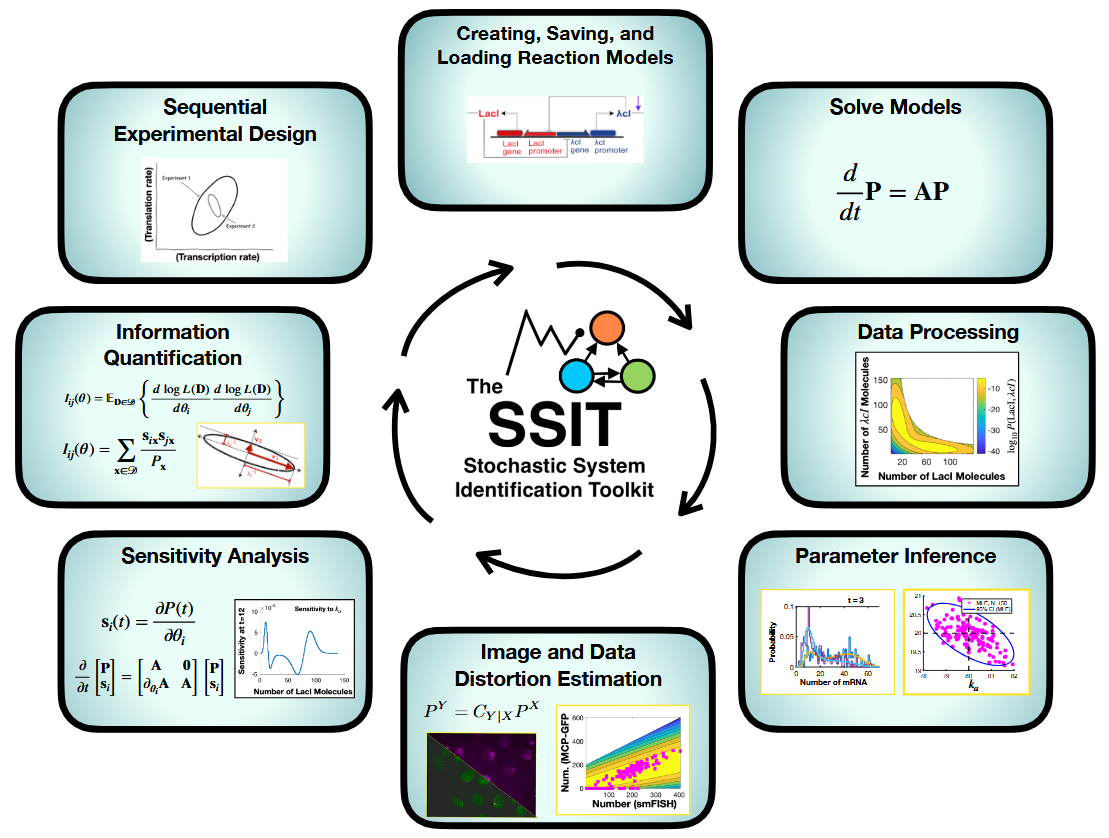
Key features of the Stochastic System Identification Toolkit (SSIT) include (top left and preceding clockwise) creating, saving and loading stochastic models; multiple solution schemes; data processing and parameter inference; image and data distortion estimation; sensitivity analysis; information quantification; and sequential experiment design.

**Fig 2.**
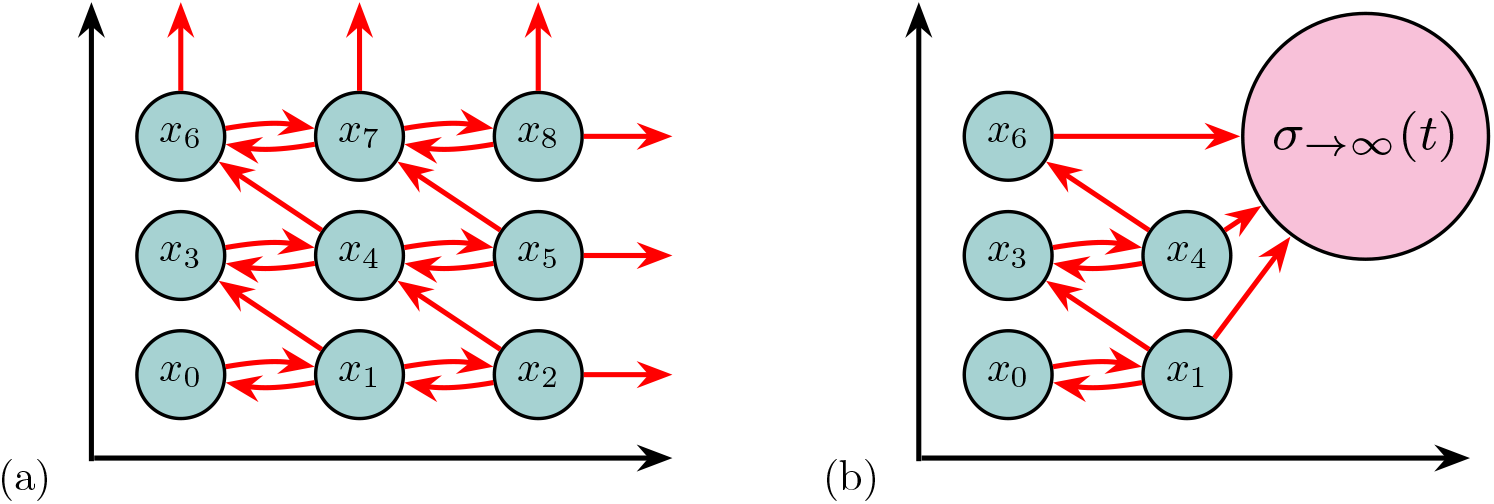
(a) A CME system where *x*_*i*_ are the possible states and each state represents a particular count or counts; (b) an FSP truncation of the CME, i.e., a subset of the full CME for which the probabilities of the system existing in these particular states at specific points in time are explicitly tracked. The probabilities of states outside of the truncated subset are absorbed into a monotonically increasing error sink state *σ*_→∞_(*t*). The total probability absorbed into *σ*_→∞_(*t*) is tracked but not the probabilities of individual states.

Unfortunately, solving the exact CME is intractable for most interesting systems, especially for complex biological systems. The Stochastic Simulation Algorithm (SSA) [6, 7] simulates the same Markov jump process and uses these simulated trajectories to estimate the CME probability distribution by Monte Carlo sampling. However, the SSA is expensive for likelihood-based parameter estimation. Meanwhile, deterministic ODE and moment-based solutions focus on mean concentrations and assume large, well-mixed populations without accounting for random fluctuations/heterogeneity [**?**].

### 1.2 The Chemical Master Equation

To compute direct CME solutions, we can truncate the infinite space of all possible states into a subset of states that retains most of the probability mass. There are different ways to handle the occurrence of a reaction near the boundary of the truncated state space that would move probability from states within the subset to those outside of the boundary. In some approaches, the boundary of the truncated space *reflects* [**?**] or *buffers* [**?, ?**] probability back into the domain when it would otherwise escape, and some approaches such as “slack reactants” *absorb* escaping probability. “Slack reactants” augment the reaction network with “slack” species that have a certain *capacity*; when molecule counts would otherwise exceed a chosen boundary, their “slack” is consumed, thus imposing a conservation measure which forbids the Markov chain from leaving the finite state space [**?**].

Among the many CME projection schemes, the *sink* state method employed by the Finite State Projection (FSP) algorithm [**?, ?**] remains one of the most powerful direct CME approximation. The main advantage of the FSP approach is that it offers a principled way to approximate the CME by projecting the infinite-dimensional state space onto a finite subset, enabling computation of the full probability distribution up to an *exact and computable error bound* [**?**]. In turn, the FSP Algorithm [**?, ?**] uses multiple absorbing sinks to automatically choose an appropriate subspace according to adjustable error thresholds, keeping track of the total absolute error from the loss of probability to states outside of the subspace (shown in Fig. 2).

### 1.3 Finite State Projection

The efficiency of the FSP and other CME truncations can be further improved by several modifications. For example, *sliding window* procedures avoid the necessity for a global boundary condition by evolving the CME on a moving subset [**?, ?**]. Subsequent projections based on timescale separations [**?**], Krylov expansions [**?**], coarse grained state spaces [**?**], or principle orthogonal decompositions [**?**] can also reduce the order of the CME. Tensor train and quantized tensor train methods *compress* both states and operators by representing CME probability vectors and generators as products of “cores”, allowing speedier matrix-vector operations [**?, ?**].

The accuracy and efficiency of the FSP has proven highly useful among computational experts to compare stochastic models to experimental data and make predictions [**?, ?, ?, ?, ?, ?**, 33, 34], yet the approach is often viewed as complicated and has not found wide adoption within the broader community. The practical implementation of stochastic modeling remains a significant barrier for many researchers in computational biology [**?, ?, ?**]. Tools often require specialized knowledge in mathematics or programming, and simulation can be prohibitively slow without optimization or parallelization [**?**]. Although there are many codes available to conduct ODE and SSA analyses of stochastic reaction networks [**?, ?, ?, ?, ?, ?, ?, ?, ?, ?, ?, ?, ?, ?, ?, ?, ?, ?, ?**, 3–5], it can be tricky to write computationally efficient implementations of FSP algorithms, and there are far fewer established, readily available packages that yield efficient FSP solutions. Additionally, many existing software packages lack flexibility, transparency, or the ability to scale to complex biological networks [**?, ?**, 3]. Therefore, there is a growing demand for efficient, accessible, and extensible software frameworks that make stochastic modeling more approachable, particularly for the rapidly expanding fields of bioengineering, biotechnology, and synthetic biology [**?, ?**]. Such tools should support model construction, simulation, inference, and visualization in an integrated and user-friendly environment. They should also enable reproducibility and interoperability with existing standards in systems biology (e.g., SBML [1, 2]).

### 1.4 The SSIT

The SSIT is designed with these considerations in mind and provides fast, transparent solutions for mechanistic models with precise error tracking [**?**], model fitting to experimental or simulated data with parameter estimation by full statistical inference with uncertainty assessment, parameter sensitivity analysis and Fisher information quantification [**?**], extrinsic noise handling by probabilistic distortion operators [34], and sequential experimental design recommendations for stochastic systems [**?**]. For efficient FSP solutions, the SSIT utilizes sparse tensors provided by the Tensor Toolbox [**?**] and Expokit for Krylov-based projection and solution of time-invariant master equations [**?**], along with optional other methods including deterministic ordinary differential equations (ODEs); moment calculations; and SSA [**?**, 6].

Model solutions can then be used to fit a model or models to experimental data. The SSIT provides tools for full statistical inference, including FSP-based likelihood sweeps, maximum likelihood estimation, and Bayesian uncertainty sampling using Metropolis-Hastings [**?, ?**]. Approximate Bayesian Computing (ABC) inference schemes [26, 28] are provided to make use of ODE, SSA and moments-based solution schemes. Model selection can be readily computed using Bayes factors [19], and models can be evaluated for how well they predict held-out data using automated cross-validation [**?**] using the SSIT.

Additionally, the SSIT supplies the machinery for computing the sensitivities of the probabilistic model to small changes in its parameters and Fisher Information Matrices (FIM), which quantifies the amount of information accessible about the values of the model parameters. The SSIT thus allows for ready use of an FSP-FIM [**?**] pipeline for informed sequential design of experiments [**?**].

Finally, the SSIT also offers various more complex modeling options. The SSIT provides probabilistic distortion operators (PDOs) to address measurement errors that result from inaccurate experimental approaches [34]; a multimodel function that combines multiple models and/or datasets to jointly estimate parameters from different experimental conditions or different types of data; and various model reduction approaches to increase efficiency, such as hybrid models that describe a subset of species deterministically and projection schemes based on proper orthogonal decomposition [**?**], eigenvalue decomposition, the quasi-steady-state assumption [**?**], and finite-element based coarse graining [**?, ?**]. The SSIT can be run from command line, within the MATLAB environment, and across high performance computing nodes. A graphical user interface is provided for increased accessibility and ease for novice programmers. The SSIT is open source and available at https://github.com/MunskyGroup/SSIT, built with future modifications in mind, and supports the rapid construction of complex modeling pipelines that can be launched on modern computational research clusters to quickly and simultaneously fit hundreds of combinations of models and datasets.

The remainder of this article is organized as follows: Our results (Section 3) begins in Section 2.1 with an illustration for how to create, save, and load models in the SSIT. In Section 2.2, we discuss various methods to solve models (Sec. 2.2.1-2.2.6), how the SSIT can be used to perform sensitivity analysis on model parameters (Sec. 2.2.7), how to compute Fisher information (Sec. 2.2.8), and how to use Fisher information to design subsequent experiments (Sec. 2.2.9). In Section 2.3, we discuss different techniques for fitting models to data and estimating parameters - including maximum likelihood estimation (Sec. 2.3.3), Bayesian inference (Sec. 2.3.4), Approximate Bayesian Computation (Sec. 2.3.5), and cross-validation (Sec. 2.3.6) - with demonstrations of each feature using a single-cell, single-molecule fluorescence in-situ hybridization (smFISH) experiment in *S. cerevisiae* [33]. Section 2.4 details complex modeling approaches in the SSIT, including model reductions (Sec. 2.4.1), hybrid deterministic-stochastic models (Sec. 2.4.2); probabilistic distortion operators to account for experimental and measurement noise (Sec. 2.4.3); and fitting parameters that are shared across multiple models or datasets (Sec. 2.4.4). The complex modeling methods are illustrated using single-cell RNA sequencing (scRNA-seq) data for 151 genes that are activated in breast cancer cells following the application of a corticosteroid, dexamethasone [29]. Section 2.5 discusses how to execute complex modeling pipelines on a research cluster or local machine. Each of these features is demonstrated through examples and scripts exploring various analyses and parameter estimation for either a four-state model of Mitogen Activated Protein Kinase (MAPK)-induced transcription in yeast or a set of coupled two-state models of glucorticoid receptor (GR)-induced transcription in cancer cells. The key steps taken to create and solve these models are shown in Example Boxes 1 - 23. Detailed example scripts are referred to throughout the text as `example_#_…`) and provided in https://github.com/MunskyGroup/SSIT/tree/main/Examples. Finally, in Section 3, we summarize and discuss the contributions, advantages, and limitations of the SSIT in the context of previous methods and software packages.

## 2 Results

### 2.1 The SSIT Provides Simple Tools to Create, Save, and Load Discrete Stochastic Models

Models of reaction systems can be used to understand how count data changes through time and to gain insight into the mechanisms and rates that drive those changes. A generalized reaction system contains population types, or count types, referred to as **species** and associated with defined **initial conditions** or initial probability distributions; a set of rules that apply changes to the count data referred to as **reactions**; model **parameters** and **input functions** that define various potentially time-varying rates of the process; a set of vectors encoding the updates to the number of each species for each possible reaction called **stoichiometry vectors**; and equations for stochastic reaction rates to define the probabilities that each reaction will occur in the next infinitesimal time step, called **propensity functions**. As a simple example, Fig. 3 illustrates the states, reactions, stoichiometry vectors and propensity functions for the common “Bursting Gene Model”, which consists of a single gene that switches between ON and OFF states to capture the temporal and cell-to-cell heterogeneity often seen in the expression of mRNA or protein molecules in single-cells [**?, ?**].

**Fig 3.**
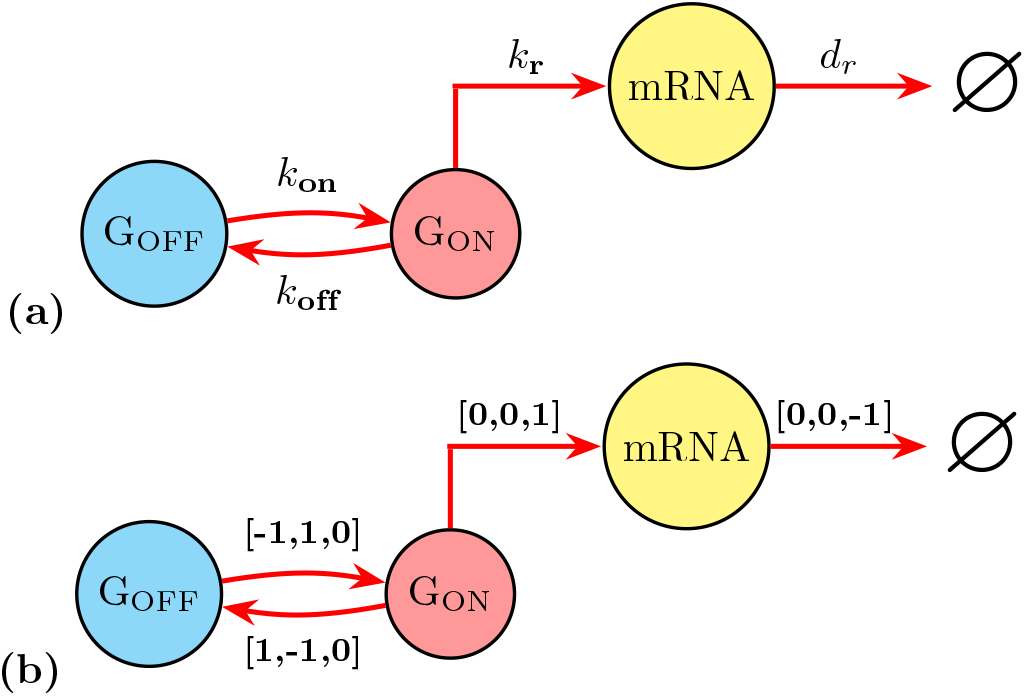
A simple Bursting Gene chemical reaction model. **(a)** There are three **species**: inactive genes (*G*_OFF_), active genes (*G*_ON_), and messenger RNA (*mRNA*). The set containing the number of each species at a given point in time is the state of the system. There are four **reactions**: *G*_OFF_ turns to *G*_ON_, *G*_ON_ turns to *G*_OFF_, *G*_ON_ transcribes *mRNA*, and *mRNA* is degraded (its removal from the system is denoted by the symbol ∅); and each reaction has a rate **parameter**: *k*_on_, *k*_off_, *k*_r_, *g*. Which reaction occurs in the next interval of time is determined by the reaction **propensities**: *w*_1_ ≡ *k*_on_[*G*_OFF_], *w*_2_ ≡ *k*_off_[*G*_ON_], *w*_3_ ≡ *k*_r_[*G*_ON_], *w*_4_ ≡ *d*_*r*_ [*mRNA*]. **(b)** Above each reaction arrow are the **stochiometry** vectors for each reaction representing updates to each species count as a result of the reaction occurring.

A more complex model, along with real experimental data, will be used to demonstrate the utility of the SSIT. We will use the STL1 yeast (*Saccharomyces cerevisiae*) dataset and a model of four distinct gene states (which we will refer to as the “4-state STL1 model”) from Neuert *et al*. [33], described and illustrated in Box 1. The data consist of mRNA counts taken at 15 different time points ranging from *t*_0_ = 0 to *t*_*N*_ = 55(min) following exposure to osmotic shock from the introduction of NaCl.

In the course of this paper, we demonstrate a simple pipeline that loads the dataset into the SSIT; filters the data to focus on desired experimental conditions; associates the filtered dataset with the models we described in Section 2.1 (shown in Fig. 3 and Box 1); performs sensitivity analyses and computation of the Fisher Information; fits the model parameters using maximum likelihood and Metropolis-Hastings; transforms probability distributions of the model parameters that have been distorted by particular experimental design choices; and finally, selecting the next best set of experiments that retain information, reduce uncertainty, and minimize cost.

#### Example Box 1: Schematic of 4-state STL1 model

In this paper, we use mRNA count data from smFISH experiments on the time-varying osmotic stress response of *Saccharomyces cerevisiae* collected by Neuert *et al*. (2013) [33] to demonstrate how to define a model; compute ODE, SSA, and FSP solutions; load and associate experimental data to the model; and fit the model to the data. To present these tools, we use a published 4-state model for the osmotic stress response yeast gene STL1 [33] as shown below:

**Figure B1.**
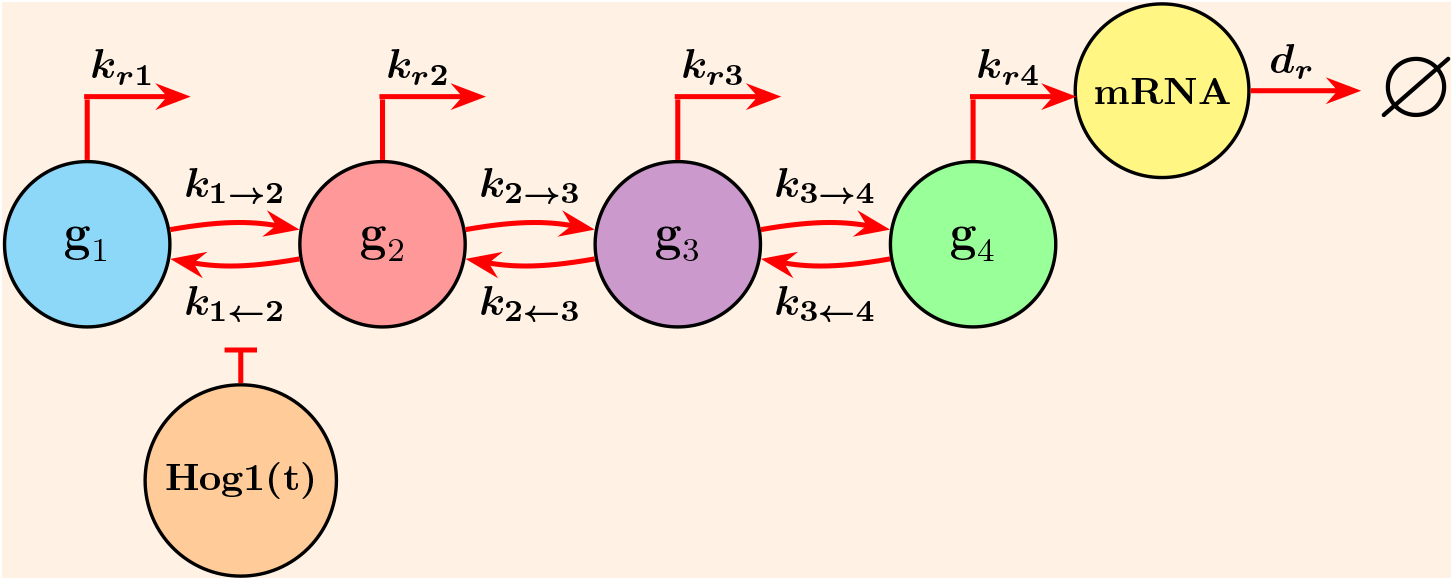
In this expansion of the simple two-state model (Fig. 3), additional states allow for more complex gene activation mechanisms. Transcription occurs in each of the four gene states **g**_1_…**g**_4_ but each transcription reaction has its own unique transcription rate **k**_*r*1_…**k**_*r*4_. Upon activation by osmotic stress caused by a step increase in the concentration of NaCl, the transition **g**_2_ → **g**_1_ is repressed by the presence of the osmotic shock kinase Hog1, which changes in time according to the function **Hog1(t)**. By preventing the system from returning to state **g**_1_, the kinase Hog1 allows the system to transition toward increasingly active states.

To generate a model in the SSIT, first ensure the source code (found in the folder ‘src’) is added to the MATLAB path. This can be done by running the command “install” in MATLAB when in the main SSIT directory. Then, one can either load a pre-built model that has been saved in the SSIT (e.g., ‘BirthDeath’, ‘BurstingGene’, ‘ToggleSwitch’, ‘Repressilator’), or create a model from scratch. In this paper, we will focus on creating the 4-state STL1 model (Box 1) from scratch (Box 2), but both options are shown in `example_1_CreateSSITModels` at https://github.com/MunskyGroup/SSIT/tree/main/Examples.

Once an instance of the SSIT has been created and the model has been named, the following subsections may completed in any order, with their accompanying example boxes: 2.1.1 Define Model Species, 2.1.2 Define a time-varying input signal, 2.1.3 ‘Define Reactions’, and 2.1.4 ‘Define Parameters (Reaction Rates)’. The subsections 2.1.5 ‘Print Model Summary’ and 2.1.6 ‘Save and Load a Model’ are also useful, for, respectively, checking that a model has been implemented as expected and for ensuring that model data is accessible later and/or on another machine.

#### Example Box 2: Add Path and Create Model

By running the installation function `install.m` after cloning the SSIT, the search path should be accurate. Alternatively, ensure that the SSIT source codes are on the search path:

~~~
  addpath(genpath(‘../src’));
~~~

Then, instantiate the SSIT and name the model. Following the 4-state STL1 model described in Box 1, we will name our model ‘STL1 4state’:

~~~
  STL1_4state = SSIT;
~~~

#### 2.1.1 Define Model Species

Specify a name for each model species. The model ‘species’ are the different population types in the system, e.g., different molecules. Refer to Box 3 for an example.

##### Example Box 3: Define Species

The species of our 4-state STL1 model are: ***x* (*t*) = [g**_**1**_ **g**_**2**_ **g**_**3**_ **g**_**4**_ **mRNA]**^⊺^, which in the SSIT can be described by specifying:

~~~
STL1_4state.species = {‘g1’; ‘g2’; ‘g3’; ‘g4’; ‘mRNA’};
~~~

#### 2.1.2 Define a time-varying input signal

The SSIT has an optional property called ‘inputExpressions’ for cases where the system is affected by a time-varying input of some sort (e.g., delivery of a drug, introduction of a chemical via an osmotic source, etc.) An input expression contains rate parameters and relates inputs to particular time points - it may represent a chemical decay process, etc. The 4-state STL1 model has a complex, experimentally-derived signal which is shown in Box 4.

##### Example Box 4: Specify Complex Input Expressions

The complex input expression that best fits the STL1 mRNA count data in our 4-state model (Box 1) is:

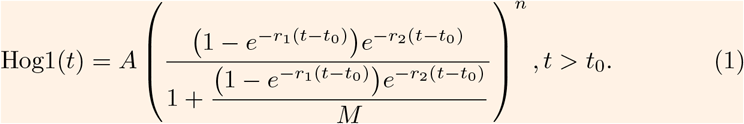

This expression was determined experimentally by Neuert *et al*. [33] and includes an inferrable lag in the parameter *t*_0_. In the SSIT, it is written as:

~~~
STL1_4state.inputExpressions = {‘Hog1’,[‘A*(((1-(exp(1)^(-r1*(t-t0))))*’,…
      ‘exp(1)^(-r2*(t-t0)))/(1+((1-(exp(1)^(-r1*(t-t0))))*’,…
      ‘exp(1)^(-r2*(t-t0)))/M))^n*(t>t0)’]};
~~~

#### 2.1.3 Define Reactions

In the SSIT, reactions are interpreted from defined propensity functions and the corresponding stoichiometry matrix. Reactions can either be added one at a time or in batches as a paired set of propensity vector and stoichiometry matrix. The order of the listed propensity functions matters, as it corresponds to the order of the *columns* of the stoichiometry matrix. Likewise, the order of model species listed (2.1.1) corresponds to the order of the *rows* of the stoichiometry matrix. Stoichiometry matrices define updates to species population vectors that are applied when a reaction occurs.

Box 5 shows an example for defining the reactions of the 4-state STL1 model from Box 1.

##### Example Box 5: Define Propensity Functions and Stoichiometry Matrix

**Propensity Functions**

As there are 11 reactions for our 4-state STL1 model (Box 1), there must be eight propensity functions given in the SSIT. The propensities for our model are:

~~~
STL1_4state.propensityFunctions = {‘k12*g1’;…
      ‘(max(0,k21o*(1-k21i*Hog1)))*g2’; ‘k23*g2’; ‘k32*g3’; ‘k34*g3’;…
      ‘k43*g4’; ‘kr1*g1’; ‘kr2*g2’; ‘kr3*g3’; ‘kr4*g4’; ‘dr*mRNA’};
~~~

**Note:** Propensity functions are compiled into symbolic expressions and saved in separate files, which the SSIT calls when needed. The function called `formPropensitiesGeneral` allows the user to specify the prefix used for saving these propensity function files, which can be very useful for telling the SSIT which files to call when solving a particular model and avoiding errors arising from the SSIT trying to use files that contain propensity functions from an incompatible model. An example of `formPropensitiesGeneral` for our 4-state STL1 model is shown below:

~~~
STL1_4state = STL1_4state.formPropensitiesGeneral(‘STL1_4state’);
~~~

**Stoichiometry Matrix**

There are five species and 11 reactions for our 4-state STL1 model (Box 1). (Note: ‘Hog1’ is not considered a model species but an input expression, Box 4.) The stoichiometry matrix in the SSIT assumes model species by row and reactions by column. The stoichiometry matrix for our model can be entered into the SSIT as:

~~~
STL1_4state.stoichiometry = [-1,1,0,0,0,0,0,0,0,0,0;… % gene state 1
                              1,-1,-1,1,0,0,0,0,0,0,0;… % gene state 2
                              0,0,1,-1,-1,1,0,0,0,0,0;… % gene state 3
                              0,0,0,0,1,-1,0,0,0,0,0;… % gene state 4
                              0,0,0,0,0,0,1,1,1,1,-1] % mRNA
            % Reactions: 1,2,3,4,5,6,7,8,9,10,11
~~~

Another way to add reactions to the SSIT is by using the ‘addReaction’ method. If we wanted to add another ‘mRNA’ degradation reaction, we could do it by:

~~~
newReaction.propensity = ‘dr1*mRNA’;
newReaction.stoichiometry = {‘mRNA’,-1};
newReaction.parameters = {‘dr1’,1};
STL1_4state = STL1_4state.addReaction(newReaction);
~~~

#### 2.1.4 Define Parameters (Reaction Rates)

The reaction rates are the parameters of the model that drive the dynamics of the system and are incorporated in the propensity functions (2.1.3). Box 6 illustrates how to set model parameters, by supplying both a name for each parameter and its value (the units of which are inferred). Parameter values may be later updated by statistical inference during model fitting (Section 2.3). Note: The model parameters do not need to be listed in a particular order since each parameter is given a unique name.

##### Example Box 6: Specify Model Parameters

There are 18 parameters for our 4-state STL1 model, one parameter per each of the 11 reactions in Box 1, with the exception of the reaction **g**_**2**_ → **g**_**1**_.  The transition from state **g**_**2**_ to **g**_**1**_ has a more complex propensity function ‘(max(0,k21o*(1-k21i*Hog1)))\***g**_**2**_’ that splits the simple ‘*k*_2→1_’ parameter from Box 1 into two parameters, ‘*k*_21*o*_’ and ‘*k*_21*i*_’. Additionally, the input ‘Hog1’ decays as a function of time according to our input expression, which requires five additional parameters. (These five parameters - ‘r1’, ‘r2’, ‘A’, ‘M’, and ‘n’ - are experimentally determined and will thus be fixed for model fitting, i.e., parameter estimation.) The 4-state STL1 parameters can thus be specified:

~~~
STL1_4state.parameters = ({‘t0’,3.17; ‘k12’,78; ‘k21o’,1.92e+05;…
     ‘k21i’,3200; ‘k23’,0.402; ‘k34’,7.8; ‘k32’,1.62; ‘k43’,2.28;…
     ‘dr’,0.294; ‘kr1’,4.68e-02; ‘kr2’,0.72; ‘kr3’, 59.4; ‘kr4’, 3.24;…
     ‘r1’,4.14e-03; ‘r2’,0.426; ‘A’,9.3e+09; ‘M’,6.4e-04; ‘n’,3.1});
~~~

To change the value of a parameter that has already been specified, one can simply refer to it by its saved parameter name using ‘changeParameter’, such as:

~~~
STL1_4state = STL1_4state.changeParameter({‘t0’,10; ‘dr’,0.1})
~~~

#### 2.1.5 Print Model Summary

It can be helpful to print a summary of an SSIT model to the screen to check for errors before proceeding to analysis. The function ‘Model.summarizeModel’ (see Box 7) will print model species, reactions, input signals, and model parameters to screen.

##### Example Box 7: Print Model Summary

The following command will print a model summary for our 4-state STL1 model (Box 1), which is shown in the Supporting Information:

~~~
STL1_4state.summarizeModel
~~~

#### 2.1.6 Save and Load a Model

To save a model, along with any associated data and/or solutions, use the ‘save’ function with a named.mat file and the name of the model. To load a previously saved model, use the ‘load’ function with the name of the.mat file and the name of the desired model. Both functions are shown in Box 8.

##### Example Box 8: Save and Load Models

Below, we save the model ‘STL1 4state’ to a file called `example_1_CreateSSITModels`, which will automatically assume the file extension ‘.mat’, and then load it by name.

~~~
save(‘example_1_CreateSSITModels’,’STL1_4state’)
load(‘example_1_CreateSSITModels.mat’)
~~~

#### 2.1.7 Compatibility with SBML and SimBiology

The SSIT has been designed for compatability with the Systems Biology Markup Language (SBML) as well as SimBiology. SBML is an open and collaborative XML-based data format for implementing and sharing computational biology systems. SBML offers a standardized structure for the representation of biological models, especially biochemical reaction networks such as gene expression, metabolic pathways, cell signaling, and epidemics. As with the SSIT, SBML models are defined by their model elements, most commonly the model species, reactions, compartments, parameters, and reaction rules that are defined by stoichiometries and propensities. [1] [2] SimBiology is a MATLAB-based MathWorks toolbox for building, simulating, and analyzing quantitative dynamic models, primarily for applications in pharmacokinetics / pharmacodynamics (PK/PD) systems, by providing for modeling processes such as drug metabolism, efficacy, and safety; signaling pathways; gene expression; and optimization of dosing schedules using ODEs, stochastic simulations, global and local parameter sensitivity analysis, and compartmental modeling. SimBiology is also compatible with SBML, utilizing the standardized format to facilitate collaboration and integration with other modeling platforms. [2] [5]

The Supporting Information provides an example for the simple Bursting Gene model example shown in Fig. 3 and Section 2.1: ‘Creating, Saving, and Loading Models in the SSIT’, in SBML format. Detailed examples for loading SBML and SimBiology models into the SSIT to make use of the SSIT machinery for fast FSP solutions, sequential experiment design, etc., are provided in the Supporting Information.

### 2.2 The SSIT provides numerous tools to find and and visualize master equation solutions

Once the model reactions have been specified, one can then use a variety of methods to solve for population sizes over time. These solutions can then be used to estimate reaction rate parameters from experimental data.

The SSIT allows users to compute deterministic ordinary differential equations (ODEs); closed moment solutions; Stochastic Simulation Algorithm (SSA) trajectories; efficient solutions to the Finite State Projection (FSP) approximation of the full chemical master equation (CME); and hybrid deterministic-stochastic solutions.

Box 9 shows two arguments for setting up for any type of model solution.

#### Example Box 9: Set up for Solutions

Solutions will be evaluated at specific time intervals. The SSIT has default time points for computing solutions given by ‘linspace(0,10,21)’, but the property ‘tSpan’ can be used to set alternative times:

~~~
STL1_4state.tSpan = linspace(0,50,101);
~~~

(The ‘linspace’ function returns a linearly spaced vector such that, for the above example, 101 time points are generated with even spacing between 0 and 50.)

Additionally, we specify the initial condition, i.e., counts for each model species at the start of the simulation or solution. The SSIT default initial condition of ‘[0]’ would only work for a single-species model, and even then may or may not be appropriate depending on the model. For our 4-state STL1 model (Box 1), each of the five species are given an initial count:

~~~
STL1_4state.initialCondition = [1;0;0;0;0];
~~~

The SSIT allows for the initial condition to be probabilistic by specifying the ‘initialCondition’ property as a set of possible states and the ‘initialProbs’ property as a vector of the corresponding probabilities. For FSP implementations, the initial condition can also be set as the steady state probability distribution assuming rate constants that are fixed at a user-specified time.

#### 2.2.1 The Chemical Master Equation (CME)

Discrete state stochastic processes describe a system that evolves through time with some degree of randomness, as its transitions between countable states proceed in time under the governance of probabilistic rules [**?**, 9]. These processes are characterized by a sequence of random variables, typically indexed by integers, where each variable represents the state of the system at a particular time [**?**]. Discrete state stochastic processes are common models for systems whose states, or counts, are updated according to probabilistic rules at distinct fixed or random time intervals [**?, ?**, 7].

The CME is composed of a set of linear ordinary differential equations where the probability mass *p*(**x**_*i*_, *t*) is described through time *t* for all states of the system **x**_*i*_ [8]. A reaction *µ* ∈ {1, …, *R*} occurs at time *t* probabilistically according to the reaction propensities, or rates, determined by function *w*_*µ*_(*t*, **x, *θ***), with model parameters ***θ*** = [*θ*_1_, *θ*_2_, …, *θ*_*n*_] ∈ ℝ_≥0_. A reaction in the CME is treated as a Markovian jump (illustrated in Fig. 4 for a 3-species model) that moves the system between the two different states. The states **x** ∈ 𝕏 in the CME represent the counts of each molecule (species) type, so for a single-species model **x** ∈ 𝕏 is represented as a scalar, where 𝕏 ⊆ ℕ_≥0_ and for multi-species models with *d* species, **x** is a d-element vector and 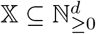. The new state at time *t* + *ϵ*, to which the system moves when a reaction occurs, is updated according to

**Fig 4.**
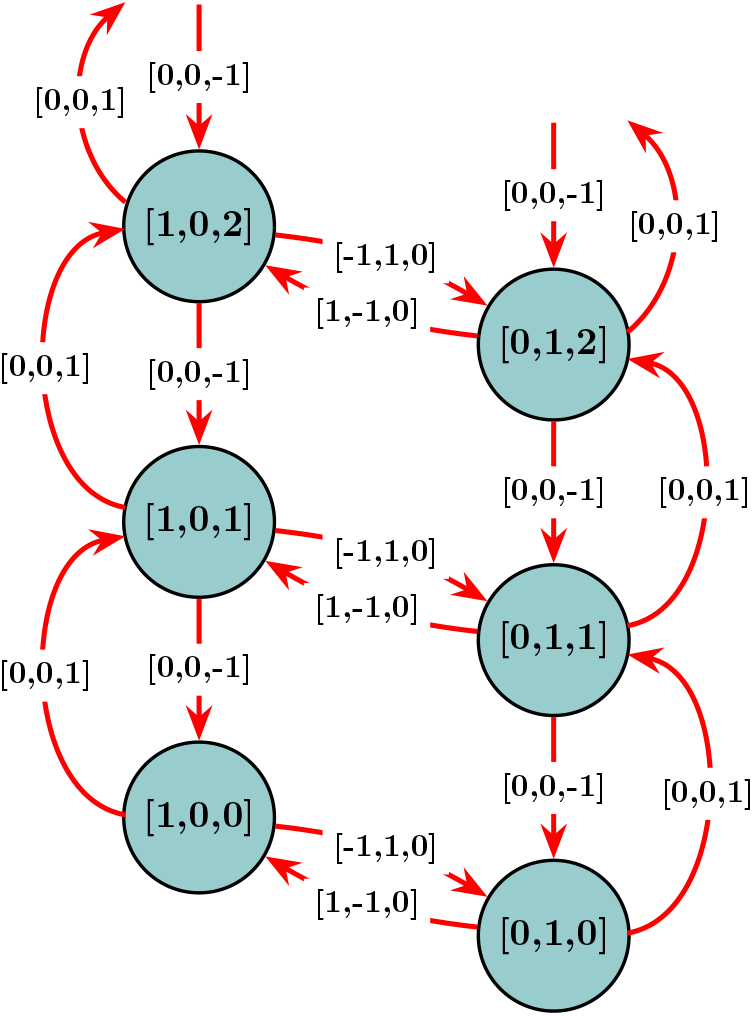
The Bursting Gene Model from Fig. 3 represented as a discrete state Markov model. The model has three species: a gene in its inactive form (“G_OFF_”), the activated gene (“G_ON_”), and mRNA. For this simple model, the gene only switches states between “G_OFF_” and (“G_ON_”), but there can, in theory, be any positive number of mRNA molecules.

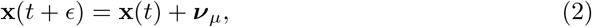

where ***ν***_*µ*_ is the stoichiometric vector coinciding with reaction *µ* ∈ [0, 1, …, *M*]. [34] The probability mass *p*(**x**_*i*_, *t*) is thus updated for each interval of time such that:

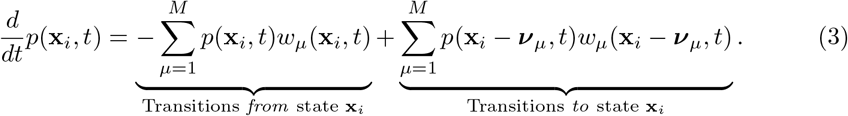

The probability mass vector for the whole state space of the CME can be written as

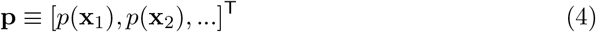

[8], and an *infinitesimal generator matrix* **A** can be constructed to capture the propensity sums, where *A*_*i,j*_ denotes the (*i, j*)th scalar entry of **A** such that

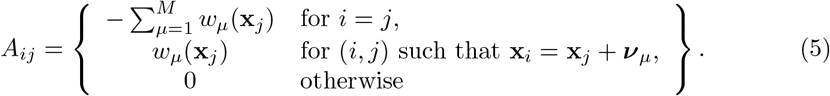

Therefore, the CME of Equation 3 can be reformulated more concisely as

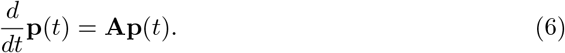

The CME is a useful method for probing stochastic discrete state time evolutions. However, solving a full CME analytically is only possible for simple, often uninteresting, systems, and the combinatorial explosion of possible states in more complex, realistic models renders exact solutions infeasible.

#### 2.2.2 Ordinary Differential Equations (ODEs)

Ordinary differential equations (ODEs) may be used for species whose populations are large enough that the changes in their sizes may be approximated by averaging over time; the sums of their changes from their initial states are sufficient due to a lack of stochasticity [**?, ?**, 11].

The Chemical Master Equation (CME) solution can be approximated using first-order ODEs to estimate the expected changes to all species in the state vector, i.e., 𝔼{**x**} with respect to time such that:

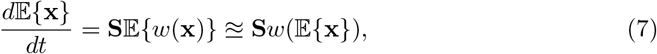

where **S** represents the stoichiometry matrix and *w*(**x**) = [*w*_1_(**x**, *t*), …, *w*_*N*_ (**x**, *t*)]^T^ is a vector containing propensities for the potential reactions of the system [**?**, 11]. This approximation is exact in the case where *w*(**x**) is constant or linear in **x**, such that 𝔼 {*w*(**x**)} = *w*(𝔼 {**x**}). The matrix-vector product **S***w* sums over all changes to **x** that may occur according to all possible reactions (i.e., summing over each reaction’s propensity and stoichiometric update) with respect to *t*. The Supporting Information shows the ODEs in component form for the simple Bursting Gene model (Fig. 3) and 4-state STL1 model (Box 1).

An example for solving the 4-state STL1 ODEs and plotting their solutions is provided in Box 10 and in more detail in `example_2_SolveSSITModels_ODE.m`, which also contains examples for the simple Bursting Gene model (Fig. 3) and a simplified STL1 model for comparison.

Note: An ODE solver is used to solve the system of ODEs. MATLAB provides a suite of precise, fast ODE solvers for both stiff and non-stiff solutions, e.g., ode23s and ode45 [**?**], which are readily employable in the SSIT by user specification. When propensity functions are defined as continuous and differentiable functions, the SSIT symbolically computes the Jacobians for use in ODE solvers; this allow for significant computational savings for numerically stiff systems.

##### Example Box 10: Deterministic Solutions (ODEs)

We demonstrate how to solve ODEs for the 4-state STL1 model (see Box 1) in the SSIT below:

~~~
% Set solution scheme to ODEs:
STL1_4state.solutionScheme = ‘ODE’;
% Solve ODEs:
STL1_4state.Solutions = STL1_4state.solve;
% Plot ODE solutions for mRNA:
STL1_4state.plotODE(STL1_4state.species(5), STL1_4state.tSpan, {‘linewidth’,4},…
    TitleFontSize=26, Title=‘4-state STL1 (mRNA)’, AxisLabelSize=20,…
    TickLabelSize=20, LegendFontSize=20, LegendLocation=‘east’,…
    Colors=[0.23,0.67,0.20], XLabel=‘Time’, YLabel=‘Molecule Count’)
% Plot ODE solutions for the four gene states:
STL1_4state.plotODE(STL1_4state.species(1:4), STL1_4state.tSpan, {‘linewidth’,4},…
     TitleFontSize=26, Title=‘4-state STL1 (gene states)’, AxisLabelSize=20,…
     TickLabelSize=20, LegendFontSize=20, LegendLocation=‘east’,…
     XLabel=‘Time’, YLabel=‘Molecule Count’)
~~~

The ODE solutions for the five species in the 4-state STL1 model are shown below. Note: We plot the full time interval *t* ∈ [0,50] and acknowledge that the dynamics for *t* ≤ 1 are dominated by a stiff fast mode that quickly decays, after which the slower gene-state dynamics of interest become clearly visible.

**Figure B10.**
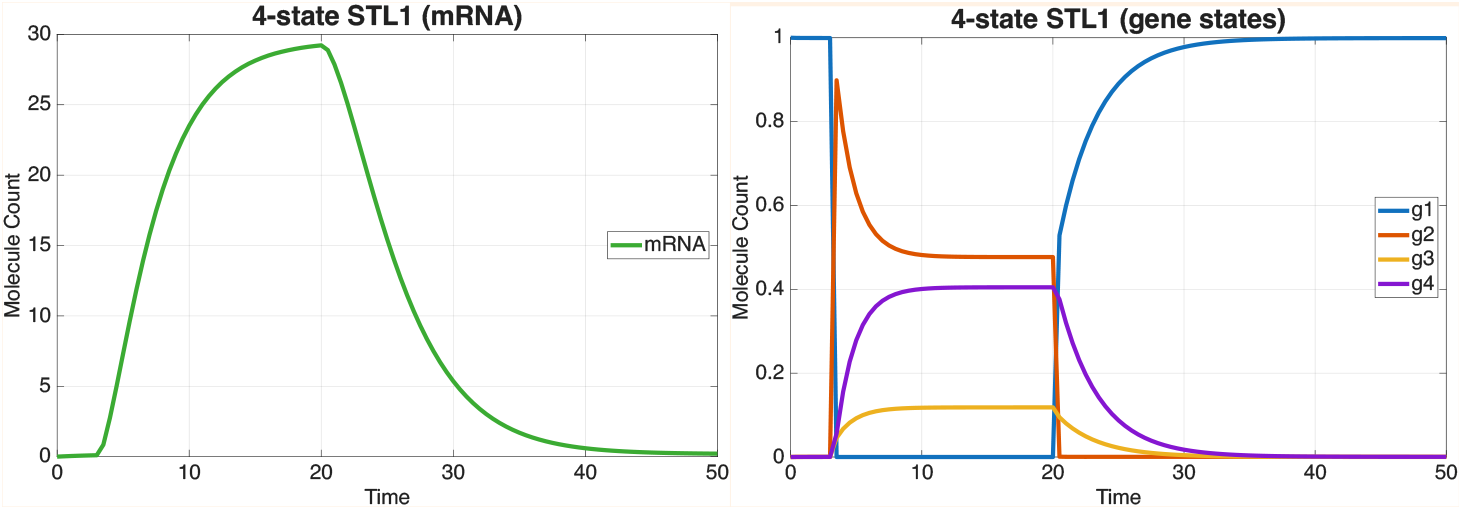
ODE solution plots for the 4-state STL1 model from Box 1 showing the average number of mRNA (left) and the fraction of cells in the **g**_1_ − **g**_4_ states (right).

#### 2.2.3 Moment Closure

Moment-based approaches are efficient ways approximate the time evolution of the system using low-order statistics by deriving exact or approximately closed moment solutions from the Chemical Master Equation (CME), assuming functional dependence between all cumulants of the probability distributions (the mean, variance, and so forth). For linear propensities, the moment closure is exact, and for non-linear reaction networks, the closure bases the approximation on a proposed distribution or generating function (e.g., Gaussian, log-normal, cumulant, or entropy-based), procuring solvable ODE systems [**?**]. The closure thereby posits a parametric form or constraint in place of the unknown higher moments. However, closures rely on assumptions [**?**] that may be inappropriate in some cases and lead to inaccurate predictions, e.g., when Gaussian assumptions are applied to distributions that are skewed, multimodal, or heavy-tailed [**?, ?**].

The SSIT offers the ability to compute simple closure approximations. For example, if the joint state distribution can reasonably be assumed as multivariate normal, then the CME might be approximated using Gaussian closures by calculating all third and higher-order moments using the means, covariances, and the assumption of a multivariate normal distribution. An example for computing moment solutions from our 4-state STL1 model (Box 1) is provided in the Supporting Information and `example_18_Moments.m`.

#### 2.2.4 Stochastic Simulation Algorithm (SSA)

Although efficient, continuous, and deterministic solutions to estimate average or normally-distributed system states as a function of time may be inappropriate when the system is built on small, discrete populations whose population sizes change dynamically through time and exhibit highly variable and non-Gaussian behavior - as is the case at the level of gene expression and cell signaling, for examples, where the numbers of species in the model may be as low as single molecules of mRNA or proteins [**?, ?, ?**]. In such cases, the stochastic fluctuations of the model species populations may become significant and unpredictable by deterministic averages or moment closure models [**?, ?**].

The Stochastic Simulation Algorithm (SSA) [6, 7] is one of the most common methods for exploring the dynamics of such reaction networks, computing relevant statistics, and validating mechanistic models. The SSA simulates exact sample paths through reaction networks, probabilistically selecting reactions to occur in a given time interval based on propensities and applying stoichiometric updates to the system states. The SSA is simple to implement and, unlike methods which use ODEs to average over population changes with respect to time, the SSA provides Monte-Carlo estimates and can yield important insights regarding low-probability events and heterogeneous behavior.

The SSA is precise in that it tracks every individual reaction over time and updates the resultant states accordingly [**?**]. A collection of many SSA trajectories (details about generating SSA trajectories are provided in the Supporting Information) may be computed by simulating the procedure with newly generated random numbers for each new trajectory, and subsequent statistical analyses may be performed on these trajectories, including computing the likelihood of the data. unfortunately, the SSA does not scale well in efficiency when required to track large numbers of reactions or when a high level of accuracy is needed to examine rare events within a population. Time-leaping methods such as the *τ* -leaping method [**?, ?**] approximate the solution by incrementing a specified interval of time and grouping reaction events within the time intervals. Naturally, these methods are more efficient but risk missing key reactions and result in a substantial loss of accuracy for systems containing low species copy numbers or propensities that lead to rapid, large changes of species counts.

The SSIT solver for the SSA supports parallelization and GPU use on capable devices. The SSIT also makes use of a modified version of the Extrande [**?**] method to handle time-varying reaction rates in the system propensity functions. Box 11 and `example_3_SolveSSITModels_SSA.m` illustrate how to use the SSIT to simulate SSA trajectories.

##### Example Box 11: Generate Stochastic Trajectories with the SSA

Here, we demonstrate how to use SSA in the SSIT for the 4-state STL1 model (Box 1). Plots of the SSA results are shown in Fig B11.

~~~
% Set solution scheme to SSA:
STL1_4state.solutionScheme = ‘SSA’;
% Set the number of simulations performed per experiment:
STL1_4state.ssaOptions.nSimsPerExpt=100;
% Use a negative initial time to allow model to equilibrate to an initial
% steady state prior to starting the subsequent simulation (burn-in):
STL1_4state.tSpan = [-100,STL1_4state.tSpan];
% Set the initial time:
STL1_4state.initialTime = STL1_4state.tSpan(1);
% Run iterations in parallel with multiple cores, or execute serially:
STL1_4state.ssaOptions.useParallel = true;
% Run SSA:
STL1_4state.Solutions = STL1_4state.solve;
~~~

**Figure B11.**
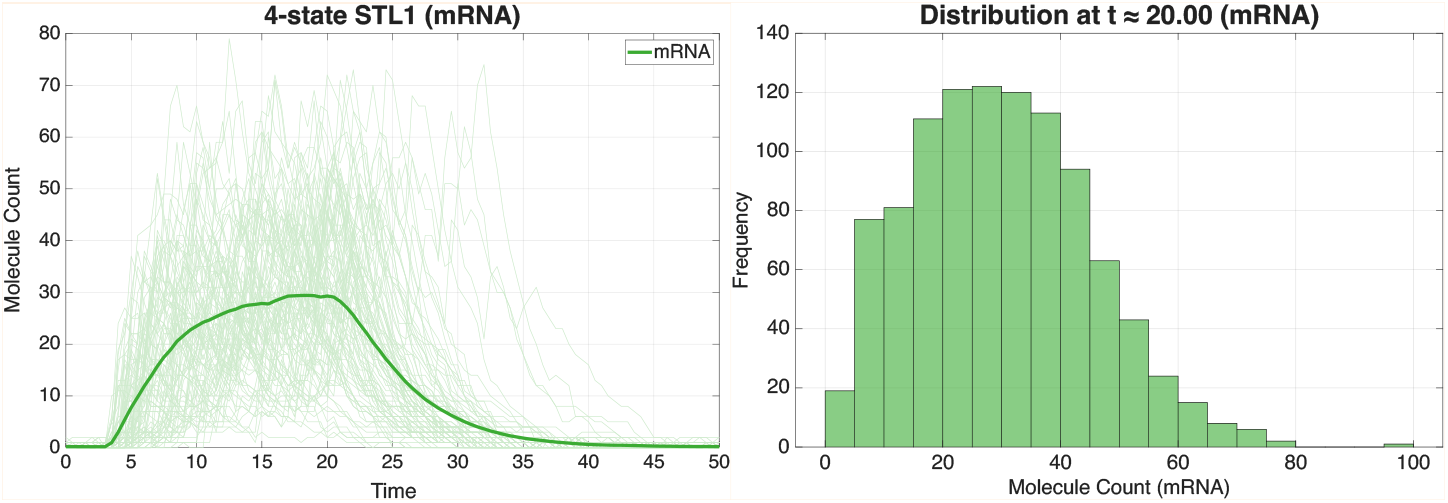
SSA means and trajectories for the 5 species of the 4-state STL1 model from Box 1 (left) and means only, zoomed in on the 4 gene states (right).

#### 2.2.5 Finite State Projection (FSP)

The Finite State Projection (FSP) algorithm [**?**] yields an exact analytical solution in the case of finite distinct population vectors and closely approximates a reduced, finite space of an otherwise possibly infinite set of states while simultaneously calculating the error of its approximation. The FSP is precise and fast under the SSIT implementation in MATLAB, and thus has proven highly valuable in the solving of CMEs for single-cell data [**?, ?, ?, ?**, 8].

The FSP algorithm divides the total states of the CME into two categories: a set of states determined to be most realistically (most probable to be) visited over the time interval of interest, for which the FSP algorithm provides solutions, and one or more absorbing, sink states that represents error (see the state labeled “*σ*_→∞_(*t*))” in Fig. 2). Probability flows between the states of interest and into the error sinks, from which it does not return. The FSP then solves a subset of linear ordinary differential equations that provide the solutions to the evolution of the probability density vectors relevant to the FSP states, a subset of the whole CME. In doing so, FSP provides efficient results and a lower bound to the exact solution [**?, ?**].

Formally, FSP solutions are computed by first defining all enumerations over all possible states reachable by the system as: {**x**_1_, **x**_2_, …} ⊆ 𝕏. Next, a finite set of indices *J* = {*j*_1_, *j*_2_, …, *j*_*N*_} is specified and used to determine a finite set of states 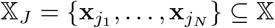. Naturally, the complement set 𝕏_*J*_*′* contains all of the states that were *not* included in 𝕏_*J*_. The full CME can then be rearranged to write as:

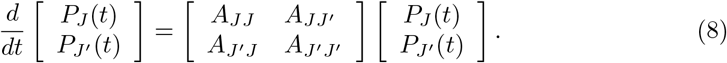

The most defining feature of the FSP approach is that the second set of states is replaced by one or more absorbing sinks that add to form 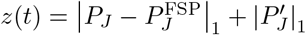, into which the probabilities of system states flow but are no longer individually tracked and therefore are lost from the finite state set 𝕏_*J*_ [8]. The total probability mass that ends up in the subset of states *z*(*t*) is accounted for as the total error - loss of probability mass as a function of time from the finite state truncation 𝕏_*J*_ of the CME - and this formalization gives rise to the finite dimensional master equation:

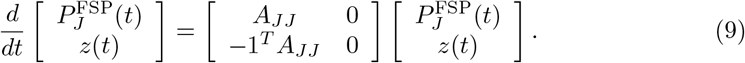

The solutions of the FSP algorithm yield several notable results:

a. The FSP provides a strict lower bound on the exact CME solution 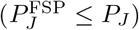 for every possible state of the process;
b. The FSP provides the exact total absolute error of the finite approximation of the full CME, 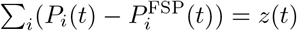
c. The error resulting from CME truncation, *z*(*t*), decreases monotonically with each additional state that is added to the set 𝕏_*J*_;
d. The exact cumulative probability distribution for the time to escape any finite set 𝕏_*J*_ is given by *z*(*t*).

Characteristics (a)-(c) of the FSP approximation listed above enable the setting of a maximum tolerable CME error as a monotonically increasing function, *ε*(*t*) *>* 0, where *ε*(*t*_1_) ≤ *ε*(*t*_2_) for every *t*_2_ *> t*_1_. The FSP algorithm is then implemented so that it proceeds through the following steps [**?, ?, ?**, 8]:

1. An initial state space 𝕏_*J*_ is set, and the initial distribution along this space is extracted from *P* (0);
2. The probabilities *P* (*t*) and the FSP approximation error *z*(*t*) are integrated forward in time using Eq. 9. If *z*(*t*) exceeds the maximum error threshold function *ϵ*(*t*) at any time, the integration is paused, and the size of the truncated CME state space, 𝕏_*J*_, is expanded by adding states from *z*(*t*) - thus shrinking *z*(*t*) and lowering the total error - before continuing the integration;
3. The process is repeated until the final time, *t*_*N*_, at which point the approximation guarantees that the FSP solution is below the error threshold at all times.

In practice, the SSIT uses multiple absorbing sinks [**?**], where each sink corresponds to the set of all states that violate a defined polynomial inequality:

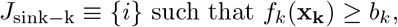

the FSP region is defined as

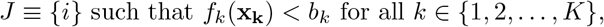

and the FSP approximation becomes:

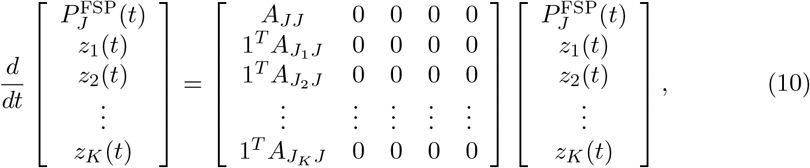

By keeping track of how much probability mass exits through each sink at each time step, the SSIT can automatically adjust the polynomial bounds *b*_*k*_ to recover that probability mass in that and subsequent time steps.

Box 12 and `example_2_SolveSSITModels_FSP.m` show examples for computing FSP solutions in the SSIT.

##### Example Box 12: Directly Solve the CME with FSP

Below is an example for computing FSP solutions in the SSIT, using our 4-state STL1 model (Box 1), with results shown in Fig. B12.

~~~
% Ensure the solution scheme is set to FSP (default):
STL1_4state.solutionScheme = ‘FSP’;
% Set FSP 1-norm error tolerance:
STL1_4state.fspOptions.fspTol = 1e-4;
% Guess initial bounds on FSP StateSpace:
STL1_4state.fspOptions.bounds = [1,1,1,1,200];
% Approximate the steady state for the initial distribution:
STL1_4state.fspOptions.initApproxSS = true;
STL1_4state.tSpan = [0:5:60];
STL1_4state.initialTime = 0;
% Compile and store the given reaction propensities:
STL1_4state = STL1_4state.formPropensitiesGeneral(‘STL1_4state’);
% Solve Model:
[∼,∼,STL1_4state] = STL1_4state.solve;
~~~

**Figure B12.**
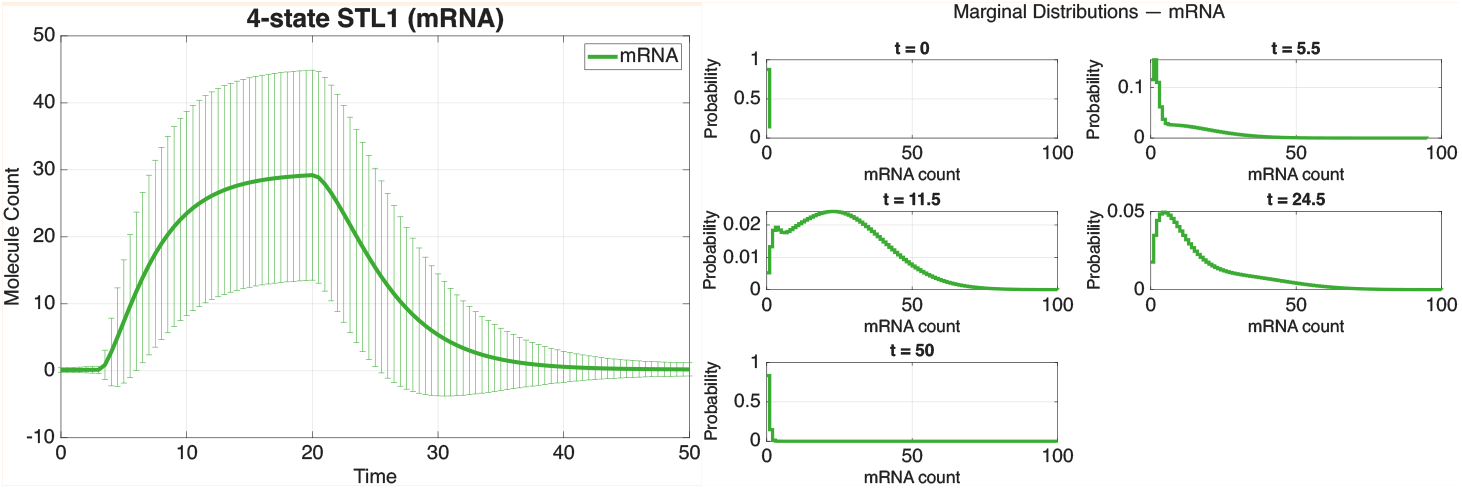
FSP means and standard deviations (top) and marginal distributions at select time points (bottom) for the 4-state STL1 model from Box 1.

#### 2.2.6 Escape (First Passage) and Waiting Time Calculations

In many applications, we are interested not only in the instantaneous probability distribution on states, but also in the *escape* behavior of the process [**?**]. For a continuous-time Markov chain *X*(*t*) with state space *S* and a subset *J* ⊂ *S*, the (random) escape time from *J* is defined as

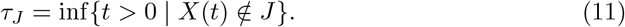

The corresponding cumulative escape probability (or escape-time distribution function) is

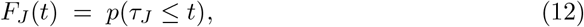

which quantifies the fraction of trajectories that have left the subset *J* by time *t*.

In the FSP formulation, the CME is approximated by a finite subset (𝕏_*J*_ ⊂ 𝕏) and all states in the complement (𝕏_*J*_*′*) are replaced by an absorbing sink variable *z*(*t*), as in Eq. 9. 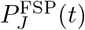 is the probability vector over the truncated state set 𝕏_*J*_, *A*_*JJ*_ is the restriction of the generator to 𝕏_*J*_, and **1** is the vector of ones of appropriate length. By construction,

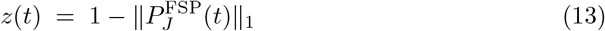

is exactly the cumulative probability of having left 𝕏_*J*_ by time *t*, and therefore

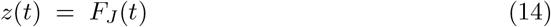

for the full CME.

To separate the desired escape behavior from numerical truncation error, it is useful to further decompose the state space into three disjoint and exhaustive subsets,

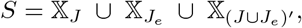

where *X*_*J*_ is the set of non-absorbing states of the FSP projection, 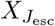 is the collection of all states that satisfy the intended escape criteria, and 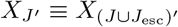 contains all remaining states that will be combined into an auxiliary absorbing “error sink”. Introducing two variables for the escape and error sinks, we can write the modified FSP as:

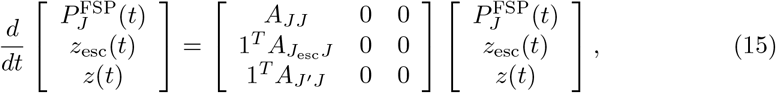

where *A*_*JJ*_ contains information about all reactions that are contained within 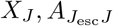 contains the reactions that transition 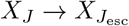, and *A*_*J*_*′*_*J*_ contains the reactions that transition *X*_*J*_ → *X*_*J*_*′*. With this reformulation, *z*_esc_ is now a strict *lower bound* estimate on the cumulative escape probability, 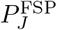 is a strict lower-bound FSP estimate for the probability mass that has not yet escaped (i.e., *P*_*J*_), and *z*(*t*) is the total error of the FSP approximation:

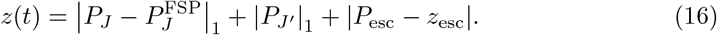

Classical quantities such as the mean escape time from an initial state *i* to an absorbing target set *J*_1_ can be recovered from the escape-time distribution via

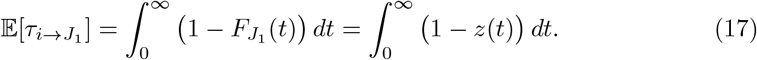

Box 13 and `example_5_SolveSSITModels_EscapeTimes.m` show examples for computing escape time solutions.

##### Example Box 13: Computing First Passage Time Probabilities with the FSP

Below, we provide an example for computing escape time solutions that solves for the time until the ‘mRNA’ (from our 4-state STL1 model, Box 1) count reaches 100. It is worth noting that although the 4-state STL1 model is non-ergodic/non-stationary in the time scale we consider, the calculation of escape time distributions from within transient responses to Hog1 input is both still useful to consider and appropriate, as they are defined in terms of when trajectories leave a chosen set of states.

The cumulative distribution function and probability distribution functions are shown in Fig. 13.

~~~
% Set the times at which distributions will be computed:
STL1_4state.tSpan = linspace(0,100,200);
% Solve for time for mRNA to reach 100:
STL1_4state.fspOptions.escapeSinks.f = {‘mRNA’};
STL1_4state.fspOptions.escapeSinks.b = 100;
[∼,∼,STL1_4state] = STL1_4state.solve;
~~~

**Figure B13.**
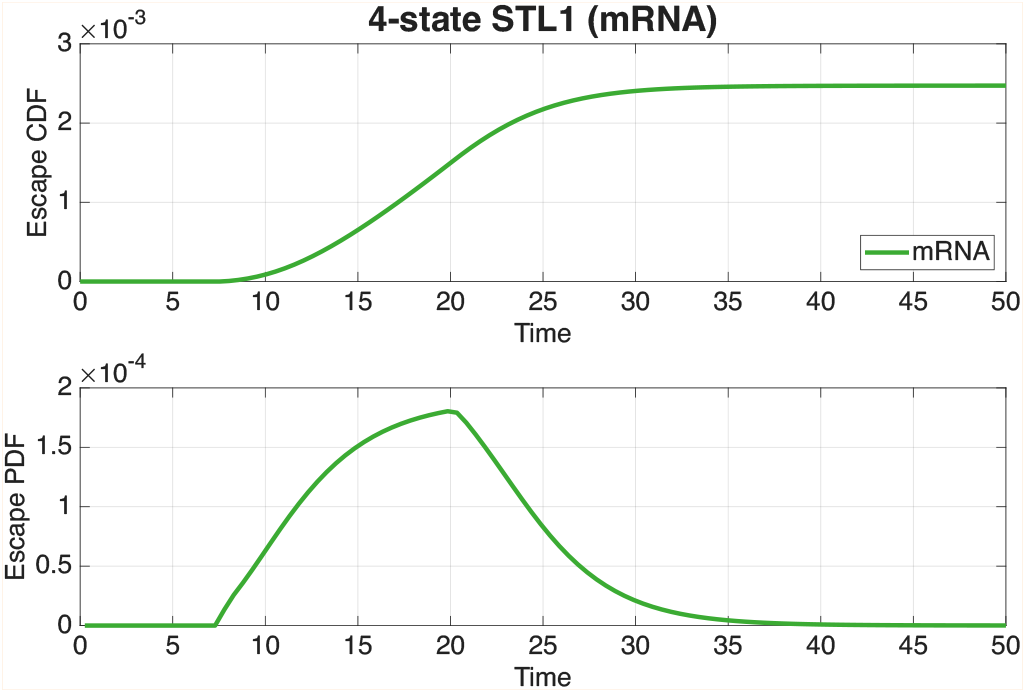
Escape CDF and PDF for time to *mRNA* = 100 for the 4-state STL1 model.

#### 2.2.7 Sensitivity Analysis

Parameter sensitivity analysis provides quantified insights on how the model parameters affect outputs, and consequently, which parameters have greater effects than others, the extent of parameter identifiability, and which experimental conditions may yield the most information about underlying mechanisms [**?, ?**]. As there are different methods for computing derivatives, there are different approaches to computing the sensitivities of a model output with respect to its parameters. The SSIT focuses on the sensitivity of the CME solutions to parameter changes:

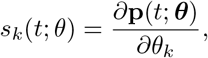

where **p**(*t*; ***θ***) denotes the probability vector (or other model output) at time *t*. To estimate this quantity, the SSIT provides two approaches: finite difference, and forward integration as follows:

##### Finite differences

A basic approach is to approximate the sensitivity *s*_*i*_(*t*; *θ*) using small perturbations *h* along the unit basis vector *e*_*i*_:

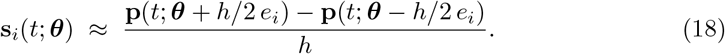

For stochastic reaction networks, variance-reduced couplings (e.g., common random numbers, pathwise/likelihood-ratio hybrids) are often used when **P** is estimated by simulation; for FSP, (18) is computed by re-solving on the same truncated state space.

##### Forward (FSP) sensitivities

The forward integration approach uses a rearangement of the time derivative of the sensitivity function:

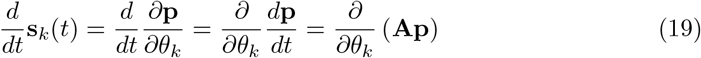

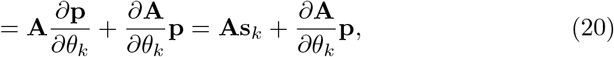

which can be solved simultaneously with **p**(*t*) and using the initial condition 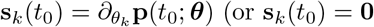 if the initial probability distribution is assumed not to be a function of the parameters).

FSP escape/absorbing states obey an analogous sensitivity balance [**?**]. Computation time for the forward sensitivity and the finite difference approach scales linearly with the number of parameters [**?**].

These sensitivities can then be used to calculate gradients of any distribution-based objective: e.g., statistical moments, prot=babilities of rare events, log-likelihoods for maximum likelihood estimation (Section 2.3.3) and Bayesian inference (Section 2.3.4), distance metrics, or Fisher Information Matrix (Section 2.2.8) entries.

Box 14 and `example_6_SensitivityAnalysis.m` show how to compute sensitivity solutions in the SSIT.

###### Example Box 14: Performing FSP-Based Sensitivity Analysis

Below, we show an example usage of the SSIT’s sensitivity solver for our 4-state STL1 model from Box 1 and show solution plots for parameter sensitivities by mRNA count at *t* = 25 in Fig. B14.

~~~
% Set solution scheme to FSP sensitivity:
STL1_4state.solutionScheme = ‘fspSens’;
% Solve the sensitivity problem:
[∼,∼,STL1_4state] = STL1_4state.solve;
~~~

**Figure B14.**
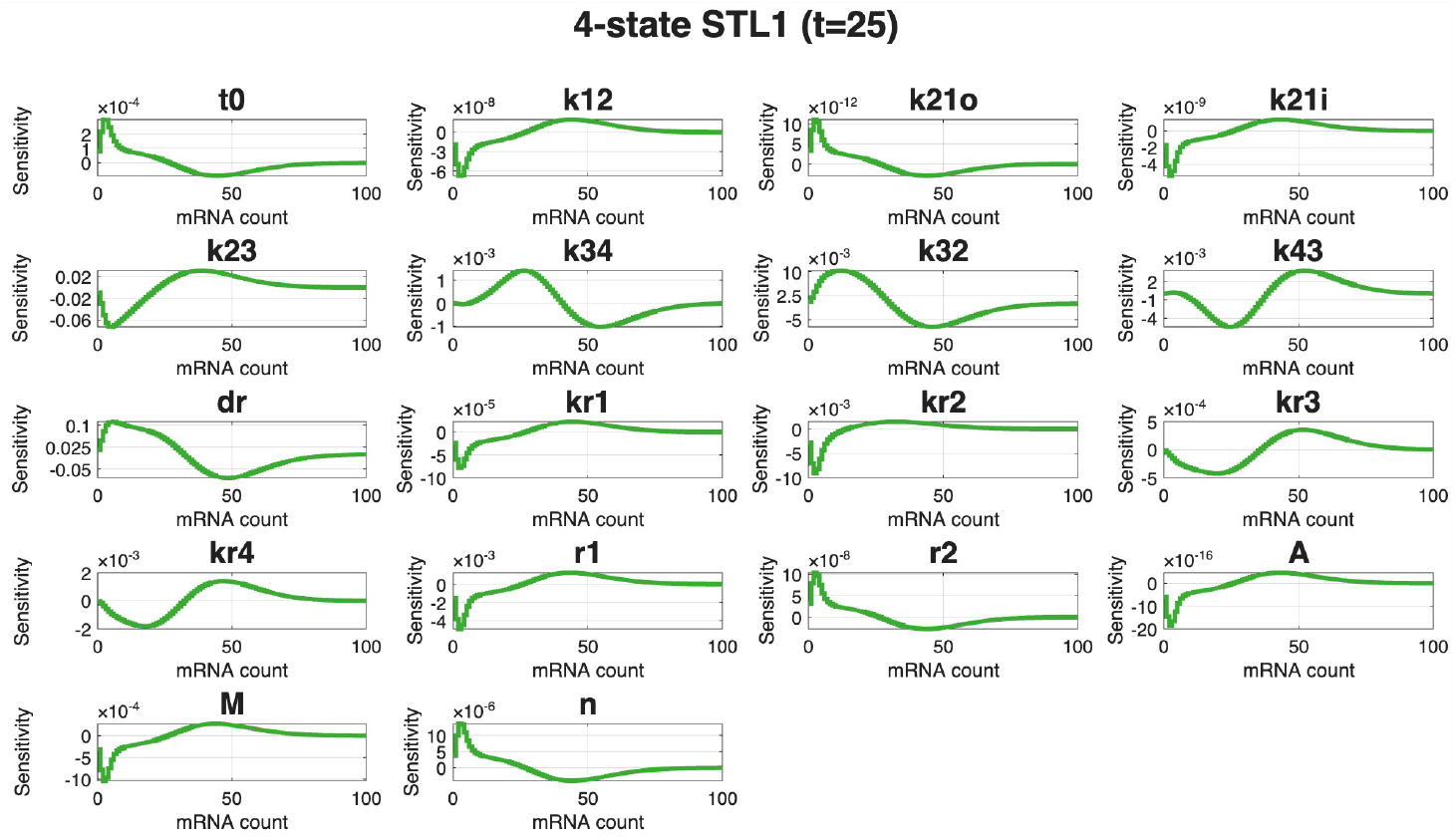
Sensitivities at *t* = 25 of each 4-state STL1 model parameter by mRNA count from the 0.2 nM dataset [33].

#### 2.2.8 Fisher Information Matrix (FIM)

The Fisher Information Matrix (FIM) is a statistical tool that quantifies how much information an observable random variable contains about unknown model parameters ***θ*** [**?**]. In other words, the FIM characterizes how tightly an experiment can, in principle, constrain ***θ*** by measuring the local curvature (precision) of the likelihood around a parameter value [**?**]. When a maximum likelihood estimate (MLE) 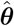 has been computed from data, the FIM determines the asymptotic covariance of 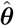. In a Bayesian setting, under standard regularity conditions and a sufficiently smooth/weakly informative prior, the posterior distribution becomes approximately Gaussian for large samples with covariance governed by the FIM [**?**]. Thus, the FIM is a natural tool for assessing parameter identifiability, for diagnosing stiffness in parameter directions, and for making decisions about sequential experiments [**?**].

The utility of an FSP-based FIM approach for single-cell experiments was demonstrated by Fox *et al*. [**?**]. They used the FSP algorithm to compute the FIM by solving the Chemical Master Equation (CME) for constitutive gene expression, including in cases where expression was strongly non-Gaussian. Unlike previous approaches that approximate population distributions as Gaussian [**?, ?**], which may be suitable only for large-population limits, the Fox *et al*. FSP-FIM framework works directly with the full discrete distribution and therefore captures the inherent stochasticity of small-copy-number systems.

Intuitively, the tighter a (locally) Gaussian distribution for 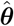, the more information the data provide about the underlying model parameters ***θ***. For sufficiently large sample size *N*_*c*_, the covariance 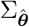 of an unbiased MLE 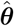 satisfies

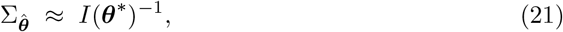

where *I*(***θ***^∗^) is the FIM evaluated at the true parameter ***θ***^∗^. The matrix *I*(***θ***^∗^)^−1^ sets the Cramér–Rao lower bound (CRB) on 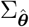 [**?**]; moreover,

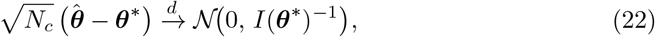

as *N*_*c*_ → ∞ [**?**]. Diagonalizing the FIM as *I*(***θ***^∗^) = *Q*Λ*Q*^⊤^, the eigenvectors (columns of *Q*) define principal directions in parameter space, and the eigenvalues in Λ determine the scale of uncertainty along those directions: large eigenvalues correspond to well-constrained combinations of parameters, while small eigenvalues indicate poorly constrained directions.

To connect with the likelihood, define the *score* (or *informant*)

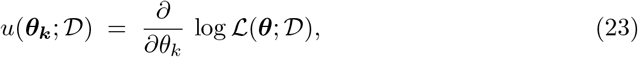

where ℒ (***θ***; 𝒟) is the likelihood for data 𝒟, (see Section 2.3.2 for likelihood calculation). The FIM is then the covariance (equivalently, the second central moment) of the score:

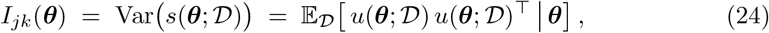

which, iterating over parameter combinations (i.e., in component form), gives

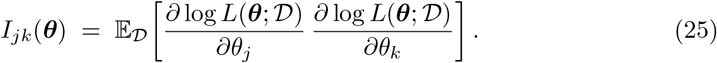

Under standard regularity assumptions, this also equals the negative expected Hessian of the log-likelihood [**?**].

For the FSP-based CME model (Section 2.2.5), we consider independent single-cell measurements drawn from the probability distribution *p*(*x, t*_*m*_; ***θ***) at one or more observation times *t*_*m*_. For a single measurement of a cell state *X* with probability mass function *p*(*x*; ***θ***), the contribution to the score is

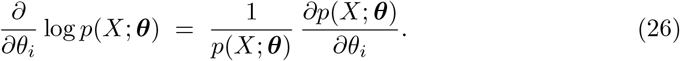

Using the sensitivities defined in Section 2.2.7, 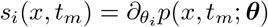 the FIM for *N*_*c*_ independent cells observed at a fixed time *t*_*m*_ can be written as

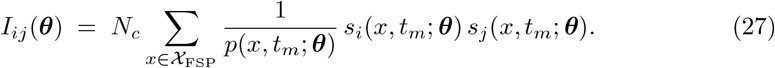

When multiple observation times and experimental conditions are used, their contributions are additive: one sums expressions of the form (27) over all times and conditions, with appropriate cell counts for each. Because (27) is derived directly from the FSP probabilities and their sensitivities, it inherits the FSP’s ability to represent non-Gaussian, multimodal, or highly skewed distributions, making it well-suited for optimal design and identifiability analysis in single-cell experiments [**?**].

Box 15 and `example_7_FIM.m` give examples for computing the FIM. Ellipses illustrating the *θ*-*θ* relationships for a subset of parameters are provided in the Supporting Information.

##### Example Box 15: Computing the Fisher Information Matrix

Below is an example for computing the FIM sub matrix for the free parameters in our 4-state STL1 model (Box 1), as the five Hog1 signal parameters (‘r1’, ‘r2’, ‘A’, ‘M’, and ‘n’) are experimentally known and can thus be fixed. In Fig. 15, we show heat maps, per-*θ* information, and eigenvalue spectra for both the full FIM and free FIM results.

~~~
% Compute the FIM sub matrix for free parameters:
STL1_4state_fimResults = STL1_4state_FIM.computeFIM([],’log’,[],freePars);
% Generate a count of measured cells:
cellCounts = 1000*ones(size(STL1_4state_FIM.tSpan));
% Evaluate the provided experiment design (in “cellCounts”)
% and produce an array of FIMs (one for each parameter set):
[STL1_4state_fimTotal,STL1_4state_mleCovEstimate,STL1_4state_fimMetrics] = …
     STL1_4state_FIM.evaluateExperiment(STL1_4state_fimResults,cellCounts)
~~~

**Figure B15.**
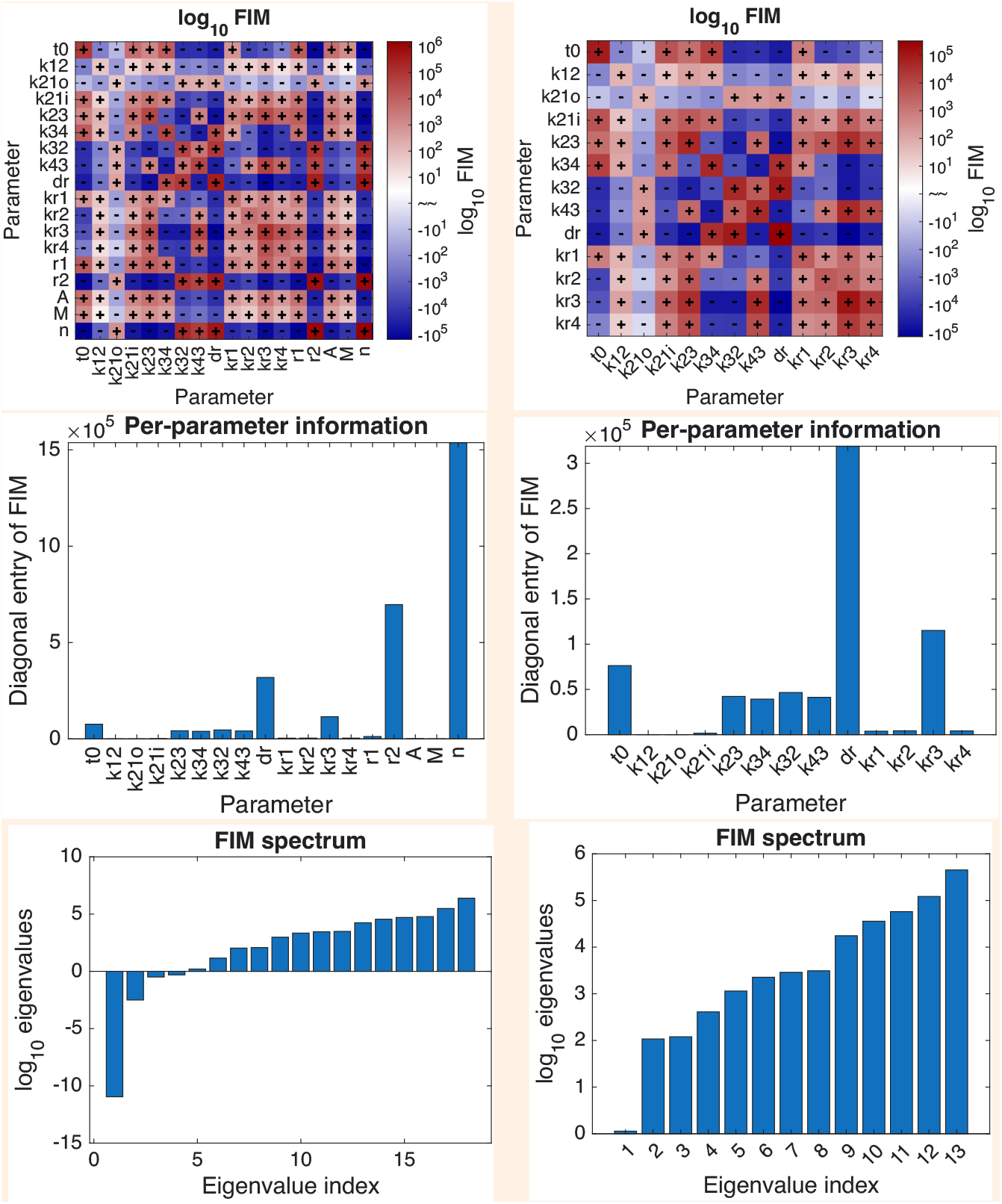
(Top) *θ*-heat maps; (middle) per-*θ* information; and (bottom) FIM spectra for (left) full FIM results and (right) results using only free parameters.

#### 2.2.9 FIM Optimality Criteria and Experiment Design

The Fisher Information Matrix *I*(***θ***; *ξ*) for a given experimental design *ξ* (e.g., choice of input signals, measurement times, doses, or imaging settings) quantifies how much information that design can provide about the model parameters ***θ***. [**?, ?, ?**] In practice, we use the FIM to compare candidate designs and to select experiments that are expected to yield the most informative data, either with respect to all parameters or to a subset of parameters of particular interest [**?, ?**, 14].

Several scalar *optimality criteria* compress the FIM into a single design score, each emphasizing different aspects of parameter uncertainty [**?, ?**]. The SSIT offers:

- **E-optimality** (default) - maximizes the smallest eigenvalue of *I*(***θ***; *ξ*), ensuring that no direction in parameter space is too poorly estimated (which can be advantageous or disadvantageous depending on the presence of weakly identifiable parameters).
- **D-optimality** - maximizes the expected determinant, det *I*(***θ***; *ξ*) - equivalently, minimizes det *I*(***θ***; *ξ*)^−1^, the volume of the confidence ellipsoid in parameter space and thus reduces overall uncertainty in all linear combinations of parameters.
- **D**_**s**_**-optimality** - maximizes the determinant of a sub-matrix of the FIM to reduce uncertainty for a particular subset of parameters. This may be especially useful for hierarchical models; the SSIT provides D_s_-style criteria for subsets of or grouped parameters.
- **Trace-based optimality** - maximizes the trace of the FIM, tr(*I*(***θ***; *ξ*)). Trace criteria is similar to **A-optimality**, which minimizes tr *I*(***θ***; *ξ*)^−1^), the average variance across parameters. A-optimality is often viewed as a more conservative variant of D-optimality because it tends to avoid extreme experimental conditions - e.g., very large doses or impractical imaging regimes.

Other FIM-based criteria target different goals. **G-optimality** minimizes the worst-case prediction variance over a specified design or prediction space, prioritizing uniform predictive accuracy rather than parameter precision alone. **T-optimality** and related model-discrimination criteria focus on maximizing differences in model predictions, thereby favoring designs that most clearly distinguish between competing mechanistic hypotheses [**?, ?**].

Within the FSP-FIM framework [**?**], each of these criteria can be evaluated by combining the distribution-level sensitivities **S**_*i*_(**x**, *t*; ***θ***) with the experiment-specific cell counts and observation times, and then optimizing over *ξ* to guide sequential experiment design [**?**]. The SSIT allows for the simultaneous calculation of the FIM at arbitrary sets of parameter values allowing the user to sample of parameter uncertainty to find and optimize the average information within an experiment for unknown parameters [34].

In Box 16, we optimize the sampling of 13k cells across 13 time points, from 0 to 60 minutes, using five different criteria (the full example code is provided in `example_7b_FIM ExperimentDesign.m`). This difference in the optimal experiment for the different optimality criteria clearly illustrates the importance for one to understand and choose the optimality criteria that is best suited to a specific research question.

##### Example Box 16: Design Optimal Experiments Using the FIM

Here is an example for computing the optimal number of cells using the FIM.

~~~
% Compute FIM results:
fimResults = STL1_4state_design.computeFIM([],’log’,[]);
% Get the number of cells from loaded experimental data using ‘nCells’:
cellCounts_data = STL1_4state_design.dataSet.nCells * …
                   ones(size(STL1_4state_design.tSpan));
%% Experiment Design: Find the FIM-based designs for a total cells
% Compute the optimal number of cells from the FIM results using different
% design criteria: ‘Trace’ maximizes the trace of the FIM;
% ‘D-cov’ minimizes the expected determinant of MLE covariance;
% ‘E-opt’ maximizes the smallest e.val of the FIM; and
% ‘D-opt-sub[$<i_1>,<i_2>$,…]’ maximizes the determinant of the FIM for
% the specified indices. The latter is shown for different parameter
% combinations, where D-opt-sub[9:13]’ are the mRNA-specific parameters
% ‘dr’ and ‘kr1’,’kr2’,’kr3’, and ‘kr4’ (degradation and transcription
% reactions). All other parameters are assumed to be known and fixed. nCol = sum(cellCounts_data);
nTotal = nCol(1);
nCellsOpt_Dcov = STL1_4state_design.optimizeCellCounts(fimResults,nTotal,’D-cov’);
nCellsOpt_Trace = STL1_4state_design.optimizeCellCounts(fimResults,nTotal,’Trace’);
nCellsOpt_Doptsub = STL1_4state_design.optimizeCellCounts(fimResults,nTotal,…
                                                               ‘D-opt-sub[1:8]’);
nCellsOpt_DoptsubR = …
      STL1_4state_design.optimizeCellCounts(fimResults,nTotal,’D-opt-sub[9:13]’);
nCellsOpt_DoptsubI = …
      STL1_4state_design.optimizeCellCounts(fimResults,nTotal,’D-opt-sub[14:18]’);
~~~

**Figure B16:**
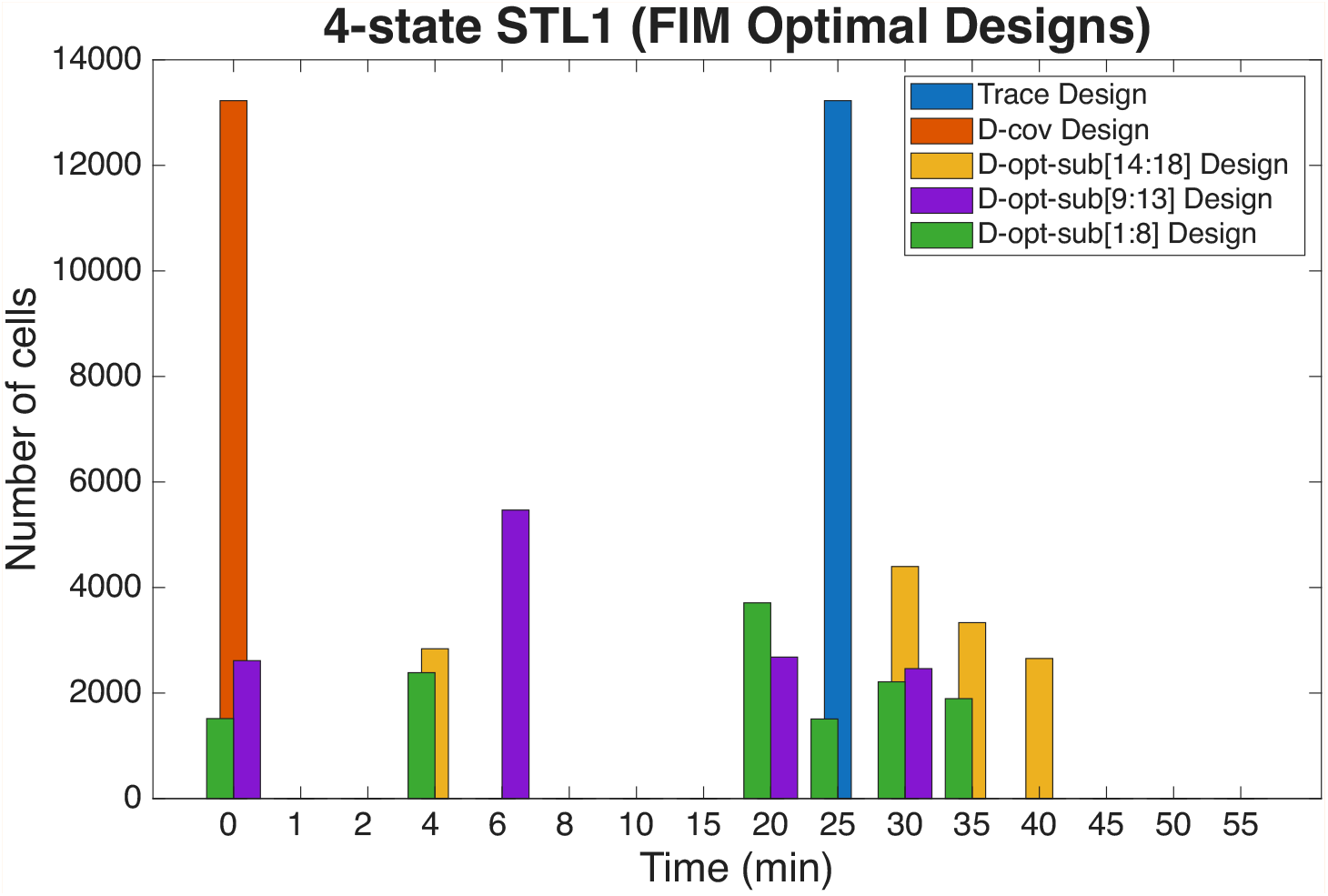
The optimal number of cells per time point based on different FIM criteria: ‘Trace’ is trace-based optimality and maximizes the trace of the FIM; ‘D-cov’ is a generalized D-optimality and minimizes the expected determinant of MLE covariance; ‘E-opt’ is E-optimality and maximizes the smallest e.val of the FIM; ‘D-opt-sub[*< i*_1_ *>, < i*_2_ *>*,…]’ is a type of D-optimality that maximizes the determinant of the FIM for the specified indices. The latter is shown for different parameter combinations, where ‘D-opt-sub[14:18]’ are the Hog1 input parameters; ‘D-opt-sub[9:13]’ are the mRNA-specific parameters ‘dr’ and ‘kr1’,’kr2’,’kr3’, and ‘kr4’ (degradation and transcription reactions); and ‘D-opt-sub[1:8]’ are the bursting gene parameters for the four gene states of the model from Box 1.

### 2.3 The SSIT provides integrated capabilities to load, process, and fit experimental data

In this section, we discuss and demonstrate how to load experimental or saved simulated data, associate it with a model in the SSIT, and fit the model parameters to the data using experimental STL1 data [33] and our 4-state STL1 model (Box 1).

#### 2.3.1 Data Loading and Handling

Although the SSIT is general and can be used for the modeling of any discrete stochastic system, for the sake of simplicity, we focus on single-cell experiments to illustrate the usage and underlying methodology of the SSIT and its associated packages. This choice is motivated by the rapidly growing body of quantitative experiments that capture the *in situ* heterogeneity and intricacies of molecular dynamics at the most fundamental level of biology [**?, ?**, 8].

The Chemical Master Equation has been successfully applied to study numerous single-cell processes, e.g., in the elucidation of transcriptional and translational bursting and the contribution of each reaction’s effects on RNA or protein levels within a cell population [33], to infer the kinetic parameters driving gene regulatory networks from time-series data [**?**], and to describe phenotypic variability between single cells [**?**].

A particularly powerful approach to capturing intracellular process data, and therefore the dynamic processes occurring within cells, is to use fluorescence microscopy [**?, ?**]. Single-molecule fluorescent imaging involves the attachment of probes [**?, ?, ?**] to individual molecules and enables their tracking and localization within individual cells. Single-molecule fluorescence in-situ hybridization (smFISH) is a technique for measuring gene expression and localizing RNA *in situ* by attaching fluorescent complementary DNA oligos to individual mRNA molecules [**?, ?**]. The cells are then imaged under an epi-fluorescence microscope, where each DNA probe on its individual mRNA is shown as a single bright spot. Each spot can then be counted, and the total number of spots is treated as a direct measurement of the number of mRNA molecules that were transcribed in the cell [**?**]. However, counting individual spots can be tricky due to the 3D nature of cells, not solely to determine localization but also to avoid miscounts due to overlapping spots.

Single-cell experiments have transformed biological research by enabling the characterization of cellular heterogeneity at unprecedented resolution. Unlike traditional bulk assays that report only population-averaged measurements, single-cell technologies quantify molecular features such as gene expression, protein abundance, and epigenetic modifications in individual cells. Advances in single-cell RNA sequencing (scRNA-seq) now make it possible to capture the transcriptomic landscape by profiling thousands to millions of cells in a single experiment, uncovering rare cell populations, delineating developmental trajectories, and revealing cell-specific responses to perturbations [**?, ?, ?**]. These approaches have also expanded beyond transcriptomics to include chromatin accessibility [**?**], DNA methylation [**?**], and protein levels [**?**], providing a more integrated view of cellular state and function.

However, single-cell data pose unique computational and statistical challenges, including technical noise, dropout events, and high dimensionality. Addressing these issues requires specialized modeling approaches, such as probabilistic frameworks, dimensionality reduction methods, and tools for data integration and imputation. The SSIT is applicable to many of these settings: even continuous measurements can be discretized, and the associated measurement error can be explicitly modeled and quantified using probabilistic distortion operators (PDOs) and related error-handling functions [34], (see Section 2.4.3).

##### File types

The SSIT reads count data, or other numeric data that may be interpreted as count data by the SSIT, such as total or average fluorescence intensities measures whose resulting distorted probability distributions can be transformed by the use of the SSIT’s *probabilistic distortion operators* (PDOs). Data files that can be read as a table or a cell by MATLAB’s ‘readtable’ and ‘iscell’ functions can be read by the SSIT, including: text files (.txt and.csv) and Microsoft Excel files (.xls),

##### Loading Data

The ‘loadData’ function is used to load data into the SSIT. The SSIT has built-in filtering capabilities, called simply by specifying which strings should be searched for and which values should be used to filter. In this paper, we use measurements of nascent STL1 mRNA expressed as part of the mitogen-activated protein kinase stress response in yeast cells [**?**, 33] and from scRNA-seq data composed of 151 genes in breast cancer cells responding to a treatment of dexamethasone [29] in demonstratation of how to create, solve, and fit models; perform statistical inference; account for extrinsic noise; and design a best next experiment in the SSIT. Box 17 and `example_8_LoadingandFittingData_DataLoading.m` illustrate how to load and filter data in one command.

###### Example Box 17: Load Data and Associate with Model

Below is the SSIT loadData usage to load STL1 mRNA count data from the osmotic shock experiments in Neuert *et al*. [33] and associate the data with the 4-state STL1 model (Box 1). The first input is the location and name of the data file, in this case a.csv file. The second input links the name(s) of the model species with the appropriate name(s) of the data file columns; in this case, our data consists of counts for the model species ‘mRNA’ which are provided in the column ‘RNA STL1 total TS3Full’. The third input is optional and may be used for filtering the data by conditions specified within the data file; in this case, only the data from the second replica of 0.2M NaCl concentration experiments are applied to the SSIT model named ‘STL1_4state data’.

~~~
% Note: Ensure the search path is correct on the local machine:
STL1_4state = STL1_4state.loadData(‘data/filtered_data_2M_NaCl_Step.csv’,…
                                     {‘mRNA’,’RNA_STL1_total_TS3Full’},…
                                     {‘Replica’,1;’Condition’,’0.2M_NaCl_Step’});
~~~

#### 2.3.2 Likelihood Calculation

Once data has been loaded into SSIT and associated with a model, the data 𝒟 can be used to confine the model parameters ***θ*** to values that most likely represent the underlying biological process which generated the data. Often this involves the calculation of a likelihood using an appropriate likelihood function ℒ (𝒟 | ***θ***), which provides a quantification of the probability of observing the data under particular values of model parameters [**?**]. In other words, the likelihood function ℒ (𝒟 | ***θ***) yields insight pertaining to the degree of support shown by the data *D* as a function of various combinations of parameter values ***θ***. The exact specification of the likelihood function depends on the type of data being studied, and the subsequent use of the likelihood once computed is dependent on the complexity (dimensionality, information content, etc.) of the data, the available computation power, and the researcher’s statisical philosophy.

Here, we provide an example of an appropriate likelihood function that would fit the type of data for which SSIT has been constructed. First, we describe observed molecular counts as 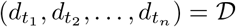 taken from single cell experimental data for a specific cell measured at time points *t* = [*t*_1_, *t*_2_, …, *t*_*n*_]. In our ‘Model’ example, these data are the numbers of mRNA from a particular cell at time points *t*. The likelihood function is:

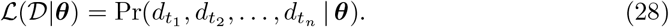

Note that in this formulation the data 𝒟 is fixed, and the likelihood is dependent only on the model parameters, ***θ***.

The likelihood function in Eq. 28 can be estimated directly from the system’s state probabilities, i.e., when (a) an exact Chemical Master Equation (CME) solution can be computed, or (b) the CME can be approximated, such as with Finite State Projection (FSP) as discussed in Section 2.2. [**?, ?, ?**]

For example, having obtained the probabilities 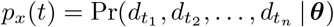 of CME states, then 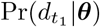 can be computed as the probability distribution at the first time point *t*_1_. Then probabilities at subsequent time points can be computed by continuing the solution forward in time and conditioning upon the previous solution. The second term is therefore 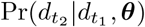 and the *n*^*th*^ term is 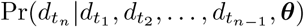, with the full iteration for calculating Eq. 28:

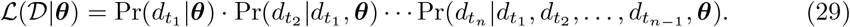

Generally, the procedure for computing the likelihoods of time-series data as in Eq. 29 is called filtering, and includes the Kalman filter [**?**] and HMM filter, the algorithm outlined above. This particular forward-time conditional description for calculating the likelihood based on state spaces is also known as the forward algorithm for state-space models [**?, ?, ?**].

Following Eq. 28 and 29 will yield the likelihood for data observed in a single cell, which does not provide much information about the biological system in question, especially when the system is based on highly stochastic processes and thus cells can exhibit highly variable dynamics. Fortunately, if we can appropriately assume in our current model that the processes taking place within each cell are independent to those of the other cells from which we are taking snapshot data, such as may be the case for smFISH and sequencing data, then it is straightforward to use Eq. 28/29 to compute the likelihood based on molecular count data taken from multiple independent cells at the same time. The independence between cells is a key feature because this means we can simply multiply the likelihoods for each molecular count at each time point from each single cell. Across *N*_*c*_ independent single-cell measurements, the likelihood can be reduced simply to:

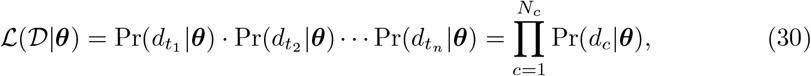

with *d*_*c*_ referring to the *c*^th^ cell, a formulation that has been widely used to fit stochastic gene-expression models to single-cell snapshot data [**?, ?**, 33]. For mathematical simplicity, we often use the log-likelihood form,

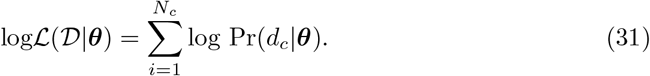

The likelihood function ℒ (𝒟 | ***θ***) is computed based on a particular set of model parameters ***θ***, the values of which must be iteratively sampled if not known *a priori* - which is always the case if the data is not simulated with known parameter values – and the likelihood recomputed for each sample of ***θ***. It might be obvious, then, that one method of determining the model parameter values that best fit the data is to find the set ***θ*** for which the likelihood of the data is the highest. In other words, the set of parameter values ***θ*** that maximizes the likelihood of the data, ℒ (𝒟 |***θ***).

#### 2.3.3 Maximum Likelihood Estimation

A maximum likelihood estimate (MLE) is the maximum value of the likelihood function, ℒ (𝒟 | ***θ***) [**?**]. An MLE is meant to reveal the set of parameter values ***θ*** that make the observed data 𝒟 most probable and is often represented with argmax [13], an argument that returns the maximum value of the function:

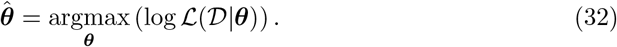

More details about log-likelihood maximization in the Supporting Information.

Box 18 and `example_9_LoadingandFittingData_MLE.m` demonstrate the use of maximum likelihood estimation to fit model parameters in the SSIT.

##### Example Box 18: Fit Model Parameters (MLE)

An example of fitting model parameters to loaded data using maximum likelihood estimation (MLE) is shown below. The results are shown in Fig. B18.

~~~
% Set fitOptions, with the maximum allowable number of iterations to fit:
fitOptions = optimset(‘Display’,’iter’,’MaxIter’,2000);
% Specify which parameters to fit (all but Hog1 input parameters, which are fixed):
fitPars = 13; STL1_4state.fittingOptions.modelVarsToFit = 1:fitPars;
% Store parameters for fitting:
STL1_4state_pars = cell2mat(STL1_4state.parameters(1:fitpars,2));
% Search to Find the MLE:
[STL1_4state_pars,STL1_4state_likelihood] = …
     STL1_4state.maximizeLikelihood(STL1_4state_pars,fitOptions);
% Update parameters:
for i=1:length(STL1_4state_pars)
     STL1_4state.parameters{i,2} = STL1_4state_pars(i);
end
~~~

**Figure B18:**
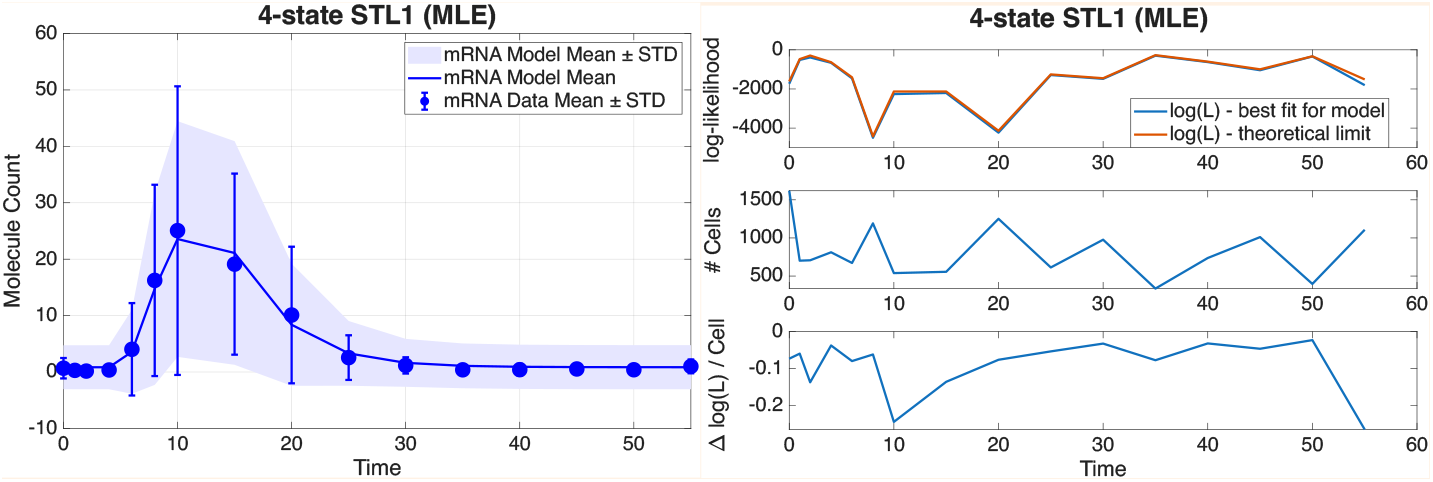
Trajectories of means and standard deviations for STL1 data [33] and the 4-state STL1 model fit to the data by MLE.

The higher the dimensionality of the model, the more parameter proposals need to be evaluated before the likelihood function will approach its global maximum. Similarly, the more parameters that must be estimated, the richer the data must be in information. When the systems being examined are stochastic in nature, there will inevitably be increased noise inherent in our data - in addition to any extraneous noise we add to it by our experimental, data collecting, and data processing choices – leading to *parameter uncertainty*. In other words, the parameter set yielded by 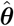 often does not actually represent the “true” parameters of the model or underlying the biological processes. We should therefore take precautions to account for uncertainty in our parameter estimates. We can use techniques such as Metropolis-Hastings to account for parameter uncertainty [20–22].

#### 2.3.4 Bayesian Inference

By its nature Bayesian inference provides measures of uncertainty surrounding parameter estimates and in the case of discrete variables (in our case, model parameters ***θ***) will yield probabilities over every hypothesis for how the data was generated, when computationally feasible. For continuous variables (***θ***), Bayesian inference yields posterior probabilities over the sampled space taken from a *posterior probability distribution*, or *posterior* [**?**, 17].

Bayesian inference is a method that takes a prior belief or knowledge Pr(***θ***) about the set of model parameters ***θ*** *a priori* to observing the data from our most recent experiment or simulations and updates that state of knowledge about ***θ*** based on the likelihood ℒ (𝒟|***θ***) of observing the current data 𝒟 given a set of ***θ*** values by computing the conditional probability Pr(***θ***|𝒟) *a posteriori* to observing some data, 𝒟, and normalizing by the probability of observing the data, Pr(𝒟), i.e.,

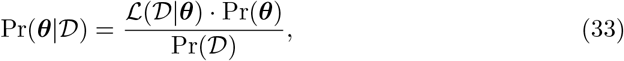

which is commonly referred to as Bayes’ theorem [23, 24].

A common method employed to sample the posterior probability distribution is Markov Chain Monte Carlo (MCMC), particularly the Metropolis-Hastings (MH) algorithm [20–22]. We elaborate on Bayesian inference, MCMC, and MH in the Supporting Information. As per Eq. 21, the inverse of the FIM, *I*(*θ*)^−1^, can be used to determine the covariance of the multivariate normal proposal distribution for the MHA, which can dramatically reduce correlations in parameter proposals and improve the sampling efficiency.

Boxes 19-20 and `example_10_LoadingandFittingData_MH.m` show an example of how to sample parameters using Metropolis-Hastings to produce the improved model fits and what to expect from the results. Example code for using *I*(*θ*)^−1^ as a proposal distribution for Metropolis-Hastings is also provided in the Supporting Information.

##### Example Box 19: Fit Model Parameters (Metropolis-Hastings): Code

~~~
% Specify how many model parameters will be fit:
fitpars = 13; STL1_4state_MH.fittingOptions.modelVarsToFit = 1:fitpars;
%% Specify Prior as log-normal distribution with wide uncertainty
% Prior log-mean:
mu_log10 = [0.5,2,5,3.5,-0.4,1,0.2,0.4,-0.5,-1.3,-0.1,2,0.5];
% Prior log-standard deviation:
sig_log10 = 2*ones(1,fitpars);
% Prior:
STL1_4state_MH.fittingOptions.logPrior = …
      @(x)-sum((log10(x)-mu_log10).^2./(2*sig_log10.^2));
% Create first parameter guess:
STL1_4state_MH_pars = [STL1_4state_MH.parameters{:,2}];
% Run Metropolis-Hastings (can be adjusted for better sampling
- generally aim for acceptance ratio around 0.3-0.4):
proposalWidthScale = 0.01;
MHOptions.proposalDistribution = @(x)x+proposalWidthScale*randn(size(x));
% Set MH runtime options (number of samples, burnin, thinning, etc.):
MHOptions.numberOfSamples = 2000; MHOptions.burnin = 500; MHOptions.thin = 2;
% Run Metropolis-Hastings:
[STL1_4state_MH_pars,∼,STL1_4state_MH_MHResults] = …
         STL1_4state_MH.maximizeLikelihood([], MHOptions, ‘MetropolisHastings’);
% Store MH parameters in model:
STL1_4state_MH.parameters([1:fitpars],2) = num2cell(STL1_4state_MH_pars);
~~~

##### Example Box 20: Fit Model Parameters (Metropolis-Hastings)

To sample parameter uncertainty and improve the fit of our 4-state STL1 model parameters (Box 1) to STL1 data [33], we iterate between computing maximum likelihood estimates and sampling by Metropolis-Hastings rejection scheme. Example code is provided in the following example block, and results are shown below in Fig. B20.

~~~
% Plot results:
STL1_4state_MH.plotMHResults(STL1_4state_MH_MHResults);
STL1_4state_MH.plotFits([], “all”, [], {‘linewidth’,2},…
       Title=‘4-state STL1’, YLabel=‘Molecule Count’,…
       LegendLocation=‘northeast’, LegendFontSize=18, ProbXLim = [0 80],…
       TimePoints=[0 8 10 15 30 55], TitleFontSize=24, AxisLabelSize=20);
~~~

**Figure B20:**
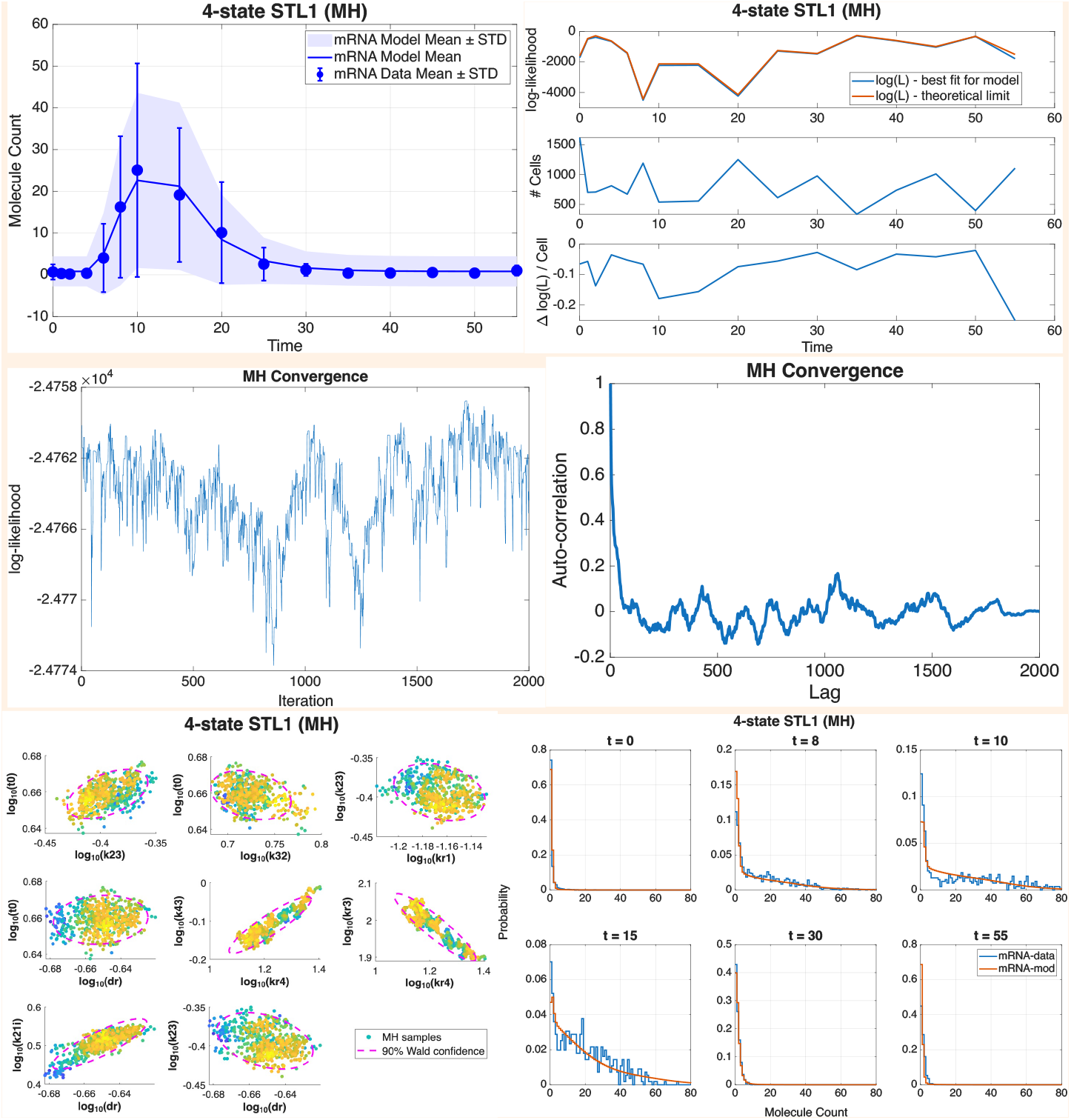
Trajectories of means and standard deviations for STL1 data; log-likelihoods (log(ℒ)) for best fit for model vs. theoretical limit; MH convergence by log(ℒ) and autolag; a subset of sampled model parameters; and probability distributions at select time points.

#### 2.3.5 Approximate Bayesian Computation

The SSIT also offers Approximate Bayesian Computation (ABC) for likelihood-free inference for cases in which a tractable likelihood is either unobtainable or infeasible to compute [26, 27]. Details about ABC implementation in the SSIT are provided in the Supporting Information and `example_SI_ABC.m`.

#### 2.3.6 Cross-Validation

The SSIT provides tools for cross-validation, where the model can be fit to different combinations of time points or replicas to determine how much parameters change and how well the model can predict some data when trained on other data. Additional details are included in the Supporting Information and `example_SI_CrossValidation.m`.

### 2.4 The SSIT provides advanced tools to create and analyze more complex combinations of models and datasets

In addition to sophisticated and time-varying propensity functions, the SSIT handles the following complexities: solving reduced models for increased computational efficiency (Section 2.4.1), including hybridized ODE-FSP solutions (Section 2.4.2); handling of probability distributions that have been distorted by data produced by particular choices in experimental design, measurement, and data processing (Section 2.4.3); and fitting multiple models and datasets (Section 2.4.4).

For Sections 2.4.3 and 2.4.4, we use an additional example dataset to showcase complex modeling in the SSIT, as these data benefit even more from probabilistic distortion handling (2.4.3) and multiple model-fitting (2.4.4) under the assumption of shared parameters. This second example data set consists of single-cell RNA-seq (scRNA-seq) measurements representing transcriptional responses of 151 different genes across five distinct time points following dexamethasone treatment in T47D A1-2 breast cancer cells [29]. Additional detail is provided in the Supporting Information).

#### 2.4.1 Additional Model Reduction Strategies

Model reduction techniques simplify complex mathematical models while preserving essential dynamical behavior. These strategies are particularly valuable in systems biology, control theory, chemical kinetics, and engineering, where full-scale models may be computationally expensive or analytically intractable due to high dimensionality or nonlinearities [**?, ?**]. The goal is to create a reduced model that is easier to simulate, analyze, and interpret, without sacrificing predictive accuracy in relevant contexts.

The FSP approximation provides an exact model reduction method to reduce the (potentially infinite) set of states of the Chemical Master Equation to a finite set while satisfying a strict error tolerance [**?, ?, ?**]. However, even after the FSP reduction many problems remain computationally burdensome or intractable. Fortunately, subsequent approximate reductions are also possible through application of additional projection and lifting operations. For most of these, one proposes a lower dimensional vector **q** such that **p** ≈ **Φq**. Substituting this approximation into 3, allows for one to formulate a reduced set of equations:

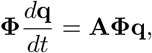

which yields

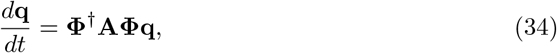

where **Φ**^*†*^ is the pseudo-inverse of **Φ**. This reduced differential equation can then be combined with the projected initial condition **q**(0) ≡ **Φ**^*†*^**p**(0) and solved using standard ODE integrators. One of the most powerful of these reduction strategies uses Krylov projections, where the projection operator is chosen as orthonormal basis spanning the space [**p, Ap, A**^2^**p**, …, **A**^*m*^**p**], which can be found efficiently using a modified implementation of Sidje’s Expokit [**?**]. For time invariant problems, the SSIT automatically uses Expokit both for the solution of the original FSP (Eq. 9) and for the solution of reduced linear ODEs (Eq. 34).

In addition to Expokit, the SSIT provide three categories of methods to define **Φ**, each using a different model reduction strategy: (1) time-scale separations that average over fast state dynamics, (2) local approximations that assume smooth probabilities functions nearby regions of state-space, and (3) data-driven learning of distribution shapes based on previous solutions.

##### Time-Scale Separation Methods

The “Quasi-Steady State Assumption” (QSSA) approach assumes that one or more species are “fast” compared to the others. These species are then assumed to equilibrate to their steady state probability distributions (conditioned on the slow species) allowing for the calculation of average reaction rates for the remaining slower species [**?**]. The SSIT model reduction scheme “Eigen Decomposition” (ED) employs a sparse eigenvalue decomposition to find the *N* largest (i.e., least negative) eigenvalues of the infinitesimal generator matrix **A** and then projects the master equation onto this lower dimensional space.

##### State Space Smoothing Methods

The “Linear State Lumping” (LNSL) and “Logarithmic State Lumping” (LGSL) model reduction strategies form coarse rectangular grids with linearly or logarithmically distributed points (within the current FSP bounds) and then combines these multiple state variables into fewer “lumped” states [**?**]. All states within each lumped region are assumed to have equal probabilities. The “Lin Lump QSSA” (LNQSSA) and “Log Lump QSSA” (LGQSSA) methods combine this lumping strategy with the time scale separation assumption to treat the states within each lump at quasi-steady equilibrium and then averages to find the rates of transitions out of the lumped states.

##### Data-Driven Learning Methods

The “Proper Orthogonal Decomposition” (POD) approach uses one or more previous full solutions of the CME at discrete time points (e.g., at similar values during a parameter search) and applies singular value decomposition (SVD) to identify a lower-dimensional subspace that best represents the dominant probability mass structures of those solutions in the least-squares sense. In other words **Φ**_PDO_ is an orthonormal matrix that spans the snapshot set [**p**(*t*_1_), **p**(*t*_2_), …]. The “Dynamic Mode Decomposition” (DMD) approach also uses a set of previous solutions, but instead of identifying modes that maximize variance in the snapshot distributions, it constructs a best-fit linear operator that maps one time snapshot to the next. Specifically, it seeks to determine an operator **Q** ≈ exp(**A**Δ*t*) such that **p**(*t*_*k*+1_) ≈ **Q p**(*t*_*k*_), and computes its eigen-decomposition. The resulting matrix **Φ**_DMD_ has columns that approximate eigenvectors of **Q**, so that **QΦ**_DMD_ ≈ **Φ**_DMD_**Λ**. Thus, span {**Φ**_DMD_} approximates an invariant subspace of the evolution operator, rather than merely the span of the snapshots themselves. The method “POD2” combines both of these approaches and defines **Φ** to span the combined set [**p**(*t*_1_), …, **p**(*t*_*N*_), **Ap**(*t*_1_), …, **Ap**(*t*_*N*_)].

We note that for the current implementations of the SSIT, the QSSA, ED, LNQSSA, LGQSSA, DMD, POD2 methods assume that the infinitesimal generator matrix **A** is constant, and are only appropriate for time invariant problems. Because the 4-state STL1 model (Box 1) example is time varying, the only appropriate methods are: POD, LNSL, and LGSL. The POD method is exemplified in the Supporting Information and `example_SI_ModelReduction.m`.

#### 2.4.2 Hybrid Solutions

The SSIT can assign upstream species to deterministic ODE solvers for efficiency, while maintaining full stochastic solutions for species of interest whose copy numbers are influenced by noise, in turn influencing the dynamics of the system (e.g., species with low copy numbers or switch-like species). Example code and results for computing hybrid ODE-FSP solutions for our 4-state STL1 model (Box 1), treating the four gene states deterministically and mRNA with full CME solutions, are provided in the Supporting Information and `example_SI_Hybrid.m`.

#### 2.4.3 Data Distortion Handling

Single-cell experiments are highly versatile and modifiable, broad in scope with nearly limitless possible designs but also varying greatly in cost with respect to both time and expense [**?**] and, accordingly, the degree to which they may yield information about the underlying models [**?**] or introduce errors [29, 34].

Most of the time, the results of single-cell experiments are assumed to have negligible measurement and data processing errors, and model development tends to assume ideal data as well [**?**]. However, while stochastic models are adopted with the idea of handling intrinsic noise, extrinsic noise can also have significant effects on inferred molecule counts and probability distributions [34]. Below are examples of the types measurement errors which may contribute to data noise and impact resulting analyses, model inference, and subsequent experimental design.

- Microscope resolution
- Photobleaching
- Optical filters
- Laser intensity
- Camera exposure time
- Imaging modality (z-stacking, max projection, etc.)
- Image processing (spot counting, intensity averaging, etc.)
- Delays due to drag/inducer diffusion or nuclear import

Capture inefficiencies such as allelic and transcript dropout in genotyping, single-cell DNA assays, and single-cell RNA sequencing are other examples of data distortions, yielding false zero or low counts for the dropout gene in the affected cell(s).

Vo *et al*. [34] laid the groundwork for quantifying and accounting for the effects of measurement and image processing errors on the accuracy and precision of model inference from single-cell fluorescence microscopy experimental data using conditional probability-based operators called *probabilistic distortion operators (PDOs)*. A PDO is simply a Markov kernel that transforms a distorted probability distribution, **p**_*Y*_ (*t*), computed using methods such as FSP (2.2.5) from data into which errors have been introduced by certain experiment design, measurement, or data processing choices, into a distribution more closely resembling the true distribution, **p**_*X*_ (*t*), or, more realistically, the distribution which would have been inferred from data containing fewer errors due to different design and/or processing choices.

If we considered a measurement distorted by noise taken at time *t* to be *y*(*t*) realized from the vector *Y* (*t*) which results from random distortions of the true process *X*(*t*) [34], then the relationship between **p**_*Y*_ (*t*) and **p**_*X*_ (*t*) can be described by the linear transformation

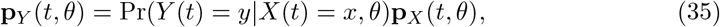

for each distorted measurement *y* and “true” measurement *x*, where Pr is either a probability mass or probability density depending on whether *Y* is discrete or continuous. The matrix **C**_*X*→*Y*_ containing all possible observations *y* indexing its rows is **C**_*X*→*Y*_ (*t*; *θ*):= Pr(*Y* (*t*) = *y* | *X*(*t*) = *x, θ*), which we call the PDO.

The steps of the data distortion handling (i.e., probability distribution transformation) process are as follows:

1. Collect molecular imaging data from chosen experiment.
2. Fit models to the data using the Finite State Projection (FSP) algorithm to efficiently solve Chemical Master Equations (CME) and yield the likelihood of the data and exact error.
3. Calibrate models to robustly handle measurement errors by computing and applying a distortion kernel [*C*_*ij*_].
4. Use Fisher information matrix (FIM) to determine how much information is obtainable from distorted experiments and approximate the sensitivity of the CME to changes in parameters.
5. Apply Bayesian experiment design methods to determine the most cost- and time-efficient experiments that can achieve sufficient information about model parameters and states of the single-cell systems under investigation.

To illustrate the application of a *probabilistic distortion operator* (PDO) on a real dataset, we treat the single-cell RNA-seq measurements from Hoffman *et al*. [29] as distorted observations of the underlying, “true” transcript counts. In particular, we interpret the observed unique molecular identifier (UMI) counts as the result of imperfect capture and detection of true mRNA molecules and model this measurement process using the “missing spot” Binomial PDO [34].

From Eq. 35, the observation model can be written as

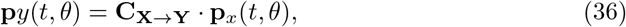

where **C**_**X**→**Y**_ is the PDO encoding the conditional probabilities of observing *y* UMIs given a true count *x*. Under the Binomial detection model, each column of **C**_**X**→**Y**_ is given by

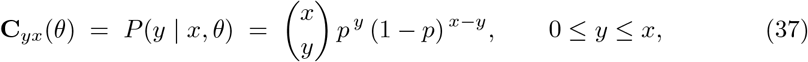

with *p* representing the effective capture probability (and 1 − *p* the corresponding missing-molecule rate). As before, a single row of the PDO may be denoted by **c***y* = **C***yx*(*θ*). The probability of observing a specific UMI count *y*_*i*_ at time *t* is then obtained by

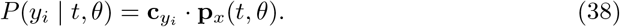

Droplet-based UMI scRNA-seq provides a sparse and incomplete sampling of cellular mRNA content; many transcripts present in the cell are not captured, reverse-transcribed, or sequenced, and low-copy transcripts are therefore frequently recorded as zeros. Hoffman *et al*. [29] reported 29, 448 UMIs per cell in their scRNA-seq measurements. A “typical” estimate of mRNAs/cell is 2 × 10^5^, which would be *p* ≈ 29, 448*/*200, 000. For our purposes, we adopt a conservative value of *p* = 0.05 (i.e., a 95% “dropout” or missing-molecule rate).

For the scRNA-seq data, we fix *p*, generate the PDO, and apply it to the model template prior to loading and fitting any data. This PDO is illustrated in Box 21 and `example_Seq1_BatchFitManyGenes.m`. As a case where one might one to calibrate a PDO - i.e., the hyperparameters that tune the conditional probability distribution - Box 21 also demonstrates the application of a Binomial PDO to the STL1 data [33] previously discussed. In the latter case, we use the nuclear mRNA count as the incomplete data with “missing” counts and the full-cell data as the “true” counts. This example, along with an affine Poisson PDO for transforming distributions from average intensity data, are provided in the Supporting Information and `example_11_ComplexModels_PDO.m`

##### Example Box 21: Use Probabilistic Distortion Operators

Below we show how to generate a Binomial PDO with an experimentally obtained failure probability 1-*p* for the scRNA-seq [29] data (and in Fig. 21 *left*):

%% Set PDO to Binomial for RNA:

~~~
Model_Template.pdoOptions.type = ‘Binomial’;
Model_Template.pdoOptions.unobservedSpecies = ‘onGene’;
Model_Template.pdoOptions.props.CaptureProbabilityS1 = 0;      % Gene State
                                                                % not measured
Model_Template.pdoOptions.props.CaptureProbabilityS2 = 0.05; % 95% drop out
[∼,Model_Template] = Model_Template.generatePDO();
~~~

An example of calibrating a Binomial PDO to data, in this case yeast STL1 [33] data, is shown below (Fig. 21 *right*):

~~~
STL1_4state_PDO_nuc = …
STL1_4state_PDO_nuc.calibratePDO(‘data/filtered_data_2M_NaCl_Step.csv’,…
          {‘mRNA’}, {‘RNA_STL1_total_TS3Full’}, {‘RNA_STL1_nuc_TS3Full’},…
           Binomial’, true, parGuess, {‘Replica’,1}, FontSize=24,…
           Title=“4-state STL1 (PDO: Nuclear mRNA)”, LegendLocation=“northwest”,…
           XLabel=“Total mRNA counts”, YLabel=“Nuclear mRNA counts”);
~~~

**Figure B21:**
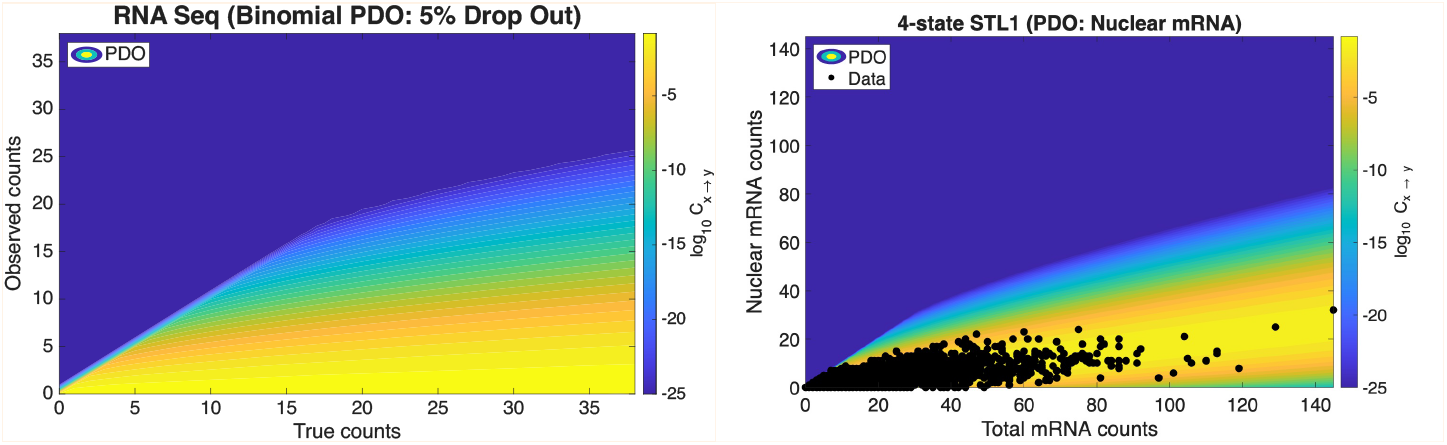
Conditional probability mass function contour maps of log_10_C_x→y_ for (left) the Binomial PDO with a conservative assumption of 95% dropout for scRNA-seq, and (right) a Binomial PDO calibrated to yeast STL1 data.

#### 2.4.4 Fitting Multiple Models and datasets with Shared Parameters

SSITMultiModel is a class included within the SSIT package that allows for combined parameter fitting across multiple datasets or models. Each dataset is associated with its own instance of an SSIT model but the group is assumed to share a subset of common parameters. Examples of situations that might benefit from SSITMultiModel include different experimental replicas expected to have batch parameter variations, analysis of the same regulatory system under different experimental conditions, or single-cell RNA seq (scRNA-seq) data consisting of different genes responding to the same upstream signals.

In Box 22 and `example_scRNAseq_1_BatchFitManyGenes.m`, we show an example demonstrating the use of SSITMultiModel to fit shared upstream input signal parameters but distinct bursting gene parameters for four different genes from scRNA-seq data [29]. In the Supporting Information and `example_15_ComplexModels_MultiModel.m`, we include examples using SSITMultiModel to fit different subsets of parameters of the 4-state STL1 model (Box 1) to two different experimental replicas [33].

##### Example Box 22: Multi Model

~~~
% Select which models to include in multimodel:
Models = {Model_DUSP1,Model_RUNX1,Model_BIRC3,Model_TSC22D3};
% Define how parameters are assigned to sub-models:
ParInds = {[1,2,3:9],[1,2,7*1+(3:9)],[1,2,7*2+(3:9)],[1,2,7*3+(3:9)]};
% Set a constraint on model parameters:
Constraint = @(x) -var(log10([x(3:9);x(7*1+(3:9));x(7*2+(3:9));x(7*3+(3:9))]));
% Create and initialize multimodel:
combinedModel = SSITMultiModel(Models, ParInds, Constraint);
combinedModel = combinedModel.initializeStateSpaces();
% Fit the multimodel:
fitOptions = optimset(‘Display’,’iter’,’MaxIter’,1000);
for i = 1:10
       [∼,∼,∼,combinedModel] = combinedModel.maximizeLikelihood([],fitOptions);
       save(‘seqModels/CombinedModel4Genes’,’combinedModel’);
end
~~~

**Figure B22:**
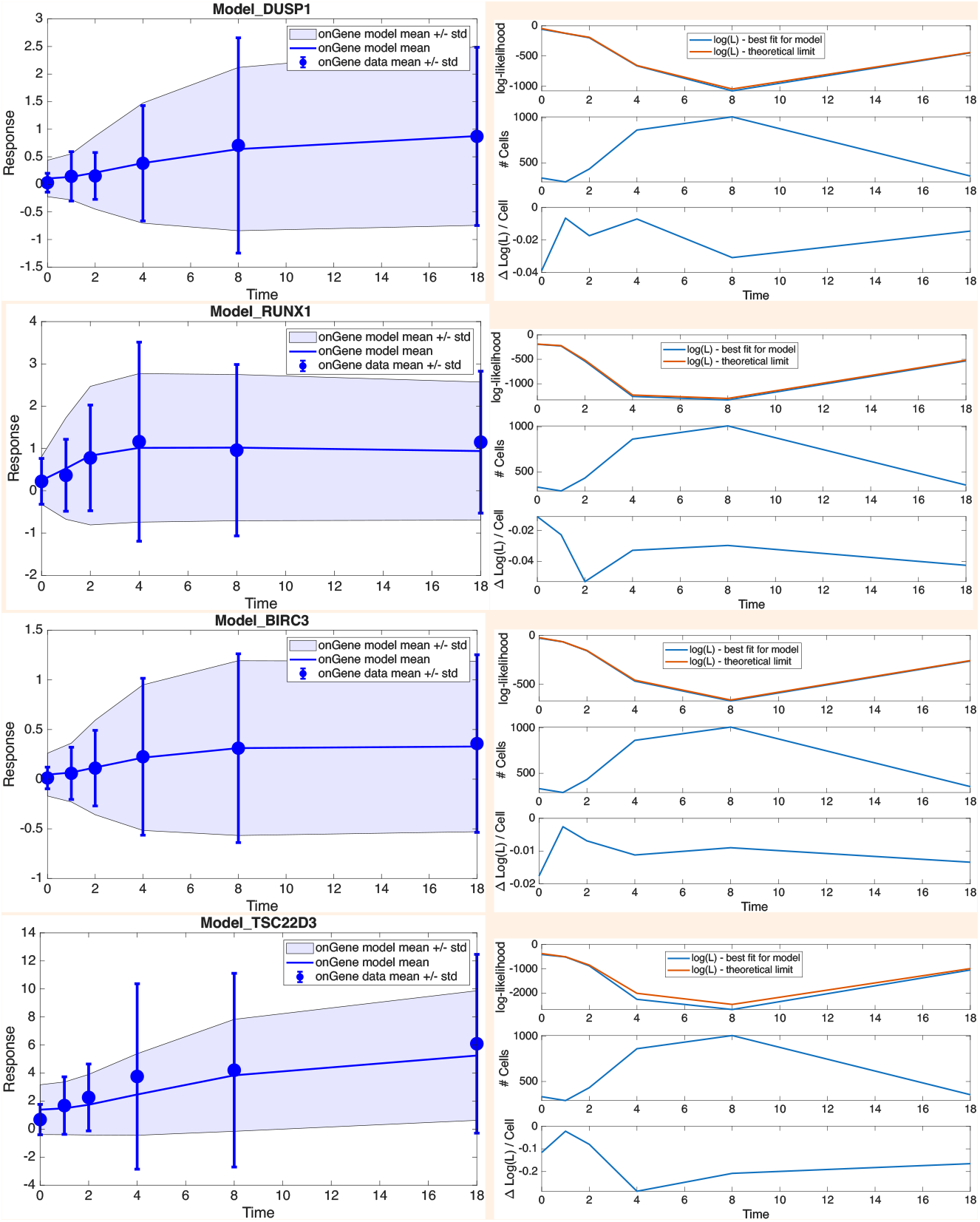
SSITMultiModel results from four different models, each representing a different gene (DUSP1, RUNX1, BIRC3, TSC22D3), fit to single-cell RNA sequencing data: (left) trajectories of means and standard deviations for each model and its associated data; (right) log-likelihoods, number of cells, and log-likelihoods/cell through time.

### 2.5 The SSIT can be extended to generate complex workflows using pipelines and cluster computing options

The SSIT can generate condensed pipelines that make it easy to launch long or repeated analyses in the background or on a high-performance computing (HPC) cluster. More computationally intensive processes that are handy to compute in the form of an SSIT pipeline include sampling using Metropolis-Hastings (2.3.4, Supporting Information), cross-validation (2.3.6), and parameter sweeps; these can be exported to a stand-alone command that runs non-interactively under the operating system’s job scheduler.

The SSIT provides a utility function `generateCommandLinePipeline`that serializes the model and pipeline configuration into a single MATLAB command-line call and returns it as a shell command string. A flexible set of input arguments are passed directly to the pipeline function, ensuring that the same configuration can be used interactively and in batch mode. Before execution, the command automatically adds the SSIT root directory and the temporary propensity-function directory (`tmpPropensityFunctions`) to the MATLAB path, so the job is self-contained and does not rely on a user’s interactive MATLAB session. When desired, the same command can be used to automatically launch the pipeline, either on a separate thread or on an open node on a shared research cluster. This is useful for launching long-running analyses from a workstation or login node without blocking the MATLAB session.

Box 23 and `example_scRNAseq_2_BatchFitManyGenes.m` provide example code to generate and execute pipelines for simultaneously fitting 151 models to the scRNA-seq data from within the MATLAB environment, from a local command line, or on a high-performance cluster. The Supporting Information and `example_scRNAseq_3_BatchFitManyGenes.m` show code for collecting the results of these fits to generate a figure (Fig. B23) to summarize and compare the fitted parameters and to identify different regulatory mechanisms across scRNA-seq genes.

#### Example Box 23: Generate and Run Pipelines (Local and HPC)

Below is the code used to generate a pipeline to fit the 151 most highly activated genes selected from the scRNA-seq dataset provided by Hoffman *et al*. [29]. Fig. 23 shows the mean fold change for comparison of the key parameters *K*_*ON*_ and *K*_*OF F*_ between all genes, each of which was fit as its own model for bursting RNA transcription. (Plotting code for Fig. 23 is shown in the Supporting Information.)

~~~
%% Call Pipeline to Fit scRNA-seq Models
% Specify pipeline to apply to model and arguments
% (“../../SSIT/src/exampleData/examplePipelines/fittingPipelineExample.m”)
Pipeline = ‘fittingPipelineExample’;
pipelineArgs.maxIter = 1000; pipelineArgs.display = ‘iter’;
pipelineArgs.makePlot = false; pipelineArgs.nRounds = 1;
%% Launch cluster jobs for all genes.
DataFileName = ‘data/Raw_DEX_UpRegulatedGenes_ForSSIT.csv’;
TAB = readtable(DataFileName);
geneNames = fields(TAB);
for iRound = 1:5
    for iGene = 2:length(geneNames)-4
         modelName = [‘Model_’,geneNames{iGene}];
         saveName = [‘seqModels/’,modelName];
         logfile = [‘logFiles/log’,modelName];
         if iGene==2
            load(saveName)
            eval([modelName,’.formPropensitiesGeneral;’]);
         end
         cmd = SSIT.generateCommandLinePipeline(saveName,modelName,Pipeline,…
            pipelineArgs=pipelineArgs,saveFileOut=saveName, …
            logFile=logfile,runNow=true,runOnCluster=true);
         pause(60); % Pause to allow for matlab licenses to reset.
     end
end
~~~

**Figure B23:**
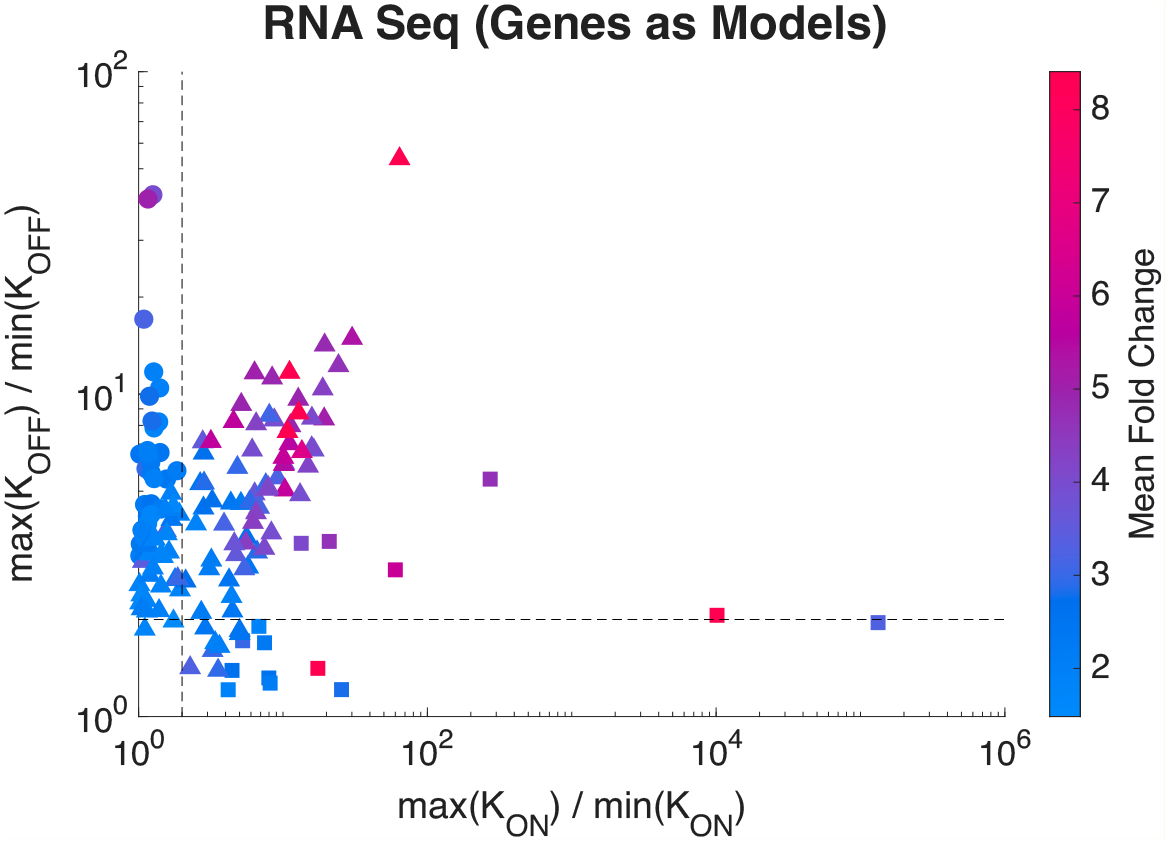
Transcriptional regulation across 151 dex-responsive genes inferred from scRNA-seq data [29]. For each gene, model parameters were fit using the SSIT pipeline code shown above, and these fitted parameters were used to compute the dex-induced fold change in the promoter switching rates, summarized as max(*K*_*ON*_)/min(*K*_*OF F*_). Points are colored by the maximum observed mean fold change in the data (log_2_ scale), and marker shapes indicate the dominant mode of regulation, separated based on the relative changes in switching rates (dashed lines denote two-fold thresholds): primarily increased activation (*K*_*ON*_; squares), primarily decreased deactivation (*K*_*OF F*_; circles), or mixed regulation (triangles).

### 2.6 The SSIT provides a graphical user interface (GUI) for novice users

In addition to its application programming interface (API), the SSIT also provides a graphical user interface (GUI) in MATLAB for interactive utilization of the main modeling and inference workflows. The SSIT GUI offers an intuitive and easy-to-use button- and tab-based environment for users rather than requiring the writing and working with scripts that can be extensive in some cases. Instead, users can load preset models or define or define their own species, parameters, propensities, stoichiometry, input expressions, and initial conditions (2.1); load, filter, and associate experimental or simulated data (2.3.1); perform simulations and solve probability distributions by ODEs (2.2.2), SSA (2.2.4), FSP (2.2.5), and hybrid methods (2.4.2); set likelihood (2.3.2) or loss functions (2.3.5); perform parameter inference through maximum likelihood estimation (2.3.3), Bayesian inference (2.3.4), or Approximate Bayesian Computation (2.3.5); compute sensitivities (2.2.7) and FIM (2.2.8); and visualize results, including simulated trajectories and FSP distributions, comparisons with empirical data, model fits and predictions, uncertainty measurements, and summary diagnostics and statistics. Models and settings specified in the GUI can be saved and used for command-line execution or batch scripts.

## 3 Discussion

The Stochastic System Identification Toolkit (SSIT) offers a platform for fast, accurate, and versatile mechanistic modeling solutions, parameter inference, information and uncertainty quantification, distortion handling, and quantitative experimental design. This tight integration between forward solvers, inference algorithms, and assessments of both model and experimental data is a central practical advantage of the SSIT.

### 3.1 Stochastic modeling and inference

Stochastic modeling is crucial for systems that experience significant fluctuations from mean behavior through time, thus affecting the inference of the underlying model and rate parameters which drive the system. Such systems include those with low copy numbers and high intrinsic noise. In the SSIT, simple, commonly used mechanistic models such as birth-death and toggle switch are already specified and readily called. Models can also be built from scratch by defining species, stoichiometry, and propensity functions, and the SSIT allows for highly complex, nonlinear functions (including logical statements, etc.) and time-varying input signals (2.3.1).

The Chemical Master Equation (CME) is a fundamental approach for describing the time evolution of probability distributions over discrete system states (2.2.1). Unfortunately, most interesting systems involve an infinite state space or a finite state space that is too large to produce computationally tractable numerical solutions. The Finite State Projection (FSP) approximation (2.2.5) truncates the full CME according to an modifiable error tolerance and computes not only the rigorously bounded subspace but also keep track of the total absolute error resulting from the loss of probability to sink states. Other important properties of the FSP lead to exact calculation of escape times and likelihood bounding for model fitting and parameter inference [**?**]. FSP is therefore a powerful tool for modeling stochastic systems, but its computational efficiency and scalability depend on the specific implementation of its state space truncation routine and evolution through time.

The SSIT specializes in producing efficient solutions of FSP truncations to the CME and integrating them into broader workflows for parameter inference, information and uncertainty quantification, and experiment design. The SSIT has also been carefully designed to efficiently simulate trajectories using parallel and GPU implementations of the Stochastic Simulation Algorithm (SSA) for exact solutions (2.2.4) and optionally averages system dynamics over time by deterministic ordinary differential equations (ODEs) - supposing large, well-mixed populations with negligible fluctuation or randomness (2.2.2) - hybridized deterministic-stochastic solutions (2.4.2), or moment closure approximations (2.2.3).

The SSIT provides full statistical inference tools for fitting mechanistic models to experimental data and estimating parameters, including maximum likelihood estimation (MLE) (2.3.3) for point estimation, Bayesian posterior sampling (2.3.4), and likelihood-free Approximate Bayesian Computation (2.3.5).

### 3.2 Sensitivity analysis, Fisher Information, and experiment design

The SSIT pairs probability distributions over system states with parameter sensitivity calculations (2.2.7) to compute Fisher Information (2.2.8), which quantifies the amount of information accessible about the values of the model parameters and can thus be used for informative experiment design (2.2.9). By computing Fisher Information matrices (FIM) over multiple realistic experimental and measurement constraints and using one of various scalar optimality criteria (2.2.9), the SSIT enables the tailoring of designs to achieve specific goals such as: minimizing overall parameter uncertainty, avoiding poorly identified directions, maximizing predictive accuracy over design space, or discriminating between competing mechanistic models/hypotheses. In practice, users can specify an experimental design variable (e.g., drug dosage, measurement times, or number of cells), and the SSIT evaluates FIM-based design scores by propagating FSP solutions and their sensitivities. The design scores can then be used in the SSIT within search or optimization routines to propose more informative experiments, or within sequential experiment design loops where the design is updated after each round of data collection and inference.

The pipeline from data and model to experiment design is fully transparent in the SSIT, and unlike in other popular methods [**?**], this transparency includes uncertainty measurements (2.3.4-2.3.6), model assessment and comparison, tunable distortion handling (2.4.3), and the option to automatically generate both local and cluster computing pipeline scripts (2.5).

### 3.3 Model assessment

The SSIT includes tools to critically assess model fit and to quantify parameter uncertainty. Likelihoods (2.3.2), Bayes factors, and Akaike and Bayesian Informaton Criteria are all comparable using varying mechanistic models and/or independent experimental measurements across different time points and experimental conditions. Building on this, the SSIT’s cross-validation module (2.3.6) provides a systematic way to assess predictive performance: data are partitioned into training and test subsets, MLEs (2.3.3) are obtained on each training partition, and their predictive log-likelihoods are evaluated on held-out data. The resulting cross-validation error summarizes how well the inferred model generalizes to unseen data and can reveal overfitting or model misspecification.

Additionally, the SSITMultiModel class (2.4.4) and cross-validation multi-model creator enable fitting the same mechanistic model to multiple replicas, doses, genetic variants, or other criteria while allowing selected parameters to vary between groups. The log-normal penalty formulation on replica-to-replica parameter deviations provides a principled way to encode prior expectations about batch variation or biological heterogeneity. This framework allows users to quantify which parameters are stable across conditions and which are sensitive to experimental context, and to test hypotheses about shared vs. condition-specific mechanisms.

### 3.4 Model reduction

For many realistic systems, the CME or FSP state space can be extremely large and an infeasible number of SSA trajectories are required for statistical significance. The SSIT therefore provides several model reduction strategies (2.4.1) that can be applied to accelerate both simulation and inference while preserving key dynamic behavior of the system.

The FSP itself acts as a controlled truncation of the CME, limiting the state space while bounding the truncation error. In addition, hybrid deterministic–stochastic methods (2.4.2) allow users to treat specified species (e.g., highly abundant species or those that undergo relatively fast reactions) with upstream reactions deterministically via ODEs, while computing fully stochastic realisations for others. This division can dramatically reduce computational cost without sacrificing fidelity in the most informative parts of the system.

Other model reduction (2.4.1) options include quasi-steady-state approximations (QSSA) for fast intermediates, lumping strategies (e.g., Log Lump QSSA) to aggregate states or compartments, and projection-based methods such as eigen-decomposition, proper orthogonal decomposition (POD), or dynamic mode decomposition (DMD) to construct low-dimensional bases from FSP solutions.

### 3.5 Distortion handling

The SSIT also contains options for handling extrinsic noise due to cell-to-cell variability or to measurement inaccuracies. Analysis of upstream cellular variability is achieved by sampling parameters over arbitrary distributions, whereas downstream measurement noise is analyzed using probabilistic distortion operators (PDOs, 2.4.3). In particular, the latter type of extrinsic noise - introduced by experimental protocols, measurement instrumentation or procedures, and downstream data-processing choices - distorts the observed data, subsequently transforming the corresponding probability distributions describing changes to the system states through time, and thereby interfering with the accuracy and precision of parameter inference, mechanistic insights, and predictions. Fortunately, in many cases, particular distortions introduced by cheaper, faster designs are systematic and therefore learnable and revertible to less distorted distributions by way of conditional probability. In the SSIT, such conditional distributions can be used as operators (i.e., PDOs) that map latent state distributions to observed data distributions through calibration from reference data. Users can choose from built-in parametric forms (e.g., affine-Poisson) or define custom distortion models.

PDOs can be incorporated directly into the SSIT modeling pipeline (2.5) to improve parameter estimates (2.3.2-,2.3.5), uncertainty quantification (2.3.4-2.3.6), and FSP-FIM-based experiment design (2.2.9). The SSIT thus jointly accounts for intrinsic stochasticity as well as additional extrinsic distortion when fitting models, making predictions, and designing subsequent experiments.

### 3.6 Pipelines, cluster computing, and the SSIT GUI

The SSIT generates simple, customizable pipelines (2.5) for execution in the background on local machines with optional parallelization or wrapped inside a schedular command (e.g., `sbatch` on `slurm`) to be run on cluster nodes. These scripts are particularly useful for analyses that involve larger systems with numerous parameters to fit, many forward simulations, long sampling chains to convergence, and multiple experiment design considerations.

The SSIT graphical user interface (GUI) (2.6) allows users to load preset models or define their own (2.1); load experimental or simulated data (2.3.1); simulate and solve models using ODEs (2.2.2), SSA (2.2.4), FSP (2.2.5), and hybrid methods (2.4.2); define likelihood (2.3.2); obtain point estimates in parameter space through maximum likelihood estimation (2.3.3) or infer posterior distributions by Bayesian inference (2.3.4); perform sensitivity analysis (2.2.7); compute FIMs (2.2.8); and optimize experiments (2.2.9); and visualize results, including simulated trajectories and FSP distributions, comparisons with empirical data, model fits and predictions, uncertainty measurements, and summary diagnostics and statistics. Models and settings specified in the GUI can be saved and used for command-line execution or batch scripts.

### 3.7 Data analyses

In this paper, we demonstrated features of the SSIT on two experimental datasets consisting of RNA count data from: (a) *Saccharomyces cerevisiae* yeast cells, focusing on the STL1 gene, in response to NaCl [33]; and (b) 151 genes from breast cancer cells in response, measured by single-cell RNA sequencing (scRNA-seq), to dexamethasone [29]. The STL1 data was modeled and fit using a complex input expression (Box 4) and four distinct gene states, each with their own transcription rate. The scRNA-seq data analysis made use of the SSIT’s complex modeling capabilities, including the incorporation of a binomial probabilistic distortion operator to handle dropout and other causes of a failure to detect counts as well as SSITMultiModel, which fit multiple genes with two shared parameters and seven distinct parameters per gene.

### 3.8 Comparison to other software

Tables 1 and 2 summarize a representative selection of software tools for simulating and analyzing reaction networks. Table 1 emphasizes the diversity of approaches, with many focusing primarily on deterministic ordinary differential equation (ODE) models, Gillespie’s Stochastic Simulation Algorithm (SSA) for sampling trajectories, or moment-based approximations. Comparatively few packages offer direct numerical solutions of the Chemical Master Equation (CME) and fewer still offer efficient implementations of the Finite State Projection [**?**] method for approximating the CME with rigorous error bounds, let alone fully developed pipelines from modeling and data processing to full statistical inference, distortion handling, and experiment design. Table 2 provides a more detailed comparison between the software in Table 1 that support FSP-based CME solutions, highlighting these additional tools offered by the SSIT.

**Table 1.** Popular mechanistic modeling software; ^∗^deterministic ODEs (means/concentration), as opposed to CME; ^*α*^model-builder only; ^*†*^”hybrid” in this table generally refers to network-based combinations of deterministic ODE/moment and stochastic model solutions, whereas BioNetGen uses “hybrid” to refer to network-free particle/population simulation and iBioSim 3 uses “hybrid” to refer to its dynamic flux-based analysis; ^*‡*^with the MomentClosure.jl [**?**] and FiniteStateProjection.jl [**?**] add-ons; ^+^moments computed under the CME solution via DeepCME; ^∼^steady-state noise estimation/Linear Noise Approximation; ^#^semi-Lagrangian partial integral differential equation (PIDE) approximation; ^*§*^ StochSS and URDME/PyURDME use Adaptive Diffusive Finite State Projection for spatial Reaction-Diffusion Master Equations and URDME uses spatial/RDME-SSA.

| Approach: |  | ODEs* | Moments | SSA | CME (FSP) | CME (other) | Hybrid |
| --- | --- | --- | --- | --- | --- | --- | --- |
| AMICI | [?] | ✓ |  |  |  |  |  |
| AMIGO2 | [?] | ✓ |  |  |  |  |  |
| BioCRNpyler<sup>α</sup> | [?] |  |  |  |  |  |  |
| BioNetGen | [?] | ✓ |  | ✓ |  |  | ✓<sup>†</sup> |
| Catalyst | [?] | ✓ | ✓<sup>‡</sup> | ✓ | ✓<sup>‡</sup> |  | ✓ |
| CERENA | [?] | ✓ | ✓ | ✓ | ✓ |  | ✓ |
| CmePy | [?] |  |  |  | ✓ |  |  |
| COPASI | [3] | ✓ |  | ✓ |  |  | ✓ |
| Data2Dynamics | [?] | ✓ |  |  |  |  |  |
| deBInfer | [?] | ✓ |  |  |  |  |  |
| DeepCME | [?] |  | ✓<sup>+</sup> |  |  | ✓ |  |
| Dizzy | [?] | ✓ | ✓<sup>~</sup> | ✓ |  |  |  |
| dMod | [?] | ✓ |  |  |  |  |  |
| GillesPy2 | [?] | ✓ |  | ✓ |  |  | ✓ |
| iBioSim 3 | [?] | ✓ |  | ✓ |  |  | ✓<sup>†</sup> |
| IDESS | [?] |  |  | ✓ |  | ✓<sup>#</sup> |  |
| iNA | [?] | ✓ | ✓<sup>~</sup> | ✓ |  | ✓ |  |
| MobsPy | [?] | ✓ |  | ✓ |  |  |  |
| modelbase | [?] | ✓ |  |  |  |  |  |
| MOCA | [?] |  | ✓ |  |  |  |  |
| Nessie | [?] |  |  | ✓ |  | ✓ |  |
| SELANSI | [?] |  |  |  |  | ✓<sup>#</sup> |  |
| SHAVE | [?] |  | ✓ |  |  | ✓ | ✓ |
| SimBiology | [5] | ✓ |  | ✓ |  |  |  |
| StochDynTools | [?] |  | ✓ |  |  |  |  |
| StochKit2 | [?] |  |  | ✓ |  |  |  |
| StochPy | [?] |  |  | ✓ |  |  |  |
| StochSS | [?] | ✓ |  | ✓ | ✓<sup>§</sup> |  |  |
| Tellurium | [4] | ✓ |  | ✓ |  |  |  |
| URDME | [?] |  |  | ✓<sup>§</sup> | ✓<sup>§</sup> |  |  |
| **The SSIT** |  | ✓ | ✓ | ✓ | ✓ |  | ✓ |

**Table 2.** Comparison of various software features offered by the packages listed in Table 1 that provide direct CME solutions by FSP; ^∗^using SSA-style trajectories; ^*†*^with SciML; ^*‡*^with Data2Dynamics [**?**]; ^*§*^with PyURDME; ^*α*^without extensive deubbing, using the current version of source code after installation per instructions, at time of submission; ^*β*^URDME requires additional setup for MEX compiler on macOS; ^∗∗^at time of submission.

|  | Catalyst<br>[?] | CERENA<br>[?] | CmePy<br>[?] | StochSS/URDME<br>[?] [?] | The<br>SSIT |
| --- | --- | --- | --- | --- | --- |
| **CME solutions** |  |  |  |  |  |
| Escape/waiting times | ✓* |  |  |  | ✓ |
| **State space adjust** |  |  |  |  |  |
| N-step expansion [?] |  |  | ✓ |  | ✓ |
| SSA |  |  |  |  | ✓ |
| Polynomial bounds |  |  |  |  | ✓ |
| **Estimation of $\theta$** |  |  |  |  |  |
| ABC | ✓<sup>†</sup> |  |  |  | ✓ |
| Maximum likelihood est. | ✓<sup>†</sup> | ✓<sup>‡</sup> |  | ✓ | ✓ |
| Bayesian/Monte Carlo | ✓<sup>†</sup> |  |  |  | ✓ |
| Cross validation |  |  |  |  | ✓ |
| **Sensitivities/FIM** |  |  |  |  |  |
| Finite difference |  |  |  |  | ✓ |
| Forward sensitivity | ✓<sup>†</sup> | ✓ |  | ✓ | ✓ |
| Fisher Information |  |  |  |  | ✓ |
| **Measurement noise** |  |  |  |  |  |
| Distortion operators [34] |  |  |  |  | ✓ |
| **Model Reduction** |  |  |  |  |  |
| Hybrid determ. + stoch. | ✓<sup>†</sup> | ✓ |  |  | ✓ |
| QSSA/Eigen. decomp. |  |  |  |  | ✓ |
| State smoothing |  |  |  |  | ✓ |
| Data-driven learning |  |  |  |  | ✓ |
| **Experiment Design** |  |  |  |  |  |
| FSP-FIM [?] |  |  |  |  | ✓ |
| Bayesian |  |  |  |  | ✓ |
| **Propensities** |  |  |  |  |  |
| Time-varying | ✓ | ✓ | ✓ |  | ✓ |
| Logicals | ✓ |  |  |  | ✓ |
| Non-linear | ✓ | ✓ | ✓ | ✓ | ✓ |
| **Compatibility** |  |  |  |  |  |
| SBML | ✓* | ✓ |  | ✓ | ✓ |
| SimBiology |  |  |  |  | ✓ |
| **Source Code/API** |  |  |  |  |  |
| Julia | ✓ |  |  |  |  |
| Python |  |  | ✓ | ✓<sup>§</sup> |  |
| MATLAB |  | ✓ |  | ✓ | ✓ |
| **User Interface** |  |  |  |  |  |
| GUI |  |  |  | ✓ | ✓ |
| Text/XML | ✓ | ✓ |  | ✓ | ✓ |
| IDE/Commandline | ✓ | ✓ | ✓ | ✓ | ✓ |
| **Accessibility** |  |  |  |  |  |
| GitHub | ✓ | ✓ | ✓ | ✓ | ✓ |
| GitHub example scripts run<sup>α</sup> | ✓ |  |  | ✓<sup>β</sup> | ✓ |
| Year last updated on GitHub** | 2026 | 2017 | 2018 | 2026 | 2026 |

Packages like Catalyst [**?**] (with FiniteStateProjection.jl) and CERENA [**?**] provide strong CME/ODE/moment capabilities, while CmePy [**?**] focuses narrowly on FSP and StochSS [**?**]/URDME [**?**] target SSA and diffusive FSP for spatial RDMEs. The SSIT complements these tools by combining (a) direct CME solutions by efficient FSP computations, iteratively expanding and contracting the truncation bounds while recomputing the resulting error, with (b) full statistical inference using likelihoods or posterior probability sampling, (c) experiment design using FSP-FIM and Bayesian utilities, (d) model-reduction & acceleration tools (hybrid ODE–FSP, POD, sparse grids, quasi-steady-state, etc.), and (e) explicit handling of extrinsic noise to allow for cost effective, efficient experimentation. Unlike model-construction libraries (e.g., BioCRNpyler [**?**], PySB [**?**]) or constraint-based frameworks (e.g., COBRA [**?**]), the SSIT is purpose-built for stochastic, mechanistic, time-evolving analyses and the data-model alignment issues that arise in modern experiments.

### 3.9 Reproducibility and interoperability

The SSIT reads models in the XML format of SBML [**?**, 2] and can interoperate with SimBiology [5]. The SSIT can be run from command line, through its own Graphical User Interface, or interactively within the MATLAB environment. The SSIT supports reproducible analyses with explicit random-seed control, versioned example scripts, and automated testing. To further ensure reproducibility, the SSIT contains a set of automated tests that check SSIT command line results against known analytical results, and these that are executed every time a push is made to SSIT on GitHub, and test results are visible to the public at: https://github.com/MunskyGroup/SSIT/actions. The graphical functions of the SSIT GUI can be tested by running the function `testGui.m` in the `tests` folder.

### 3.10 Limitations and future directions

At present, the SSIT targets well-mixed systems. The current version does not yet include spatial Reaction-Diffusion Master Equation models or DFSP, although diffusive modeling in the SSIT is currently under development. Second, while hybridization, POD/sparse-grid surrogates, and N-step/SSA-guided expansion extend FSP to moderately large systems, very high-dimensional networks can still be challenging.

Finally, the SSIT’s core implementation in MATLAB eases integration with SimBiology/SBML pipelines and allows for the use of built-in toolkits to expand the SSIT’s capabilities in the future, but MATLAB may not match the raw performance of low-level Julia/C++ stacks under some regimes. Wrapping core solvers in Julia/C++ could broaden performance and providing portable Python-based workflows may allow for broader adoption.

## 4 Conclusion

The Stochastic System Identification Toolkit (SSIT) is an open-source, versatile, actively developed software package based in MATLAB that provides an integrated pipeline from (i) mechanistic model building and simulation; to (ii) data importation, filtering by condition, and connection to model species; to (iii) model fitting/parameter inference based on likelihood, Bayesian, and simulation/loss function; to (iv) data distortion handling conditional probability operators [34]; to (v) parameter sensitivity analysis, calculation of Fisher Information [**?**], and experiment design using

FSP-FIM [**?**]. Additionally, the SSIT includes several model reduction techniques and options for automated parallelization within the MATLAB environment, on command line, or via high-performance cluster computing to increase speed.

The SSIT makes use of built-in MATLAB packages while remaining fully open-source and readily accessible, with full documentation and detailed examples. The source code is freely available with README, download and installation instructions, and example scripts for every tool discussed in this paper at: http://github.com/MunskyGroup/SSIT/.

## 5 Acknowledgments

This research was supported by the NIH National Institute for General Medical Sciences (Award R35GM124747) and the National Science Foundation (Award MCB-1941870). The authors would like to thank Rachel Keating, Torin Moore, Stuart McKnight, Ethan Green, Eric Ron, and Joshua Cook for feedback and assistance with testing throughout the development of the the SSIT.

