## Supporting Information for "The Stochastic System Identification Toolkit (SSIT) to model, fit, predict, and design experiments"

March 7, 2026

### 1 Features & Functionality

The Stochastic System Identification Toolkit (SSIT) allows users to specify and efficiently solve the Chemical Master Equation for discrete stochastic models. The SSIT is especially useful for, but not limited to, analyses of single-cell gene regulation. The SSIT functionality includes the following:

**Function 1: Create, save, and load models defined by their species, parameters, propensity functions, stoichiometries, and initial conditions**

- Easily formulate model stoichiometries and propensity functions, including for non-linear models, time-varying propensity functions (e.g., environmental input signals), and propensities defined by logical switches
- Save, load, and modify models
- Import and export models to SimBiology or SBML formats to use with other computational approaches

**Function 2: Solve models**

- Ordinary differential equations (ODE)
- Basic moment closure analysis
- Gillepsie's Stochastic Simulation Algorithm (SSA) trajectories
- Finite State Projection (FSP) truncation of the Chemical Master Equation
- Compute first passage or escape time distributions

**Function 3: Sensitivity analysis**

- Sensitivity of model distributions to parameter variations
- Gradients of likelihood functions

**Function 4: Fisher Information Matrix calculation**

- Estimate parameter inference uncertainties
- Optimize designs of experiment

**Function 5: Fitting data and model parameter estimation**

- Likelihood calculation, parameter sweeps, and maximum likelihood estimation searches
- Bayesian inference, Markov chain Monte Carlo, Metropolis-Hastings
- Cross-Validation

**Function 6: Complex modeling**

- "Multi models" to infer shared parameter sets across models associated with different data sets / experimental conditions
- Model reduction (reduced order models) by:
  - Hybrid solutions (deterministic + stochastic populations)
  - Quasi-Steady State Approximations (QSSA) on fast species
  - Eigenvalue decomposition
  - Coarse meshes
  - Principle orthogonal decomposition (POD)
- Model and account for distortion of data due to extrinsic experimental, measurement, and data processing noise by calibrating empirical probability distortion operators (PDOs) to quantify effects of data distortion

- Pipelines for complex modeling and cluster computing

**Function 7: Sequential experiment design**

- FSP-FIM approach to produce optimally informative experiments
- Probabilistic inference from Bayesian sequential experiment design
- Adjustment of data distortion effects in design comparison

#### 1.1 Model Summary

The model summary that is printed to the command window when 'STL1_4state.summarizeModel' is called is shown below:

```
Species:
     g1; IC = 1;  discrete stochastic
     g2; IC = 0;  discrete stochastic
     g3; IC = 0;  discrete stochastic
     g4; IC = 0;  discrete stochastic
     mRNA; IC = 0;  discrete stochastic

Reactions:
  Reaction 1:
     s1: 1*g1 --> 1*g2
     w1: k12*g1
  Reaction 2:
     s2: 1*g2 --> 1*g1
     w2: (max(0,k21o*(1-k21i*Hog1)))*g2
  Reaction 3:
     s3: 1*g2 --> 1*g3
     w3: k23*g2
  Reaction 4:
     s4: 1*g3 --> 1*g2
     w4: k32*g3
  Reaction 5:
     s5: 1*g3 --> 1*g4
     w5: k34*g3
  Reaction 6:
     s6: 1*g4 --> 1*g3
     w6: k43*g4
  Reaction 7:
     s7: NULL --> 1*mRNA
     w7: kr1*g1
  Reaction 8:
     s8: NULL --> 1*mRNA
     w8: kr2*g2
  Reaction 9:
     s9: NULL --> 1*mRNA
     w9: kr3*g3
  Reaction 10:
     s10: NULL --> 1*mRNA
     w10: kr4*g4
  Reaction 11:
     s11: 1*mRNA --> NULL
     w11: dr*mRNA

Input Signals:
     Hog1(t) = A*(((1-(exp(1)^(-r1*(t-t0))))*exp(1)^(-r2*(t-t0)))/...
                 (1+((1-(exp(1)^(-r1*(t-t0))))*exp(1)^(-r2*(t-t0)))/M))^n*(t>t0)

Model Parameters:
    {'t0'  }    {[    3.1700]}
    {'k12' }    {[        78]}
    {'k21o'}    {[    192000]}
    {'k21i'}    {[      3200]}
    {'k23' }    {[    0.4020]}
    {'k34' }    {[    7.8000]}
```

```
{‘k32’ }    {[    1.6200]}
{‘k43’ }    {[    2.2800]}
{‘dr’  }    {[    0.2940]}
{‘kr1’ }    {[    0.0468]}
{‘kr2’ }    {[    0.7200]}
{‘kr3’ }    {[   59.4000]}
{‘kr4’ }    {[    3.2400]}
{‘r1’  }    {[    0.0041]}
{‘r2’  }    {[    0.4260]}
{‘A’   }    {[9.3000e+09]}
{‘M’   }    {[6.4000e-04]}
{‘n’   }    {[    3.1000]}
```

#### 1.2 SBML: Systems Biology Markup Language

The Systems Biology Markup Language (SBML) [57, 58] enables consistent sharing, loading, and interoperability of models across different research groups and software tools, such as COPASI [40], Tellurium [54], and MATLAB. SBML is based on models that can be represented mathematically using ordinary differential equations (ODEs) or stochastic descriptions. The standardized nature of SBML promotes model reproducibility in systems biology and synthetic biology research.

Built in MATLAB, the SSIT facilitates the creation, storing, interoperability, and reloading of SBML models. The Bursting Gene model example shown in Fig 3 in the main text would look something like this:

```
<?xml version="1.0" encoding="UTF-8"?>
<sbml xmlns="http://www.sbml.org/sbml/level2/version4"
level="2" version="4">
  <model id="BurstingGeneExpression" name="BurstingGeneExpression">

    <!-- List of parameters -->
    <listOfParameters>
      <parameter id="kon" value="0.2"/>
      <parameter id="koff" value="0.2"/>
      <parameter id="kr" value="100"/>
      <parameter id="gr" value="5"/>
    </listOfParameters>

    <!-- List of species -->
    <listOfSpecies>
      <species id="offGene" compartment="cell" initialAmount="1"/>
      <species id="onGene"  compartment="cell" initialAmount="0"/>
      <species id="mRNA"    compartment="cell" initialAmount="0"/>
    </listOfSpecies>

    <!-- Compartment (required even if only one) -->
    <listOfCompartments>
      <compartment id="cell" size="1"/>
    </listOfCompartments>

    <!-- List of reactions -->
    <listOfReactions>
      <!-- offGene to onGene -->
      <reaction id="r1" reversible="false">
        <listOfReactants>
          <speciesReference species="offGene" stoichiometry="1"/>
        </listOfReactants>
        <listOfProducts>
          <speciesReference species="onGene" stoichiometry="1"/>
        </listOfProducts>
        <kineticLaw>
          <math xmlns="http://www.w3.org/1998/Math/MathML">
            <apply><times/>
              <ci> kon </ci>
              <ci> offGene </ci>
            </apply>
          </math>
        </kineticLaw>
```

```
      </reaction>

      <!-- onGene to offGene -->
      <reaction id="r2" reversible="false">
        <listOfReactants>
          <speciesReference species="onGene" stoichiometry="1"/>
        </listOfReactants>
        <listOfProducts>
          <speciesReference species="offGene" stoichiometry="1"/>
        </listOfProducts>
        <kineticLaw>
          <math xmlns="http://www.w3.org/1998/Math/MathML">
            <apply><times/>
              <ci> koff </ci>
              <ci> onGene </ci>
            </apply>
          </math>
        </kineticLaw>
      </reaction>

      <!-- mRNA production -->
      <reaction id="r3" reversible="false">
        <listOfReactants>
        </listOfReactants>
        <listOfProducts>
          <speciesReference species="mRNA" stoichiometry="1"/>
        </listOfProducts>
        <kineticLaw>
          <math xmlns="http://www.w3.org/1998/Math/MathML">
            <apply><times/>
              <ci> kr </ci>
              <ci> onGene </ci>
            </apply>
          </math>
        </kineticLaw>
      </reaction>

      <!-- mRNA degradation -->
      <reaction id="r4" reversible="false">
        <listOfReactants>
          <speciesReference species="mRNA" stoichiometry="1"/>
        </listOfReactants>
        <listOfProducts>
        </listOfProducts>
        <kineticLaw>
          <math xmlns="http://www.w3.org/1998/Math/MathML">
            <apply><times/>
              <ci> gr </ci>
              <ci> mRNA </ci>
            </apply>
          </math>
        </kineticLaw>
      </reaction>
    </listOfReactions>
  </model>
</sbml>
```

#### 1.3 SimBiology: MATLAB Toolbox for Systems Pharmacology and Biology

SimBiology [51] is a MATLAB-based MathWorks toolbox for building, simulating, and analyzing quantitative dynamic models, primarily for applications in pharmacokinetics/pharmacodynamics (PK/PD) systems. SymBiology supplies a graphical user interface for the user to interactively construct models through a block diagram editor. As it is constructed in MATLAB, the user can also build SymBiology models in the MATLAB programming language.

SimBiology provides for modeling processes such as drug metabolism, efficacy, and safety, signaling pathways, gene expression, and optimization of dosing schedules using ODEs, stochastic simulations, global and local parameter sensitivity analysis, and compartmental modeling. It is also compatible with SBML, utilizing the standardized format to facilitate collaboration and integration with other modeling platforms. [58] [51]

### 2 Solving reaction models

The SSIT efficiently computes solutions to master equations by way of ordinary differential equations (ODEs), Stochastic Simulation Algorithm (SSA) trajectories, and - most significantly - Finite State Projection (FSP) truncation of the full chemical master equation (CME).

#### 2.1 Ordinary Differential Equations

The full ODE description of the approximation for $\mathbb{E}\{\mathbf{x}\}$ (Eq 7 in the main text) is:

$$\frac{d\mathbb{E}\{\mathbf{x}\}}{dt} = \mathbf{S}w = \sum_{\mu=1}^{N} \boldsymbol{\nu}_\mu w_\mu(\mathbb{E}\{\mathbf{x}\}) = \boldsymbol{\nu}_1 w_1(\mathbb{E}\{\mathbf{x}\}) + \boldsymbol{\nu}_2 w_2(\mathbb{E}\{\mathbf{x}\}) + ... + \boldsymbol{\nu}_N w_N(\mathbb{E}\{\mathbf{x}\}), \quad (1)$$

The Bursting Gene model (Fig 3 in the main text) can be described by a system of ODEs for its three species (two gene states plus mRNA): $\mathbf{G_{OFF}}$, $\mathbf{G_{ON}}$, $\mathbf{mRNA}$. The ODEs in component form are:

$$\begin{aligned}
\frac{d\mathbf{G_{OFF}}}{dt} &= -w_1 + w_2 && = -k_{on} \cdot \mathbf{G_{OFF}} + k_{off} \cdot \mathbf{G_{ON}} \\
\frac{d\mathbf{G_{ON}}}{dt} &= w_1 - w_2 && = k_{on} \cdot \mathbf{G_{OFF}} - k_{off} \cdot \mathbf{G_{ON}} \\
\frac{d\mathbf{mRNA}}{dt} &= w_3 - w_4 && = k_r \cdot \mathbf{G_{ON}} - d_r \cdot \mathbf{mRNA}
\end{aligned}$$

The basic version of the STL1 model is identical to the Bursting Gene model, but includes a simple time-varying input signal: $\text{Hog1}(t) = a_0 + a_1 e^{-r_1 t}\big(1 - e^{-r_2 t}\big)(t > 0)$. The ODEs are then:

$$\begin{aligned}
\frac{d\mathbf{G_{OFF}}}{dt} &= -w_1 + w_2 && = -k_{on} \cdot \mathbf{G_{OFF}} + k_{off} \cdot \mathbf{G_{ON}}/(1 + \mathbf{Hog1}) \\
\frac{d\mathbf{G_{ON}}}{dt} &= w_1 - w_2 && = k_{on} \cdot \mathbf{G_{OFF}} - k_{off} \cdot \mathbf{G_{ON}}/(1 + \mathbf{Hog1}) \\
\frac{d\mathbf{mRNA}}{dt} &= w_3 - w_4 && = k_r \cdot \mathbf{G_{ON}} - d_r \cdot \mathbf{mRNA}
\end{aligned}$$

The 4-state STL1 model (Box 1 in the main text) ODEs in component form are:

$$\begin{aligned}
\frac{d\mathbf{g_1}}{dt} &= -w_1 + w_2 && = -k_{12} \cdot \mathbf{g_1} + \max(0, k_{21o} \cdot (1 - k_{21i} \cdot \text{Hog1})) \cdot \mathbf{g_2} \\
\frac{d\mathbf{g_2}}{dt} &= w_1 - w_2 - w_3 + w_4 && = k_{12} \cdot \mathbf{g_1} - \max(0, k_{21o} \cdot (1 - k_{21i} \cdot \text{Hog1})) \cdot \mathbf{g_2} + k_{32} \cdot \mathbf{g_3} \\
\frac{d\mathbf{g_3}}{dt} &= w_3 - w_4 - w_5 + w_6 && = k_{23} \cdot \mathbf{g_2} - (k_{32} + k_{34}) \cdot \mathbf{g_3} + k_{43} \cdot \mathbf{g_4} \\
\frac{d\mathbf{g_4}}{dt} &= w_5 - w_6 && = k_{34} \cdot \mathbf{g_3} - k_{43} \cdot \mathbf{g_4} \\
\frac{d\mathbf{mRNA}}{dt} &= w_7 - w_8 && = k_{r4} \cdot \mathbf{g_4} - d_r \cdot \mathbf{mRNA}
\end{aligned}$$

In Fig. S1, we show the ODE solutions for the Bursting Gene and simplified STL1 models.

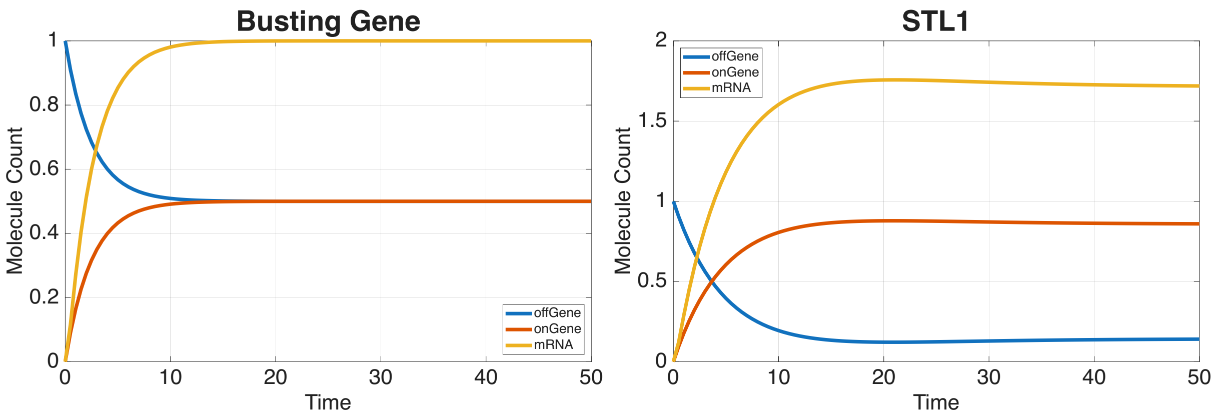


Figure S1: **ODE solution plots** for the Bursting Gene model from Fig. 3 in the main text and the simplified STL1 yeast gene model, which uses the Bursting Gene model but with a time-varying input of Hog1.

### 2.2 Moment Closure

Here, we illustrate how to compute moment solutions in the SSIT using the 4-state STL1 model (Fig B1 in the main text) and plot the means and standard deviations. The results are shown in Fig. S2.

```
% Set solution scheme to ‘moments’ and solve:
STL1_4state_mom.solutionScheme = ‘moments’; [~,~,STL1_4state_mom] = STL1_4state_mom.solve;

% Number of species:
nSp = numel(STL1_4state_mom.species);

%% First moments from the "moments" solver
% Moments: [ (#means + #secondMoments) x nTimes ]
mom = STL1_4state_mom.Solutions.moments;

% First nSp rows are the means ⟨x_i⟩
means_mom = mom(1:nSp, :);   % size: nSp x nTimes

%% Species trajectories from the ODE solver
% Get ODE solution:
Y_ode = STL1_4state_ODE.Solutions.ode;   % nTimes x nSpecies
means_ode = Y_ode.’;                     % transpose: nSp x nTimes

%% Compare ODEs vs moments
% Compare mRNA:
i_mRNA = find(strcmp(STL1_4state_mom.species,‘mRNA’));
den_mRNA = max(abs(means_ode(i_mRNA,:)), 1e-12);
errMean_mRNA = max(abs(means_mom(i_mRNA,:) - means_ode(i_mRNA,:)) ./ den_mRNA);

% Compare all species:
den_all = max(abs(means_ode), 1e-12);
errMean_all = max(max(abs(means_mom - means_ode) ./ den_all));
tol = 1e-2;   % 1% tolerance

if isequal(errMean_mRNA<tol,true)
    disp(‘4-state STL1: mRNA mean from moments matches ODE within 1%’);
else
    disp(‘4-state STL1: mRNA mean from moments does not match ODE within 1%’);
end
if isequal(errMean_all<tol,true)
    disp(‘4-state STL1: species means from moments match ODE within 1%’);
else
    disp(‘4-state STL1: species means from moments do not match ODE within 1%’);
end

%% Plot results:
STL1_4state_mom.plotMoments(STL1_4state_mom.Solutions,...
```

```
    STL1_4state.species(5), "meansanddevs", STL1_4state_mom.tSpan, [],...
    {'linewidth',4}, Title='4-state STL1 (mRNA)', TitleFontSize=26,...
    LegendLocation='east', Colors=[0.23,0.67,0.20], LegendFontSize=20,...
    YLabel='Molecule Count', AxisLabelSize=20, TickLabelSize=20);

STL1_4state_mom.plotMoments(STL1_4state_mom.Solutions,...
    STL1_4state.species(1:4), "meansanddevs", STL1_4state_mom.tSpan, [],...
    {'linewidth',4}, Title='4-state STL1 (mRNA)', TitleFontSize=26,...
    LegendLocation='east', YLabel='Molecule Count',...
    AxisLabelSize=20, TickLabelSize=20, LegendFontSize=20);
```

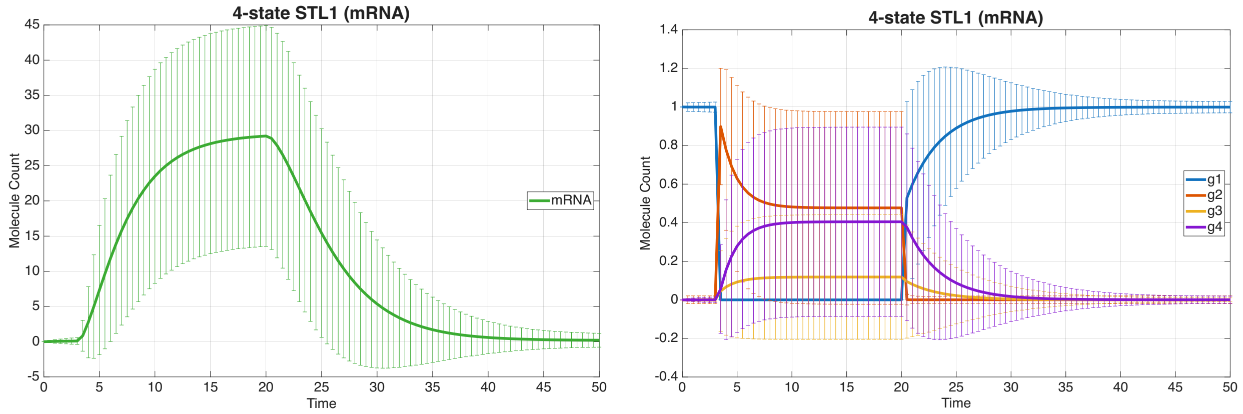


Figure S2: **Moment solutions** for the 4-state STL1 model species 'mRNA' (left) and the four gene states (right). The mean is shown as the solid lines and the standard deviation is shown using vertical bars of the same color.

#### 2.3 Stochastic Simulation Algorithm

Gillespie's stochastic simulation algorithm (SSA) [7,8] provides one method for generating statistically correct sample paths of the inherently discrete Chemical Master Equation (CME), which governs the probability distribution over molecular counts with respect to time. The SSA is a kinetic Monte Carlo (KMC) method paired with random number generators to sample the time evolution as the process moves between points in discrete state space [72]. Fundamentally, Gillespie's SSA generates a population trajectory for the system by following two key steps:

1. Compute the waiting time until the next reaction occurs.
2. Decide which reaction will occur.

The SSA decides upon the next reaction and the state to which it leads based on the propensity functions $\boldsymbol{w}$ and stoichiometry matrix $\mathbf{S}$, respectively. The behavior of the system is thus simulated from an initial state $\mathbf{x}_0$ and steps probabilistically along a series of states chosen using $\mathbf{S}$ and $\boldsymbol{w}$ until a specified final time $t_N$. The SSA proceeds through these steps according to the following procedure:

- Compute the propensity functions $w_\mu(x)$. The propensity functions give probabilities per unit time that a possible reaction $\mu$ will be the next reaction to fire based upon the current state $x$.
- Sum the current propensities: $w_0(x) = \sum_{\mu=1}^{N} w_\mu(x)$.
- Sample the exponentially distributed waiting time $\tau \sim \mathrm{Exp}(\mathrm{w}_0(\mathbf{x}))$ to the next reaction at rate $w_0(\mathbf{x})$ by sampling a uniform random number $r_1 \sim$ Uniform(0,1) and then $\tau$ using the transformation $\tau = \frac{1}{w_0}\ln\left(\frac{1}{r_1}\right)$.
- Select the next reaction index, $k$, according to the categorical probability distribution $k \sim \mathrm{Cat}\{\mathrm{w}_1/\mathrm{w}_0, \ldots, \mathrm{w}_\mathrm{N}/\mathrm{w}_0\}$ by drawing a second uniform random number $r_2 \sim$ Uniform(0,1) and finding the smallest reaction index $k$, so that: $\sum_{\mu=1}^{k-1} w_\mu < r_2 w_0 \leq \sum_{\mu=1}^{k} w_\mu$.

- Update the current time $t \leftarrow t + \tau$. If the new $t < t_{\text{MAX}}$, then update the system state $\mathbf{x}_1 \leftarrow \mathbf{x}_0 + \boldsymbol{s}_k$ according to the $k$-indexed stoichiometric updates $\boldsymbol{s}_k$ of the chosen reaction $\mu_k$.
- Repeat ↺

In Fig. S3, we show SSA trajectories and frequencies for each of the three species from the Bursting Gene and simplified STL1 models.

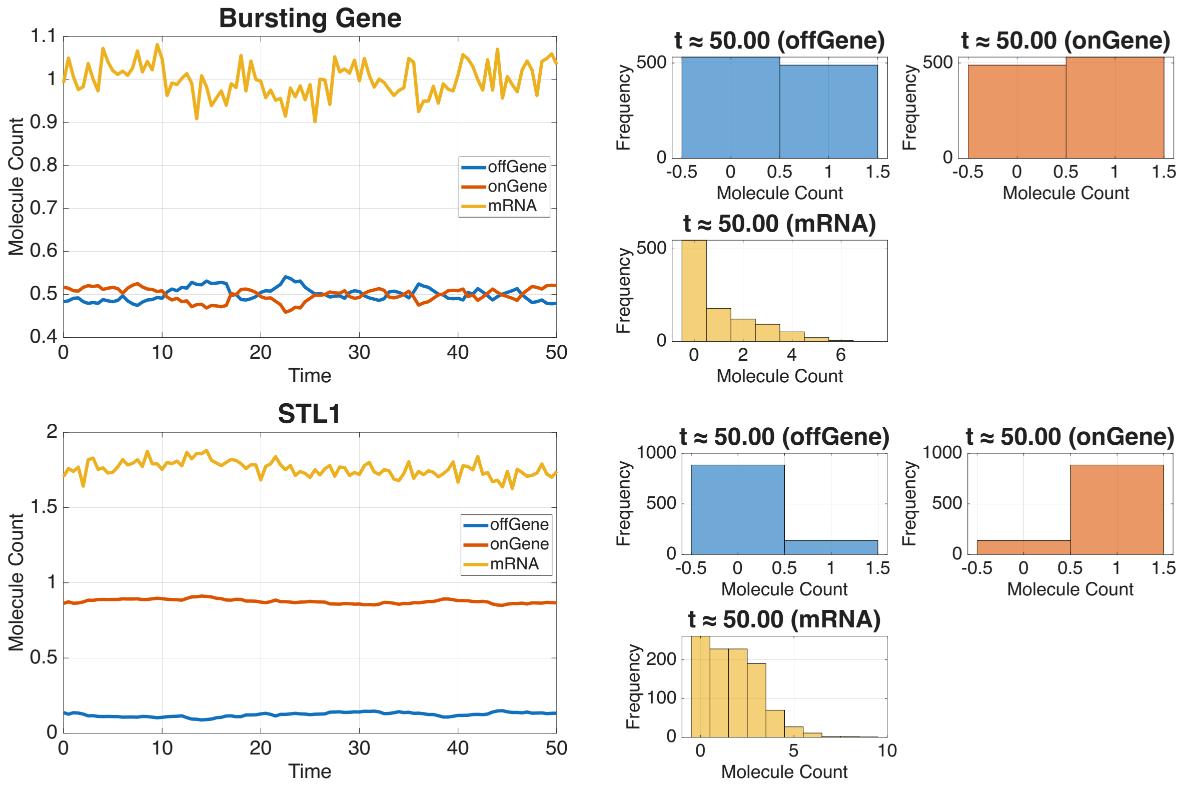


Figure S3: **SSA trajectories** (left, molecular counts over time) and distributions (right, frequencies of molecular counts across trajectories) for the Bursting Gene model from Fig 3 in the main text and the simplified STL1 model with a time-varying input of Hog1.

### 2.4 Finite State Projection

In the SSIT, the user sets this maximum error threshold using a property that defines the total FSP tolerance, 'fspOptions.fspTol' (which as a default is set to 0.001). The SSIT implementation of the FSP algorithm uses N-step/SSA-guided expansion and polynomial bounds to iteratively contract and expand the truncated subspace of the full CME, recomputing the error after each change and recalculating the likelihood when appropriate.

Fig. S4 shows plots of FSP means and standard deviations, as well as marginal distributions for each species of the Bursting Gene and simplified STL1.

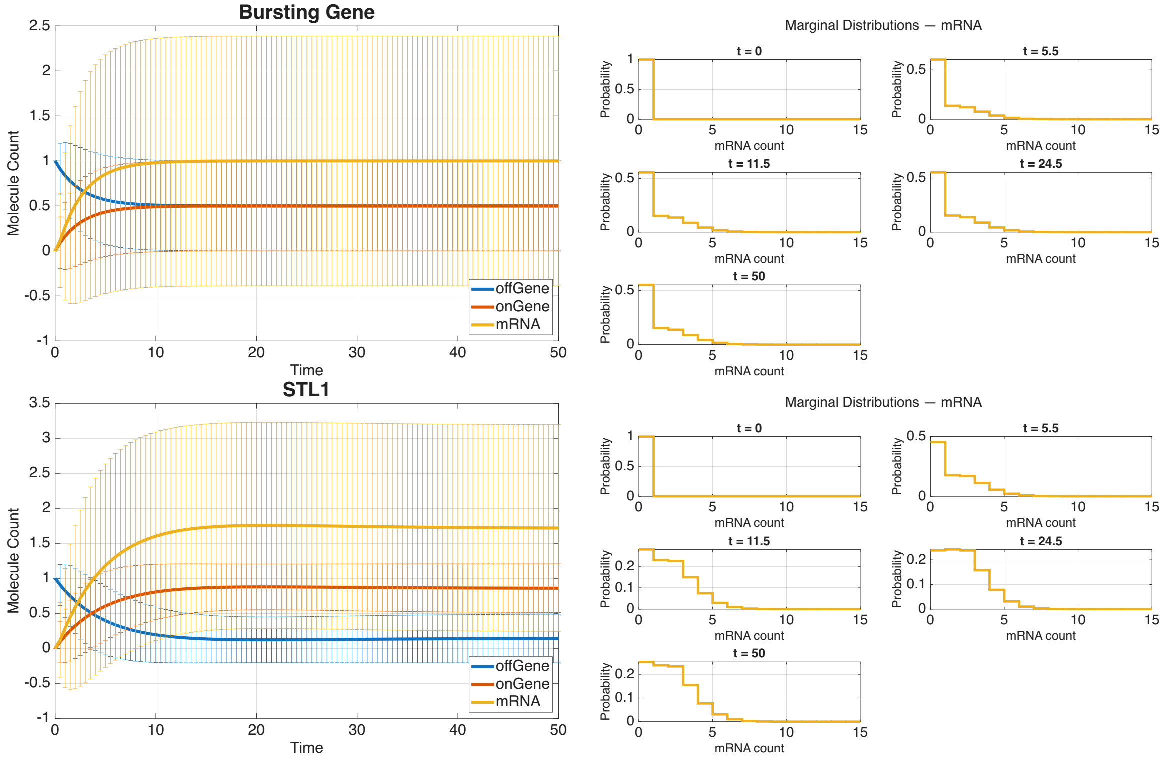


Figure S4: **FSP means and standard deviations (left) and marginal distributions for mRNA (right)** for the Bursting Gene (top) and simplified STL1 (bottom) models.

### 2.5 Escape and waiting time calculations

Equivalent to $\mathbb{E}[\tau_{i\to J_1}] = \int_0^\infty \left(1 - F_{J_1}(t)\right) dt$ from the main text, the mean escape time from an initial state $i$ to an absorbing target set $J_1$ can be obtained by solving the standard system of linear equations

$$-1 \;=\; \sum_k a_{ik}\, \mathbb{E}[\tau_{k\to J_1}], \quad i \notin J_1, \qquad \mathbb{E}[\tau_{j\to J_1}] = 0 \;\text{ for } j \in J_1, \tag{2}$$

where $a_{ik}$ are the entries of the generator matrix $A$ [**?**, 71, 74].

As a example, consider the simple case, where $J$ contains only a single state $J = i$ and $J_{\text{esc}}$ is all other states ($J'$ is empty). Eq. 15 from the main text becomes:

$$\frac{d}{dt}\begin{bmatrix} P_J^{\text{FSP}}(t) \\ z_{\text{esc}}(t) \\ z(t) \end{bmatrix} = \begin{bmatrix} -r_i & 0 & 0 \\ r_i & 0 & 0 \\ 0 & 0 & 0 \end{bmatrix} \begin{bmatrix} P_J^{\text{FSP}}(t) \\ z_{\text{esc}}(t) \\ z(t) \end{bmatrix},$$

where $r_i$ is the rate to leave the state $\mathbf{x}_i$. We can solve this simple equation for

$$F_J(t) = z_{\text{esc}}(t) = 1 - e^{-r_i},$$

corresponding to an exponentially distributed waiting time. The mean waiting time is then $\mathbb{E}[T_i] = \int_0^\infty e^{-r_i}\, dt = 1/r_i$, as expected for the a one step exponential waiting time [71, 74].

Fig. S5 shows escape CDFs and PDFs for the Bursting Gene and simplified STL1 models.

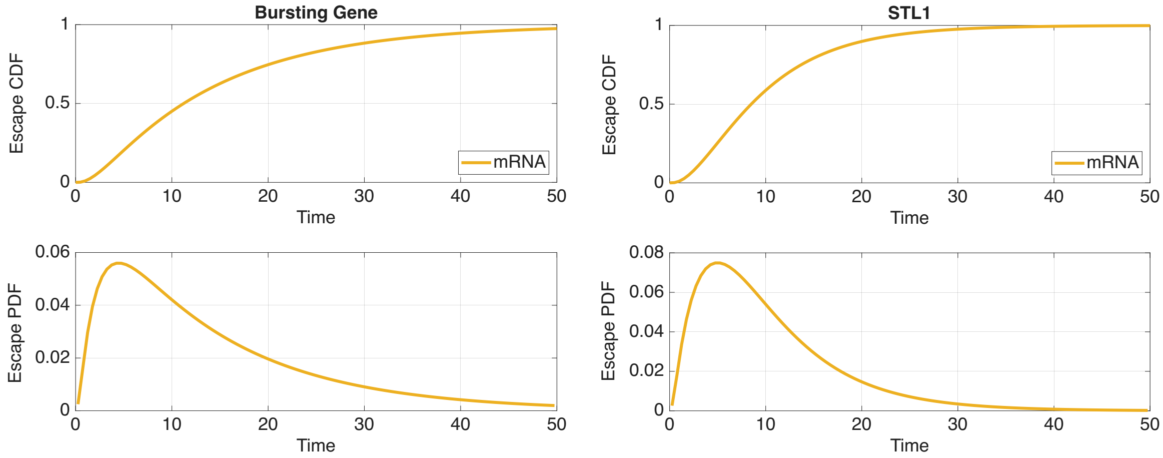


Figure S5: **Escape time CDFs and PDFs** for **mRNA** in the Bursting Gene model (left) and simplified STL1 model (right).

### 3 Sensitivity analysis and Fisher Information Matrix

Fig. S6 shows sensitivities and Fig. S7 shows FIM results ($\theta$-heat maps, per-$\theta$ information, spectra, and ellipses) for the Bursting Gene and simplified STL1 models.

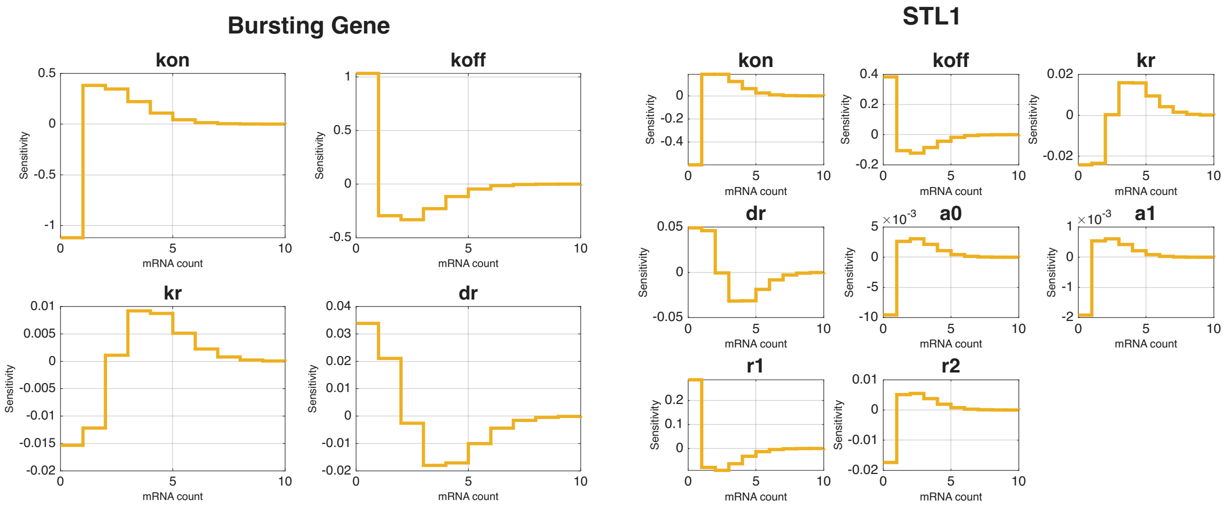


Figure S6: **Sensitivities** for each model parameter with respect to mRNA count for (left) the Bursting Gene and (right) simplified STL1 models.

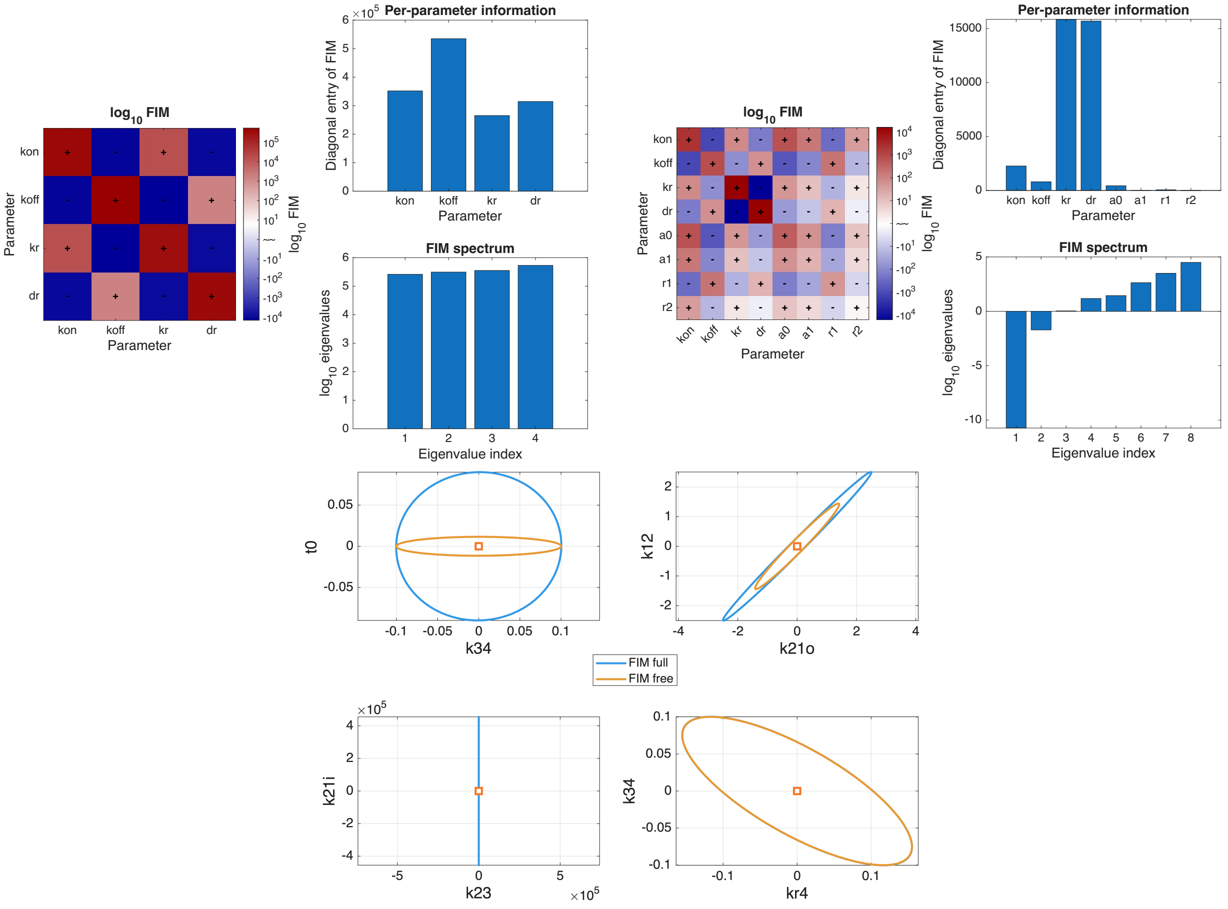


Figure S7: **FIM results** for (top, left) the Bursting Gene and (top, right) the simplified STL1 models are summarized by $\theta$-heat maps, per-$\theta$ information, and FIM spectra. (Bottom) Ellipses illustrating $\theta$-$\theta$ relationships for a subset of 4-state STL1 model parameters, comparing FIM results when all 18 model parameters are free to vary (FIM full) and results when only the 13 non-input signal parameters are fit, since the input signal parameters are experimentally determined.

#### 3.1 FIM optimality criteria and experiment design

There are several criteria that may be employed to optimize the next set of experiments based on the FIM. Currently, the SSIT uses E-optimality by default and comes equipped with the following FIM metrics:

- D-optimality (full FIM and subsets; via determinant of FIM or covariance)
- E-optimality (max smallest eigenvalue)
- Trace-based FIM optimality (maximize trace(FIM))
- Ds-style criteria for subsets or grouped parameters

These criteria are set with the argument 'FIMmetric':

- 'D-opt' - maximize the expected determinant of the FIM
- 'D-cov' - minimize the expected determinant of MLE covariance
- 'E-opt' - maximize the smallest eigenvalue of the FIM
- 'Trace' - maximize the trace of the FIM
- 'D-opt-sub-inv[$< i1 >, < i2 >, ...$]' - minimize the determinant of the inverse FIM for the specified indices, (all other parameters are assumed to be free)
- 'D-opt-sub[$< i1 >, < i2 >, ...$]' - maximize the determinant of the FIM for the specified indices, (only the parameters in 'obj.fittingOptions.modelVarsToFit' are assumed to be free)

Below we provide some elaborate on different FIM optimality criteria:

- **D-optimality:** D-optimality seeks to maximize the determinant of the FIM, thereby minimizing the volume of the confidence ellipsoid for $\theta$ and thus reducing overall parameter uncertainty and yielding balanced Fisher Information over all model parameters. The balanced approach of D-optimality has upsides and downsides; by definition it does not prioritize parameters of interest and it can be sensitive to parameters with high collinearity, producing near-singular FIMs in poorly identifiable models. D-optimality has the potential to overweight highly sensitive directions thus ignoring that some parameters may already be well estimated. However, the direct aim to reduce parameter uncertainty is intuitive and also a potential advantage of D-optimality is that it results in the maximization of the differential Shannon information of the parameter estimates.
  - Target: $\mathbf{max(det}(\mathcal{I}(\theta)))$, equivalent: $\mathbf{min}(\mathcal{I}^{-1}(\theta))$
  - Relative computational efficiency: **Slow**
- **Ds-optimality:** A variation of D-optimality is Ds-optimality, which maximizes the determinant of a sub-matrix of the FIM to reduce uncertainty for a particular subset of parameters. This may be especially useful for hierarchical models.
  - Target: $\mathbf{max(det}(\mathcal{I}_s(\theta)))$, equivalent: $\mathbf{min}(\mathcal{I}_s^{-1}(\theta))$
  - Relative computational efficiency: **Fast**
- **E-optimality:** E-optimality maximizes the minimum eigenvalue of the FIM, an approach that aspires for the informed estimation of all model parameters by avoiding flattened FIM space in any direction. E-optimality essentially computes worst-case scenarios for proposed experiment designs. This can be useful when a model borders on the unidentifiable in certain directions; however, the flip side is that in situations where worst-case scenarios are edge cases and E-optimality's inherent focus on improving a potentially weakly identifiable parameter neglects experiments that benefit the inference of more estimable parameters or parameters that are of more interest to the researcher.
  - Target: $\mathbf{max}(\lambda(\mathcal{I}(\theta)))$
  - Relative computational efficiency: **Slow**

### 4 Loading and Fitting Experimental Data

In the following subsections, we provide additional details about data fitting methods in the SSIT.

#### 4.1 Maximum Likelihood Estimation

It is common to work with log-likelihoods as they transform products into sums which are easier to worth with, numerically more stable, and often easier to optimize; accordingly, we maximize the log-likelihood function with respect to parameters $\boldsymbol{\theta}$ to obtain the maximum likelihood estimate (MLE).

We set a score function [**?**, 119] that takes the derivative of the log-likelihood from Eq. 31 in the main paper and sets it to zero:

$$\frac{\partial}{\partial\boldsymbol{\theta}}\log\mathcal{L}(\mathcal{D}|\boldsymbol{\theta}) = \sum_{i=1}^{N_c} \frac{\partial}{\partial\boldsymbol{\theta}}\log p(d_c, t, \boldsymbol{\theta}) = 0. \tag{3}$$

The score can be solved using the chain rule so that:

$$\sum_{i=1}^{N_c} \frac{1}{p(d_c, t, \boldsymbol{\theta})} \cdot \frac{\partial}{\partial\boldsymbol{\theta}} p(d_c, t, \boldsymbol{\theta}) = 0. \tag{4}$$

Computing an argmax, including an MLE ($\hat{\boldsymbol{\theta}}$), is an optimization problem and can thus be accomplished using standard optimization routines. However, although that may sound simple in theory, solving the optimization of the likelihood function is not uncomplicated. For one thing, the likelihood function itself may be difficult to write down, inefficient to compute, may produce a complex landscape riddled with local maxima - thus confounding attempts to return $\hat{\boldsymbol{\theta}}$, which is meant to represent the global maximum. [72] Such a likelihood landscape becomes increasing likely if the model has too many parameters for the amount of information obtainable from the data, leading to *parameter unidentifiability* [**?**, **?**, 94].

Fig. S8 shows MLE results for the Bursting Gene and simplified STL1 models for the same number of iterations as performed for the 4-state STL1 model fit shown in Example Box 18 in the main text.

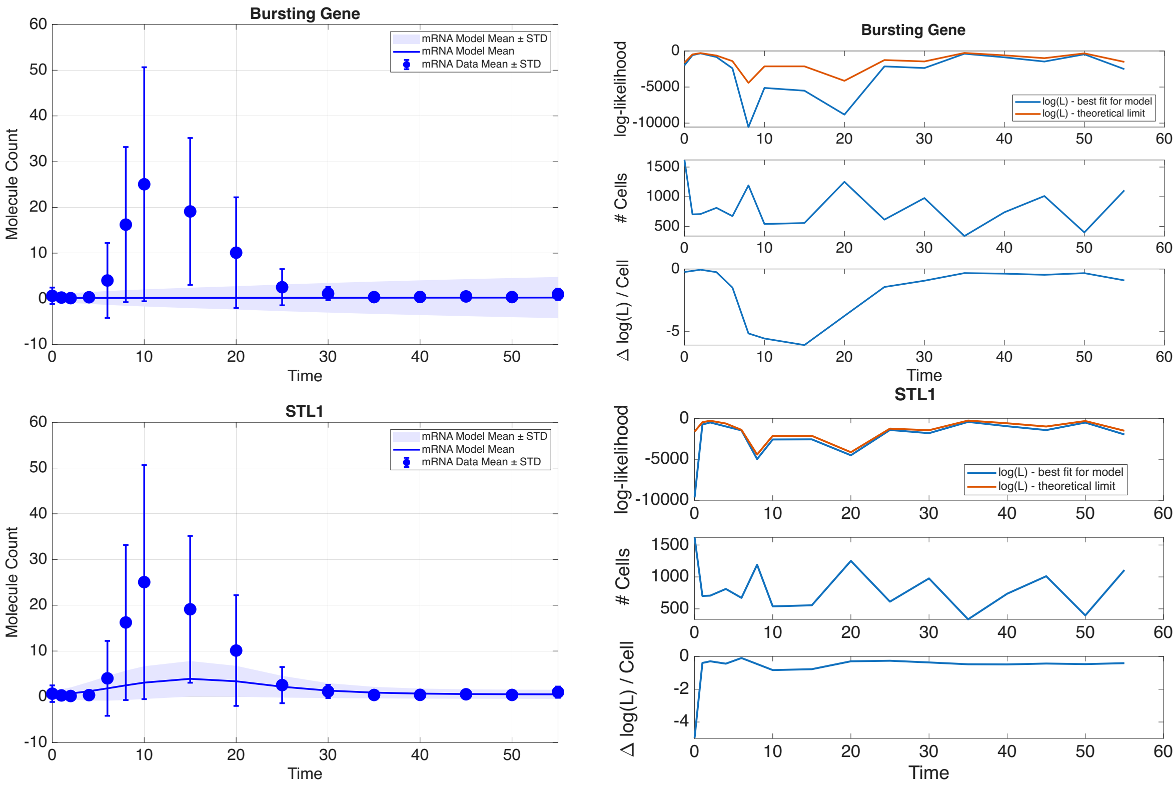


Figure S8: **Maximum likelihood estimation** result plots for (top) the Bursting Gene model and (bottom) the simplified STL1 model, with (left) model vs. data means and standard deviations and (right) log-likelihoods through time.

### 4.2 Bayesian Inference

The prior $\Pr(\boldsymbol{\theta})$ is a distribution of continuous model parameter values that represent the shape of values among which we think the "true" parameters should or might lie based on information we have. In cases where we have minimal information this could be a so-called *uninformative prior* such as a uniform distribution, although the intentional use of such priors is sometimes tritely referred to as the type of prior one uses under the assumption that we have been born a second before being presented with the data - the point being that, with minimal effort, we can usually uncover information about our system from previous experiments or theoretical results and then it would be responsible of us to both find and incorporate this information explicitly in our current statistical analyses [**?**, 123]. Any data that strongly contradicts our prior will weigh in via the likelihood.

To simplify computing the conditional probability of the a set of parameters given data $\Pr(\boldsymbol{\theta}|\mathcal{D})$, known as the *posterior probability* or *posterior probability distribution*, the marginal likelihood (also called the model evidence an prior predictive) $\Pr(\mathcal{D})$ is often omitted and proportional probabilities reported instead, so that:

$$\Pr(\boldsymbol{\theta}|D) \propto \mathcal{L}(\mathcal{D}|\boldsymbol{\theta}) \cdot \Pr(\boldsymbol{\theta}). \tag{5}$$

The $\Pr(\mathcal{D})$ term covers the total probability for all possible occurrences of the data, which may be difficult to obtain and also may not be strictly necessary to answer the researcher's questions. The proportional posterior $\Pr(\boldsymbol{\theta}|\mathcal{D})$ is adequate for the purpose of comparing finite hypotheses, e.g., $\Pr(H_1|D)$ vs. $\Pr(H_2|D)$ where $H_1$ represents one set of parameter values and $H_2$ represents a different set [?, 65].

Computing Eq. 5 can be tricky since the proportionality factor is not always known; with continuous parameters a probability distribution must be computed such that it integrates to one - i.e., over all parameter values $\boldsymbol{\theta}$ in the set $\Theta$:

$$\Pr(D) = \int_{\Theta} \Pr(\mathcal{D}|\boldsymbol{\theta}) \cdot \Pr(\boldsymbol{\theta}) d\theta, \tag{6}$$

Fortunately, it is often the case that most of the probability is contained within a finite, accessible area of the total space and thus can be well sampled using clever algorithms [122, 123].

#### 4.3 Markov Chain Monte Carlo (MCMC)

The maximum likelihood estimate discussed in a previous subsection provides a point estimate - i.e., one set of model parameter values $\boldsymbol{\theta}$ that are particularly likely but may differ from the "true" parameters. The MLE elucidates the height of the probability distribution but does not itself quantify any measure of uncertainty surrounding this estimate. To capture the uncertainty, we use the *Metropolis-Hastings* algorithm (MHA) [120, 121] to sample the target posterior probability distribution from Bayes' theorem [125, 126], Eq. ??/5.

As mentioned in the previous subsection, the MHA falls under the class of numerical methods known as *Markov chain Monte Carlo* (MCMC). MCMC draws samples from its stationary distribution, $\pi(X)$, using a *Markov chain* to approximate a target distribution, commonly the proportional posterior probability distribution for Bayesian inference. MCMC then estimates expectations by *Monte Carlo* integration, a numerical method for computing a definite integral by averaging over randomly selected points at which the integrands are computed.

The *Markov chain* is a memoryless random walk through a sequence of random variables, $X_0, X_1, X_2, \ldots$, that are sampled through a stochastic process that moves from the current state $X_n$ to the next value $X_{n+1}$ according to a transition distribution $q(X_{n+1}|X_n)$. The next state $X_{n+1}$ is only dependent on $X_n$ and thus the Markov chain only on its initial distribution over $X_0$. See Fig. S9.

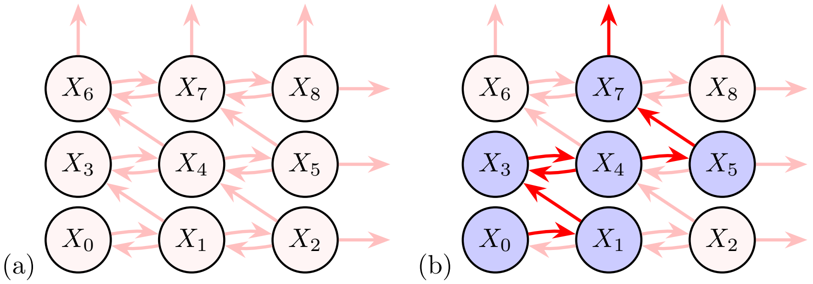


Figure S9: (a) Possible values $\{X_0, X_1, X_2, X_3, X_4, X_5, X_6, X_7, X_8, ...\}$ able to be sampled by a Markov chain; and (b) the random path, shown in red, through states $\{X_0, X_1, X_2, X_3, X_4, X_5, X_6, X_7, X_8, ...\}$ taken by one iteration of a Markov chain. This particular chain includes one backtrack from $X_4$ to $X_3$ so that the full sequence of the path is: $\{X_0, X_1, X_3, X_4, X_3, X_4, X_5, X_7, ...\}$.

From an initial distribution $p(X_0)$, the stationary distribution of the Markov chain $\pi(X)$ can be approached asymptotically by computing the distributions $p(X_i)$ for all times $i$ in any given Markov chain. Convergence occurs when these distributions $p(X_0)$, $p(X_1)$, $p(X_2)$, ... become very similar to one another and can be reached within reasonable run time. Quantitative measures to check for convergence include autocorrelation, effective sample size (ESS), and the Gelman–Rubin diagnostic.

In short, we use MCMC to sample our posterior $\Pr(\boldsymbol{\theta}|\mathcal{D}) \propto \mathcal{L}(\mathcal{D}) \cdot \Pr(\boldsymbol{\theta}) \sim \pi(X)$ by drawing samples from $\pi(X)$ according to the basic procedure:

(i) Choose an initial distribution $p(X_0)$

(ii) Compute transition probabilities $q(X_{n+1}|X_n)$

(iii) Allow chain to converge to equilibrium (after burn-in period, the length of which depends on $p(X_0)$)

(iv) Use converged samples $p(X_i)$ to approximate expectations.

MCMC thereby generates correlated samples from complicated distributions. However, the method for generating the transition probabilities $q(X_{n+1}|X_n)$ must be carefully selected to ensure a reasonable degree of computational efficiency, and herein lies the role of the MHA.

#### 4.4 Metropolis-Hastings

The *Metropolis-Hastings algorithm* (MHA) updates the *Markov chain* discussed in the previous subsection by iteratively and proportionally drawing a new state $\boldsymbol{\theta'}$ from a proposal distribution $f(\theta'|\theta_t)$ given the current state $\theta_t$ and probabilistically accepting or rejecting $\theta'$ according to

$$\alpha(\theta', \theta_t) = \min\left(1, \frac{\Pr(\theta'|\mathcal{D})f(\theta_t|\theta')}{\Pr(\theta_t|\mathcal{D})f(\theta'|\theta_t)}\right) \tag{7}$$

and a randomly drawn number (e. g., $u \sim \text{Uniform}(0,1)$). The proposed state $\theta'$ becomes the new state $\theta_{t+1} = \theta'$ if $u \leq \alpha(\theta', \theta_t)$; otherwise, the proposed state $\theta'$ is rejected in favor of maintaining the current state, $\theta_t = \theta_{t+1}$. The process is repeated for $t = 1, 2, \ldots$ to generate samples from the target distribution $\pi(X)$.

In other words, the MHA follows the steps:

(i) Propose new state $\theta'$ from a proposal distribution $q(\theta'|\theta)$.

(ii) Accept $\theta'$ with probability $\alpha(\theta', \theta_t)$ from Eq. 7.

(iii) If $\alpha(\theta', \theta_t)$ is lower than a defined threshold, reject $\theta'$ and stay at state $\theta$. Otherwise, accept and update the state so that $\theta = \theta'$.

(iv) Repeat ↻ until convergence is reached.

Below, we provide an example for using the FIM for proposals in the sampling of parameter uncertainty / posterior distributions. Table 6 shows the MH posterior sample means and standard deviations.

```
%% Model: FIM inverse
% The inverse of the FIM provides an estimate of the model uncertainty.  Here, we look at the FIM for the log
% of the model parameters and use that to compute the covariance of the log of the parameters.  (Because
% parameters are positive values, but can very significantly in their magnitudes, it is often useful to
% examine them in a log-scale).
Model_FIMlog = diag([Model_MH.parameters{:,2}]) * Model_fimTotal{1} * diag([Model_MH.parameters{:,2}]);
Model_covLog = Model_FIMlog^-1;

% The eigenvalues of covLog tells us what to expect for the uncertainty in the parameters.
[Model_eigVec,Model_eigVal] = eig(Model_covLog);
Model_eigVal = diag(Model_eigVal)

% Here, we see there is one large direction of uncertainty, but the rest are pretty well constrained.
% The direction of the greatest uncertainty is:
[~,j] = max(Model_eigVal);
Model_largestEigVec = Model_eigVec(:,j)

%% Model: Metropolis Hastings
% Now that we have an estimate of the shape of the uncertainty using the FIM we can now search parameter space
% and see what other parameter combinations are also closely matching to our data.  For this, we are going to
% use the Metropolis Hastings algorithm, where the proposal distribution is a multi-variate gaussian with a
% covariance that is proportional to the inverse FIM.

% As the FIM has some very small eigenvalues, we better may be better off reducing the step size in those
% directions.  Here, we set it to at most one order of magnitude by adding an identity matrix to the
% FIM before inverting.
Model_covLogMod = (Model_FIMlog+1*diag(size(Model_FIMlog,1)))^(-1);
```

```
% Here, we set up the MH parameters:
% Set solutions scheme to FSP Sensitivity
Model_MH.solutionScheme = 'FSP';
Model_MH.fittingOptions.modelVarsToFit = 1:4;
Model_MHOptions = struct('numberOfSamples', 1000, 'burnin', 100, 'thin', 3);
proposalWidthScale = 0.001;
Model_MHOptions.proposalDistribution  = @(x)mvnrnd(x,proposalWidthScale * (Model_covLogMod + Model_covLogMod')/2);

% Next, we call the codes to sample the posterior parameter space:
[Modelpars,Model_likelihood,Model_chainResults] = ...
Model_MH.maximizeLikelihood([],Model_MHOptions,'MetropolisHastings');

% Note:  When this runs, you want to see an acceptance of about 0.3 to 0.4, meaning that about a third
% of the proposals are accepted.  If the number is too small you need to decrease the proposal width;
% if it is too large you may need to increase the proposal width. For the default data set and model,
% We find that a scale of .5 to 5% of the FIM-based COV led to an okay acceptance rate, but this is
% variable and will change depending on the initial value in the chain.

% And now to plot the MH results and compare to the FIM.
Model_MH.plotMHResults(Model_chainResults,Model_FIMlog);

% Often the MH search can reveal a better parameter set, so ensure we update our model if it does:
Model_MH.parameters(:,2) = num2cell(Modelpars);
Model_MH.makeFitPlot
% If you do notice better fits, it would be good to re-run the 'fminsearch' again - it is possible to
% find a better model to explain your data.  This can take several rounds of iteration before convergence.
% Let's create a while loop to make it automated.

%% Model: Iterating between MLE and MH
% Run a few rounds of MLE and MH to see if we can get better convergence.
Model_MH.parameters(:,2) = num2cell(Modelpars);
for i=1:3
    % Maximize likelihood
    Modelpars = Model_MH.maximizeLikelihood([],fitOptions);
    % Update parameters in the model:
    Model_MH.parameters(:,2) = num2cell(Modelpars);

    % Compute FIM
    % Set solutions scheme to FSP Sensitivity
    Model_MH.solutionScheme = 'fspSens';
    % Solve the sensitivity problem
    [Model_sensSoln] = Model_MH.solve;
    Model_fimResults = Model_MH.computeFIM(Model_sensSoln.sens,'log');
    Model_FIMlog = Model_FIM.evaluateExperiment(Model_fimResults, Model_MH.dataSet.nCells);

    % Run Metropolis-Hastings
    Model_covLogMod = (Model_FIMlog{1} + diag(size(Model_FIMlog{1},1)))^(-1); % Adjusted proposal
    proposalWidthScale = 0.0001;                                             % Distribution covariance
    Model_MHOptions.proposalDistribution  = @(x)mvnrnd(x,proposalWidthScale * (Model_covLogMod+Model_covLogMod')/2);
    [Modelpars,Model_likelihood,Model_chainResults] = ...
    Model_MH.maximizeLikelihood([], Model_MHOptions, 'MetropolisHastings');
    % Update parameters in the model:
    Model_MH.parameters(:,2) = num2cell(Modelpars);
end
Model_MH.plotMHResults(Model_chainResults,Model_FIMlog);
Model_MH.makeFitPlot

%% Model: Evaluating the MH results
% Here we will generate three plots.  The first one will show the likelihood function as we search over
% parameter space.  In order to get a good estimate of the parameter uncertainty, we want this to quickly
% reach the maximum value and then to fluctuate around that value for a significant amount of time.
% If you see that it is still increasing, you know that the fit has not yet converged.
figure
subplot(1,3,1)
plot(Model_chainResults.mhValue)
title('MH Convergence')
xlabel('Iteration Number')
```

```
ylabel('LogLikelihood')

%% Compute FIM
% Set solutions scheme to FSP Sensitivity
Model_MH.solutionScheme = 'fspSens';
% Solve the sensitivity problem
[Model_sensSoln] = Model_MH.solve;
Model_fimResultsLog = Model_MH.computeFIM(Model_sensSoln.sens,'log');
Model_FIMlog = Model_MH.evaluateExperiment(Model_fimResultsLog, Model_MH.dataSet.nCells);
Model_fimResults= Model_MH.computeFIM(Model_sensSoln.sens,'lin');
Model_FIM = Model_MH.evaluateExperiment(Model_fimResults, Model_MH.dataSet.nCells);

% Next, we will show the scatter plot of a couple parameters.  It is helpful to show these in linear scale
% as well as in a natural log scale.  For illustration, we also compare the spread of the posterior to the
% covariance predicted by the FIM from before.

% Choose which parameters to compare.
Q = [3,4];
subplot(1,3,2)

% Plot uncertainty in linear scale
Model_MH.makeMleFimPlot(exp(Model_chainResults.mhSamples)', Model_FIM{1},Q,0.95,1); hold on
title('Posterior -- Linear Scale')
xlabel(Model_MH.parameters{Q(1),1})
ylabel(Model_MH.parameters{Q(2),1})

% Plot uncertainty in log scale
subplot(1,3,3)
Model_MH.makeMleFimPlot(Model_chainResults.mhSamples', Model_FIMlog{1},Q,0.95,1); hold on
title('Posterior -- Natural Log Scale')
xlabel(['log',Model_MH.parameters{Q(1),1}])
ylabel(['log',Model_MH.parameters{Q(2),1}])

%% Model: Effective Sample Size
% For the MH, it is important to get a sense of how well it has sampled the posterior.  For this, we
% determine the effective sample size (i.e., the number of effectively indpendent samples within the MH chain).
% This is found by examining the autocorrelation of the parameter chain to figure out the number of steps
% needed for correlations to decay and then divide the total number of steps by the de-correlation step.
figure
ipar = 4;
ac = xcorr(Model_chainResults.mhSamples(:,ipar) - mean(Model_chainResults.mhSamples(:,ipar)),'normalized');
ac = ac(size(Model_chainResults.mhSamples,1):end);
plot(ac,'LineWidth',3)
N = size(Model_chainResults.mhSamples,1);
tau = 1+2*sum(abs(ac(2:N/5)));
Neff = N/tau
```

#### 4.5 Approximate Bayesian Computation (ABC)

Unlike full Bayesian inference which obtains the posterior distribution $\Pr(\theta|\mathcal{D})$ using the likelihood term $\mathcal{L}(\mathcal{D}|\theta)$, ABC approaches simulate data $\hat{\mathcal{D}}_\theta$ [?, 63] by:

(i) Drawing values from the prior distribution $\Pr(\theta)$ to obtain a sample parameter vector $\hat{\theta}$;

(ii) Applying $\hat{\theta}$ to our reaction model to generate simulated data $\hat{\mathcal{D}}_\theta$ (e.g., SSA trajectories or FSP solutions);

(iii) Comparing the simulated data $\hat{\mathcal{D}}_\theta$ to real, experimental data $\mathcal{D}$ using a chosen metric to define the distance between the simulated data and real experimental data or based on summary statistics;

(iv) Accepting or rejecting $\hat{\theta}$ for inclusion in the approximation of $\Pr(\theta|\mathcal{D})$ depending on comparison of some threshold $\epsilon$ and the distance between the simulated data $\hat{\mathcal{D}}_\theta$ and experimental data $\mathcal{D}$, or between the respective summary statistics $\mathcal{S}(\hat{\mathcal{D}}_\theta)$ and $\mathcal{S}(\mathcal{D})$, such that $d(\hat{\mathcal{D}}_\theta, \mathcal{D}) \leq \epsilon$ or $d(\mathcal{S}(\hat{\mathcal{D}}_\theta), \mathcal{S}(\mathcal{D})) \leq \epsilon$.

For single-cell experimental data, a molecule count and time point are drawn from the reaction model and either the full distributions (CDFs of counts across cells) or statistical summaries (means, variances, etc.)

of the simulated data and experimental data are compared [64, 127]. A very simple, unweighted distance metric for $N$ species would be:

$$d = \sum_{i=1}^{N} \mid \mathrm{CDF}_i^{\mathrm{SSA}} - \mathrm{CDF}_i^{\mathrm{Data}} \mid \quad (8)$$

Below, we provide example code for the application of ABC for inferring the parameters for a random gene from scRNA-seq data [68]. Results from the ABC search are shown in Fig. S10.

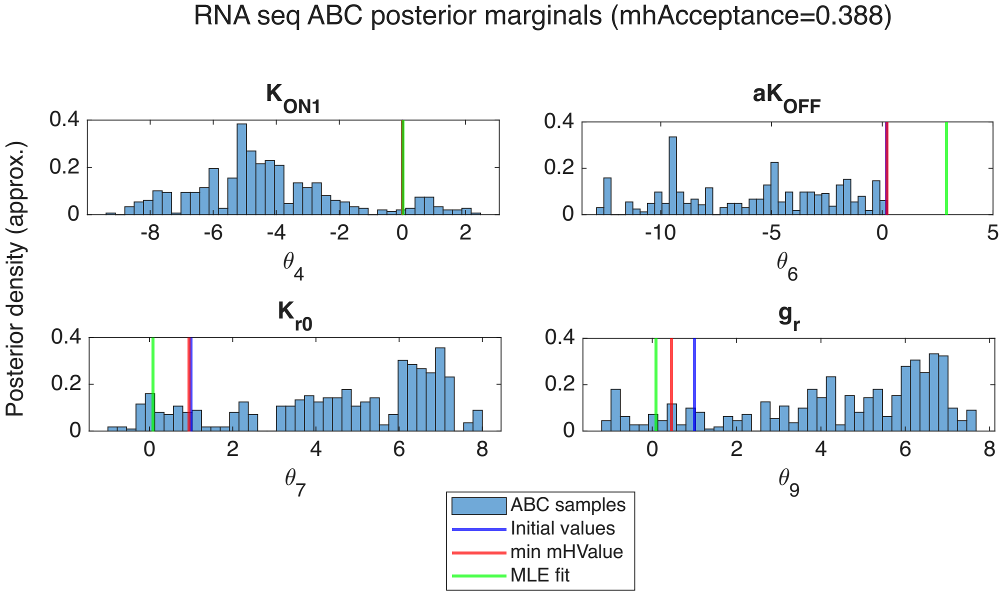


Figure S10: **Approximate Bayesian Computation** results for gene TSC22D3, a randomly selected gene from the 151 genes we fit in the scRNA-seq example scripts ('example_scRNAseq_...') showing the approximate posterior densities for a subset of the fitted parameters. The blue lines show the initial parameter values (given by parGuess), red lines show the min. mh values, and the green lines are the MLE-fitted parameter values from the pipeline in 'example_scRNAseq_2_BatchFitManyGenes.m'.

```
%%%%%%%%%%%%%%%%%%%%%%%%%%%%%%%%%%%%%%%%%%%%%%%%%%%%%%%%%%%%%%%%%%%%%%%%%%%
%% Approximate Bayesian Computation (ABC) in the SSIT using 'runABCsearch'
%
%   1. Load a template model for scRNA-seq genes with SSA solution scheme.
%   2. Associate the template model with scRNA-seq data for gene TSC22D3.
%   3. Define a prior over parameters.
%   4. Run ABC via Metropolis{Hastings using 'cdf_one_norm' loss.
%   5. Visualize the ABC results.
%%%%%%%%%%%%%%%%%%%%%%%%%%%%%%%%%%%%%%%%%%%%%%%%%%%%%%%%%%%%%%%%%%%%%%%%%%%

% Make copy of scRNAseq template model and choose which parameters to fit:
scRNAseq = Model_Template;
fitpars = 1:9;  scRNAseq.fittingOptions.modelVarsToFit = [fitpars];

%% Set SSA options:
scRNAseq.solutionScheme = 'SSA';            %Set solution scheme to SSA
scRNAseq.ssaOptions.nSimsPerExpt=100;       % Set number of simulations performed per experiment (small # for demo)
scRNAseq.tSpan = [-100,scRNAseq.tSpan];     % Equilibrate before starting (burn-in)
scRNAseq.ssaOptions.useParallel = true;     % Run iterations in parallel with multiple cores

%% Associate scRNA-seq data for gene TSC22D3:
scRNAseq = scRNAseq.loadData('data/Raw_DEX_UpRegulatedGenes_ForSSIT.csv', {'rna','TSC22D3'});
```

```
%% Set up a prior over parameters (logPriorLoss)
% logPriorLoss should return a *loss* (positive penalty); smaller is better.
% A convenient choice is a quadratic penalty in log10-parameter space, corresponding to a log-normal prior.
theta0 = cell2mat(scRNAseq.parameters([fitpars],2));
log10_mu = log10(theta0(:));
log10_sigma = 2 * ones(size(log10_mu));                              % std dev in log10-space
fitOptions.logPrior = @(x) -sum((x-log10_mu).^2./(2*log10_sigma.^2));   % log prior

% Define prior "loss" (default, @(x)allFitOptions.obj(exp(x))):
logPriorLoss = [];

%% Set ABC / MCMC options
% runABCsearch passes 'fitOptions' to maximizeLikelihood with the 'MetropolisHastings' algorithm.
% Tune these depending on your problem size.

fitOptions = struct();
fitOptions.numberOfSamples       = 500;          % Total MH iterations (small # for demo)
fitOptions.burnIn                = 10;           % Discard burn-in samples
fitOptions.thin                  = 1;            % Keep every nth sample
proposalWidthScale               = 0.5;          % Proposal scale (acceptance should be approx. 0.3-0.4)
fitOptions.proposalDistribution  = @(x)x+proposalWidthScale*randn(size(x));     % Proposal distribution

% Initial parameter guess (optional, default: current Model.parameters):
% parGuess = [];
parGuess = cell2mat(scRNAseq.parameters((fitpars),2));  % equiv. to 'parGuess = [];' which would use default

% Choose loss function for ABC (default: 'cdf_one_norm'):
lossFunction = 'cdf_one_norm';

% Enforce independence by downsampling SSA trajectories:
enforceIndependence = true;

%% Run ABC search
% This will:
%   * repeatedly simulate SSA trajectories,
%   * compute a CDF-based loss against the data,
%   * add the prior penalty, and
%   * perform MH sampling to approximate the posterior.
% Outputs:
%   pars                    - "best" (minimum-loss) parameter set found
%   minimumLossFunction     - value of the loss at that point
%   Results                 - MH/ABC diagnostics and chains
%   ModelABC                - model updated with 'pars'

% Compile and store the given reaction propensities:
scRNAseq = scRNAseq.formPropensitiesGeneral('scRNAseq');

% Run ABC search:
[parsABC, minimumLoss, ResultsABC, scRNAseq] = scRNAseq.runABCsearch(parGuess, lossFunction, logPriorLoss,...
                                                                     fitOptions, enforceIndependence);

%% Inspect ABC results:
fprintf('ABC completed.\n');            fprintf('Minimum loss value: %g\n', minimumLoss);
disp('Best-fit parameters (ABC):');     disp(parsABC(:).');

% The 'ResultsABC' struct is returned by maximizeLikelihood with the
% 'MetropolisHastings' algorithm.
%       ResultsABC.mhSamples        - MCMC chain of parameter samples
%       ResultsABC.mhValue          - corresponding loss values
%       ResultsABC.mhAcceptance     - MH acceptance fraction
% Below we show a simple marginal histogram for each fitted parameter.

% Plot results:
if isfield(ResultsABC, 'mhSamples')
    parChain = ResultsABC.mhSamples;   % size: [numberOfSamples x nPars]
    nPars    = size(parChain, 2);
    figure;
    for k = 1:nPars
```

```
        subplot(ceil(nPars/2), 2, k);
        histogram(parChain(:,k), 40, 'Normalization', 'pdf');
        hold on;
        xline(parGuess(k), 'b', 'LineWidth', 1.5);
        xline(parsABC(k), 'r', 'LineWidth', 1.5);
        xline(cell2mat(Model_TSC22D3.parameters(k,2)), 'g', 'LineWidth',1.5);
        title(sprintf('Parameter %d', k));
        xlabel('\theta_k');
        ylabel('Posterior density (approx.)');
    end
    sgtitle('ABC posterior marginals (approximate)');
else
    warning('ResultsABC.mhSamples not found.');
end

%% Compare initial vs. final (ABC) parameter losses (experimental data vs. data simulated by the SSA):
% Minimum loss from ABC run:
minLoss = minimumLoss;
nTimes  = sum(scRNAseq.fittingOptions.timesToFit);
% Set the number of species being fitting (in this case, only RNA):
nSpecies = 1;
% Replicate groups / dose groups etc., if applicable:
nConds   = 1;
avgLossPerCDF = minLoss / (nTimes * nSpecies * nConds);
fprintf('Average CDF L1 discrepancy per time/species: %.4f\n',avgLossPerCDF);
L_init = scRNAseq.computeLossFunctionSSA(lossFunction, theta0, enforceIndependence);
L_min  = minimumLoss;
fprintf('Initial loss: %.3f,  Final (min) loss: %.3f\n', L_init, L_min);
fprintf('Relative improvement: %.1f%%\n', 100 * (L_init - L_min)/L_init);

%% Compare ABC posterior sample to MLE fit:
% load('seqModels/Model_TSC22D3.mat')    % If not in memory, load MLE-fitted parameters for gene TSC22D3
theta_TSC22D3 = cell2mat(Model_TSC22D3.parameters(1:9,2));  % Get MLE-fitted parameters for gene TSC22D3
L_MLE = scRNAseq.computeLossFunctionSSA(lossFunction, theta_TSC22D3, enforceIndependence);
L_min  = minimumLoss;
fprintf('MLE loss: %.3f,  Final (min) ABC loss: %.3f\n', L_MLE, L_min);
fprintf('Relative improvement: %.1f%%\n', 100 * (L_MLE - L_min)/L_MLE);
```

#### 4.6 Cross-validation

Cross-validation is commonly used to compare the abilities of different models or machine learning methods to learn the model parameters that best predict test data based on training data by partitioning the data $\mathcal{D}$ into different subsets $d_i \subset \mathcal{D}$, one containing data samples to train a model and one containing samples to test the prediction, and then iteratively resampling from $\{d_1, ...d_n\}$ to estimate how a model will generalize for unseen data [?, ?, ?] (see Fig. S11).

Following the maximum likelihood estimation from the main section, we can independently optimize the log-likelihood $\log(\mathcal{L}(d_i|\theta))$ for each $d_i$, returning

$$\theta_i = \underset{\theta}{\text{argmax}}[\log(\mathcal{L}(d_i|\theta))], \tag{9}$$

and fit each set of $\theta_i$ by computing the log-likelihood $\log(\mathcal{L}_\rangle)$ for all data,

$$\log(\mathcal{L}_\rangle) = \log(\mathcal{L}(\mathcal{D}|\theta_i)) \leq \log(\mathcal{L}_{\text{Fit}}). \tag{10}$$

This technique can thus be used to quantify the amount of parameter uncertainty by comparing the predictions of models computed based on the training data set to the model fits on the test data, as in [25], with the cross-validation (CV) error computed as the average,

$$\log(\mathcal{L}_{\text{CV}}) = \frac{1}{n}\sum_{i=1}^{n}\log(\mathcal{L}(\mathcal{D}|\theta_i)) \leq \log(\mathcal{L}_{\text{Fit}}). \tag{11}$$

Below, we provide an example for computing replica-to-replica variation using cross-validation in the SSIT for the 4-state STL1 model discussed in the main paper (Fig B1). The results are shown in Fig. S12.

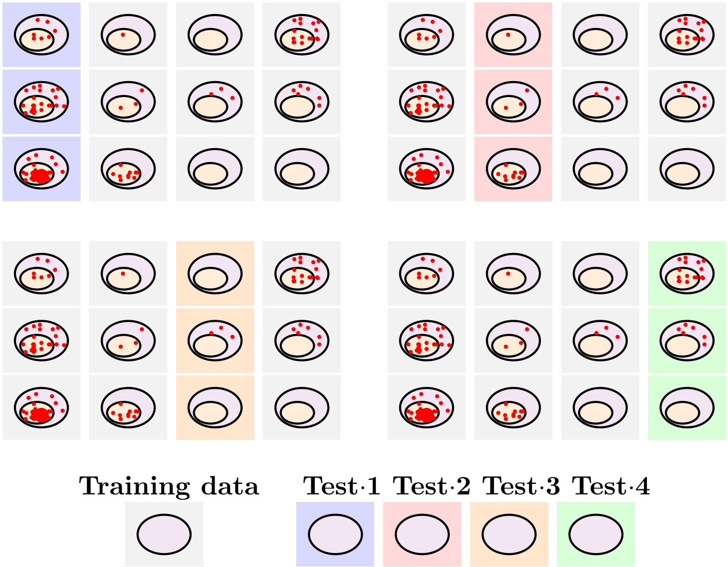


Figure S11: Cross-validation tests the ability of models to predict data by repetitively dividing a data set into training data, on which to train the model, and test data, used to test model predictions.

```
%%%%%%%%%%%%%%%%%%%%%%%%%%%%%%%%%%%%%%%%%%%%%%%%%%%%%%%%%%%%%%%%%%%%%%%%%%%%%%%%%%%%%%%%%%%%%%%%%%%%%%%%%%%%%%%%
%% Perform Cross-Validation fitting different replicas at the same time using SSITMultiModel
%  * In this example, we use the SSITMultiModel to allow parameters to change for different replica data sets
% (e.g., to allow for batch variations, or explore how parameters change under different genetic variations).
%%%%%%%%%%%%%%%%%%%%%%%%%%%%%%%%%%%%%%%%%%%%%%%%%%%%%%%%%%%%%%%%%%%%%%%%%%%%%%%%%%%%%%%%%%%%%%%%%%%%%%%%%%%%%%%%

%% Set Fitting Options:
fitAlgorithm = ‘fminsearch’;    fitOptions = optimset(‘Display’,‘final’,‘MaxIter’,200);
% Note: ‘MaxIter’, 200 for fast run; Set to ‘MaxIter’, 2000 for accuracy

% Make a copy of our 4-state STL1 model:
STL1_4state_CrossVal = STL1_4state_MH;

% Specify datafile name and species linking rules:
DataFileName = ‘data/filtered_data_2M_NaCl_Step.csv’;   LinkedSpecies = {‘mRNA’,‘RNA_STL1_total_TS3Full’};

% Suppose we only wish to fit the data at times before 25 minutes.
ConditionsGlobal = {[], [], ‘TAB.time<=25’};

% Split up the replicas to be separate:
ConditionsReplicas = {‘TAB.Replica==1’; ‘TAB.Replica==2’};

% Specify constraints on rep-to-rep parameter variations. Here, we specify that there is an expected 0.1
% log10 deviation expected in some parameters and smaller in others.  No deviation at all is indicated by 0.
Log10Constraints = [0.1,0.1,0.1,0.1,0.1,0.1,0.1,0.1,0.02,0.02,0.02,0.02,0.02];

% Create full model:
CrossValidationModel = SSITMultiModel.createCrossValMultiModel(STL1_4state_CrossVal, DataFileName,...
                        LinkedSpecies, ConditionsGlobal, ConditionsReplicas, Log10Constraints);
CrossValidationModel = CrossValidationModel.initializeStateSpaces;

% Run the model fitting routines:
crossValPars = CrossValidationModel.parameters;
crossValPars = CrossValidationModel.maximizeLikelihood(crossValPars, fitOptions, fitAlgorithm);
CrossValidationModel = CrossValidationModel.updateModels(crossValPars);
CrossValidationModel.parameters = crossValPars;

% Make a figure to explore how much the parameters changed between replicas:
fignum = 11;  useRelative = true;  CrossValidationModel.compareParameters(fignum,useRelative);
```

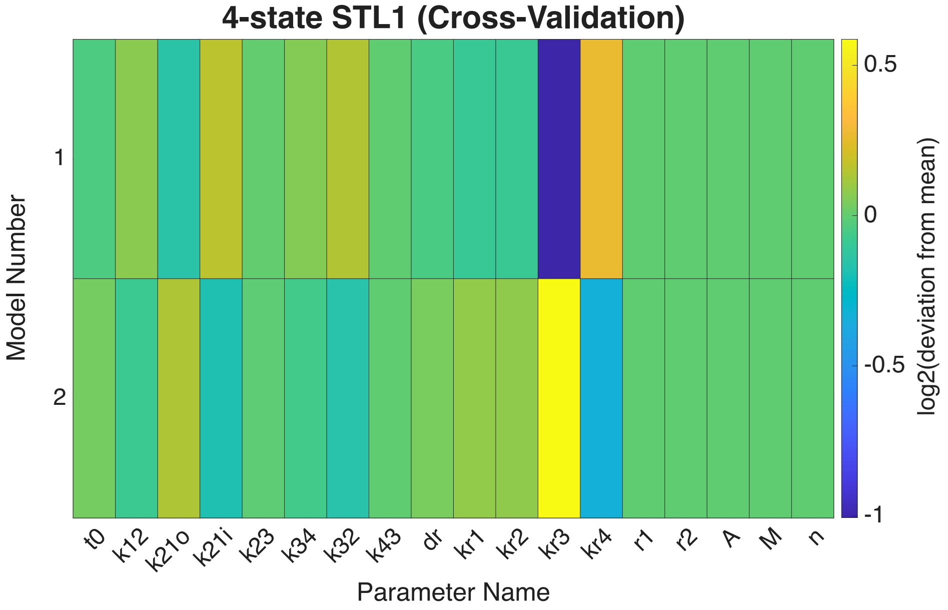


Figure S12: **Cross-validation** example showing a replica-to-replica comparison of the deviation from the mean across all parameters of the 4-state STL1 model from the main paper (Fig B1).

### 5 Complex models

The SSIT has capabilities for handling several different types of complexity in solving reaction models, including: model reduction techniques for computational efficiency, which itself includes hybrid deterministic-stochastic solutions; fitting multiple datasets and/or models which share some or all parameters; and transformation of probability distributions that have been affected by distorted data (using probabilistic distortion operators, or PDOs [30]). Below, we elaborate on these techniques.

#### 5.1 Basic model reduction strategies

In this subsection, we add details for the use of different model reduction strategies in the SSIT. The user specifies the model reduction tactic they wish to engage with 'reductionType', e.g.:

```
reductionType = 'POD';
```

Below, we briefly explain model reduction techniques that are currently available in SSIT, though as SSIT is continually developed there will likely be additional methods available in short time.

- **Finite State Projection:** To set the FSP tolerance in the SSIT, simply specify:

```
Model.fspOptions.fspTol = 1e-4;
```

- **Hybrid Reduction Methods:** In the SSIT user specification may look something like:

```
Model.useHybrid = true;
Model.hybridOptions.upstreamODEs = {'g1','g2','g3','g4'};
Model.solutionScheme = 'FSP';
```

  where four model species, 'g1', 'g2', 'g3', and 'g4' are handled using ODEs and the remaining model species will be solved by FSP. Any FSP arguments may be made as normal.

- **Quasi-Steady-State Approximation (QSSA):** In the SSIT, the list of species to be assumed at QSSA must be specified in a vector 'reductionSpecies', like so:

```
qssaSpecies = 2;
reductionSpecies = {'g1','g2','g3','g4'};
```

- **Log Lump QSSA:** In the SSIT, the number of grid lines of the log lump QSSA must be specified using 'reductionOrder':

```
reductionOrder = 20;
```

- **Eigen Decomposition Initial:** Eigen decomposition can be used to consider only the space spanned by the initial condition plus the eigvenvectors corresponding to the eigenvalues with the largest real values. This reduction works well when a few eigenmodes dominate the system dynamics due to the initial condition. The user specifies the number of modes to consider in the reduction with 'reductionOrder'. For time-varying systems, the basis vectors are found using the infinitesimal generator at t=0.
- **Proper Orthogonal Decomposition (POD):** Model reduction by POD is best employed using a fine time resolution in the calculation of the FSP. In the SSIT, the user must specify the size of the reduced model as 'reductionOrder':

```
reductionOrder = 20;
```

Below, we provide example code for performing the 'Proper Orthogonal Decomposition' model reduction on our 4-state STL1 model (Fig B1 from main text). One thing to note is that because for this method we need to generate a basis set using solutions at finer resolution, POD will be inefficient for the initial set up of the reduction. The benefits typically come from solving the model multiple times with different parameter sets. In the case of our 4-state STL1 model, an example run gave the following solution times for the full FSP and reduced FSP:

```
STL1_SolveTime = 2.5360e-01
STL1_SolveTimeReduced = 7.4901e-02
```

Fig. S13 shows the results from the POD example.

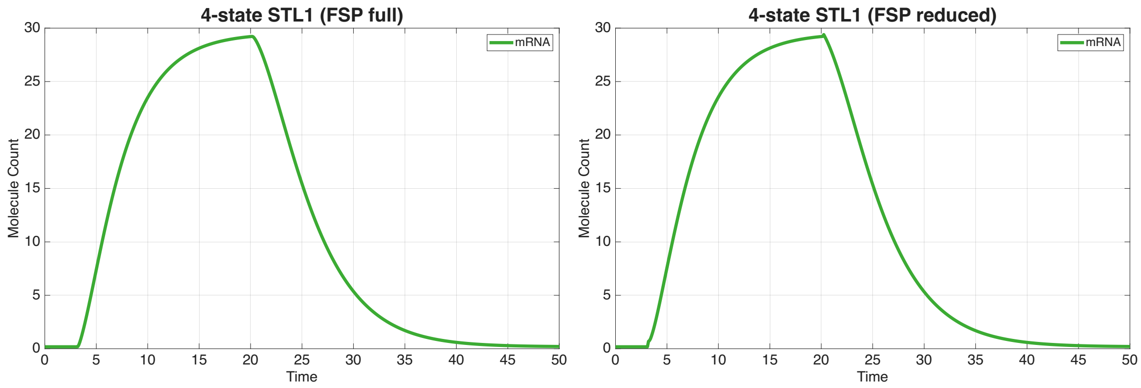


Figure S13: **Model reduction** by proper orthogonal decomposition: (left) mRNA counts through time from the full FSP solution, versus (right) mRNA counts according to the POD reduction. Care should be taken when selecting POD options as additional errors appear in reduced solutions; these errors are not qualitatively noticeable for 'podTimeSetSize=30' and 'reductionOrder = 50'.

```
% Make a copy of the STL1 model to set up for model reduction:
STL1_MR_setup = STL1_4state_FSP;

reductionType = 'POD';
reductionOrder = 50;
podTimeSetSize = 30;
```

```
% Set up to solve FSP solution again to time following expansion:
STL1_MR_setup.fspOptions.initApproxSS = true; STL1_MR_setup.tSpan = linspace(0,50,30);

% Print the computation time to solve the FSP using "tic" and "toc":
tic
[~,~,STL1_MR_setup] = STL1_MR_setup.solve;
STL1_SolveTime = toc

% Turn off further FSP expansion:
STL1_MR_setup.fspOptions.fspTol = inf;

%% Solve the POD reduced model for STL1:
% Make a copy of the full model:
STL1_MR = STL1_MR_setup;

% Set the solver to use ModelReduction:
STL1_MR.modelReductionOptions.useModReduction = true;
% FSP expansion should be suppressed when using Model Reductions

% Set type and order of Model Reduction:
STL1_MR.modelReductionOptions.reductionType = reductionType;
STL1_MR.modelReductionOptions.reductionOrder = reductionOrder;

% Call the SSIT to compute the model reduction transformation matrices:
STL1_MR = STL1_MR.computeModelReductionTransformMatrices;

% Solve the reduced model:
tic
[~,~,STL1_MR] = STL1_MR.solve;
STL1_SolveTimeReduced = toc

%% Plot the full and reduced FSP solutions:
STL1_MR_setup.plotFSP([],STL1_MR_setup.species(5), 'means', [], [], {'linewidth',4},...
    Title='4-state STL1 (FSP full)', TitleFontSize=24, AxisLabelSize=18,...
    TickLabelSize=18, XLabel='Time', YLabel='Molecule Count',...
    LegendFontSize=15, LegendLocation='northeast', Colors=[0.23,0.67,0.2],...
    YLim = [0,32]);

STL1_MR.plotFSP([],STL1_MR.species(5), 'means', [], [], {'linewidth',4}, XLabel='Time',...
    YLabel='Molecule Count', Title='4-state STL1 (FSP reduced)',...
    TitleFontSize=24, AxisLabelSize=18, TickLabelSize=18,...
    Colors=[0.23,0.67,0.2], LegendFontSize=15, LegendLocation='northeast',...
    YLim = [0,32]);
```

#### 5.2 Hybrid deterministic-stochastic solutions

Example code for telling the SSIT to compute deterministic solutions for certain upstream model species, such as the four gene states in our 4-state STL1 model (Fig B1 in the main text), but compute full FSP solutions for downstream species, such as mRNA, is provided below. Fig. S14 shows the solutions for mRNA.

```
% Set 'useHybrid' to true:
STL1_hybrid.useHybrid = true;

% Define which species will be solved by ODEs:
STL1_hybrid.hybridOptions.upstreamODEs = {'g1','g2','g3','g4'};

% Set solution scheme to FSP:
STL1_hybrid.solutionScheme = 'FSP';

% Set FSP 1-norm error tolerance:
STL1_hybrid.fspOptions.fspTol = 1e-4;

% Guess initial bounds on FSP StateSpace:
STL1_hybrid.fspOptions.bounds = [1,1,1,1,300];
```

```
% Compile and store the given reaction propensities:
STL1_hybrid = STL1_hybrid.formPropensitiesGeneral(‘STL1_hybrid’);

% Have FSP approximate the steady state for the initial distribution:
STL1_hybrid.fspOptions.initApproxSS = true;

% Solve STL1_hybrid:
[STL1_hybrid_FSPsoln,STL1_hybrid.fspOptions.bounds] = STL1_hybrid.solve;

% Plot means and standard deviations, then marginal distributions:
STL1_hybrid.plotFSP(STL1_hybrid_FSPsoln, STL1_hybrid.species(5), ‘meansAndDevs’, [], [], {‘linewidth’,4},...
    Colors=[0.23,0.67,0.2], Title=‘4-state STL1 (hybrid)’, TitleFontSize=24, YLim=[-10,40],...
    LegendFontSize=15, LegendLocation=‘northeast’);
STL1_hybrid.plotFSP(STL1_hybrid_FSPsoln, STL1_hybrid.species(5), ‘marginals’, [1,12,24,50,101,200], [],...
    {‘linewidth’,3}, XLim=[0,100], Colors=[0.23,0.67,0.2], AxisLabelSize=18, TickLabelSize=15)
```

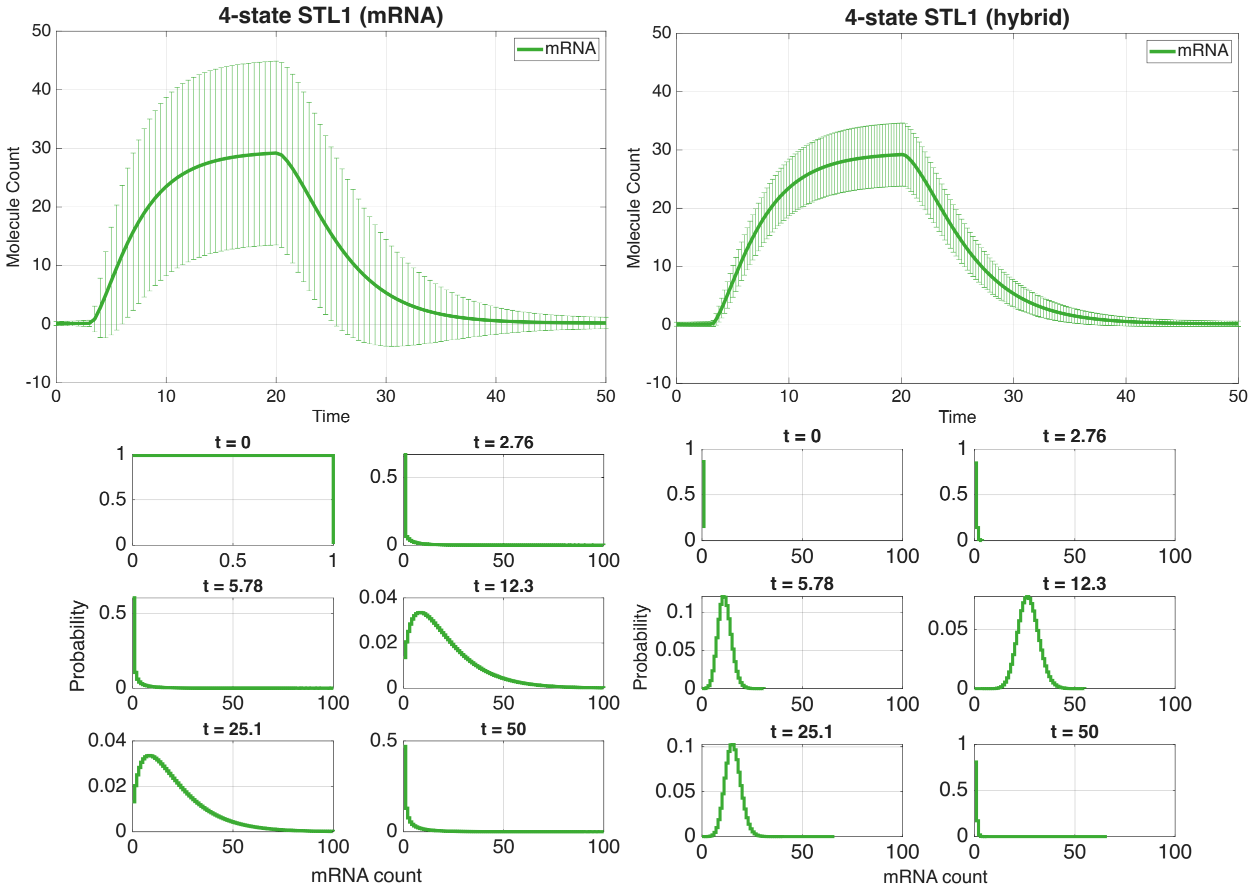


Figure S14: **Hybrid deterministic-stochastic solutions** can be computed by the SSIT for improved computational efficiency. Here, the four gene states of the 4-state STL1 model are treated deterministically, while mRNA is stochastic. The means and standard deviations are shown in the first row and the marginal distributions are shown below, with the FSP solution for mRNA when the (upstream) gene species were also solved using FSP shown on the left and the FSP solution for mRNA from the hybrid solutions (when the four gene species were treated deterministically) shown on the right.

#### 5.3 Probabilistic distortion operators (PDOs)

Below we provide the code for calibrating PDOs for STL1 data [25]. In addition to the binomial PDO for missing spot counts discussed in the main text, we include an affine Poisson PDO for treating average intensity data in lieu of spot counts. The function ‘calibratePDO’ will generate the plots shown in Fig. S15.

```
STL1_4state_PDO_cyt = STL1_4state_PDO_cyt.calibratePDO('data/filtered_data_2M_NaCl_Step.csv',...
    {'mRNA'}, {'RNA_STL1_total_TS3Full'}, {'RNA_STL1_cyto_TS3Full'}, 'Binomial', true, [], {'Replica',1},...
    LegendLocation=''northwest", Title="4-state STL1 (Binomial PDO: Cytoplasmic mRNA)", FontSize=24,...
    XLabel="Total mRNA counts", YLabel="Cytoplasmic mRNA counts");

STL1_4state_PDO_nuc = STL1_4state_PDO_nuc.calibratePDO('data/filtered_data_2M_NaCl_Step.csv',...
    {'mRNA'}, {'RNA_STL1_total_TS3Full'}, {'RNA_STL1_nuc_TS3Full'}, 'Binomial', true, [], {'Replica',1},...
    LegendLocation="northwest", Title="4-state STL1 (Binomial PDO: Nuclear mRNA)", FontSize=24,...
    XLabel="Total mRNA counts", YLabel="Nuclear mRNA counts");

%%%%%%%%%%%%%%%%%%%%%%%%%%%%%%%%%%%%%%%%%%%%%%%%%%%%%%%%%%%%%%%%%%%%%%%%%%%%%%%%%%%%%%%%%%%%
%% Ex(2): Calibrate PDO from average intensity data
% Calibrate the PDO from empirical data. Here, the number of spots has been measured using
% different assays in data column 'RNA_STL1_total_TS3Full' for the 'true' data set (mRNA spot
% counts) and in the columns 'STL1_avg_int_TS3Full' for integrated intensity.  We calibrate
% an 'AffinePoiss' PDO where the obervation probability is a Poisson distribution where the
% mean value is affine linearly related to the true value: P(y|x) = Poiss(a0 + a1*x)
%%%%%%%%%%%%%%%%%%%%%%%%%%%%%%%%%%%%%%%%%%%%%%%%%%%%%%%%%%%%%%%%%%%%%%%%%%%%%%%%%%%%%%%%%%%%
% Make guesses for the PDO hyperparameters λ, in this case: λ1, λ2, λ3
% Model: μ(x) = max(λ1, λ2 + λ3·x)
parGuess = [0, 2500, 5];

STL1_4state_PDO_intens = STL1_4state_PDO;
STL1_4state_PDO_intens = STL1_4state_PDO_intens.calibratePDO( 'data/filtered_data_2M_NaCl_Step.csv',{'mRNA'},...
    {'RNA_STL1_total_TS3Full'}, {'STL1_avg_int_TS3Full'}, 'AffinePoiss', true, parGuess, {'Replica',1},
    LegendLocation="southeast", Title="4-state STL1 (Affine PDO: Average intensity)", FontSize=24,...
    XLabel="True mRNA counts",YLabel="Average intensities (binned)");
```

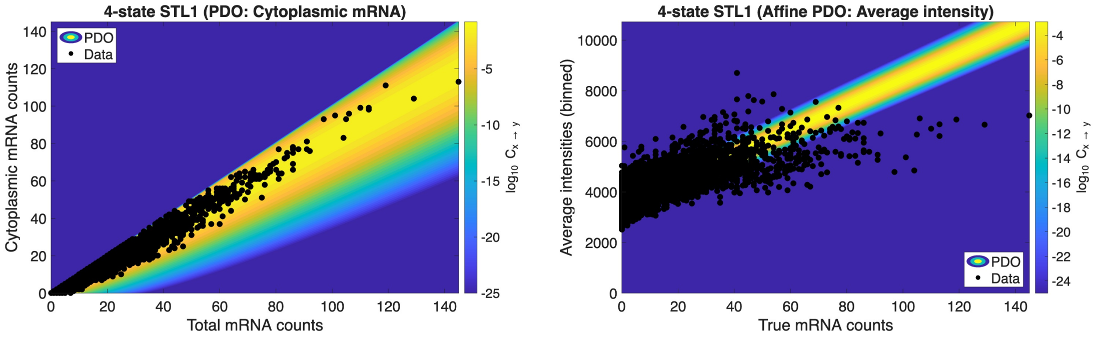


Figure S15: **Probabilistic distortion operators** transform probability distributions computed from distorted data to more closely resemble those computed from truer data (often more costly to obtain). In this case, a simple binomial PDO for "missing spot" counts as described in Huy *et al.* [30] was applied for (left) cytoplasmic mRNA counts, and (right) nuclear mRNA counts.

### 5.4 SSITMultiModel Class for Combinations of Related Models and Data Sets

Here, we provide example code for using SSITMultiModel to fit shared 4-state STL1 model parameters using data from two experimental replicas. In this example, STL1_4state_multi_1 ($1^{st}$ replica) and STL1_4state_multi_2 ($2^{nd}$ replica) share parameters 1-11, but each model infers its own 'kr3' and 'kr4'. These parameters were selected due to having the highest deviation from the mean in the cross validation results (Fig. S12) MultiModel results are shown in Fig. S16. Table 6 shows the properties of SSITMultiModel.

```
% Load and associate STL1 data for both models:
STL1_4state_multi_1 = STL1_4state_multi_1.loadData('data/filtered_data_2M_NaCl_Step.csv',...
                      {'mRNA','RNA_STL1_total_TS3Full'}, {'Replica',1;'Condition','0.2M_NaCl_Step'});

STL1_4state_multi_2 = STL1_4state_multi_2.loadData('data/filtered_data_2M_NaCl_Step.csv',...
```

```
                    {'mRNA','RNA_STL1_total_TS3Full'}, {'Replica',2;'Condition','0.2M_NaCl_Step'});

% Select which models to include in SSITMultiModel:
Models_mix = {STL1_4state_multi_1, STL1_4state_multi_2};

%% Define how parameters are assigned to sub-models by their indices.
% The first 11 parameters are shared and each model has two of its own parameters stored in separate indices:
ParsIndices_mix = {[1:11,12:13],[1:11,14:15]};

% Combine models into one "MultiModel", specify parameters, and initialize:
combinedModelMixed = SSITMultiModel(Models_mix, ParsIndices_mix);
combinedModelMixed = combinedModelMixed.initializeStateSpaces;

% Store parameters for later updating:
allParsMixed = ([STL1_4state_multi_1_mix.parameters{:,2},STL1_4state_multi_2_mix.parameters{:,2}]);

% Fit parameters using maximum likelihood estimation:
allParsMixed = combinedModelMixed.maximizeLikelihood(allParsMixed, fitOptions, fitAlgorithm);

% Update model parameters and plot results:
combinedModelMixed = combinedModelMixed.updateModels(allParsMixed, true);
```

### 6 Single-cell RNA sequence data

Samples were obtained from the GSE141834 series [68] to construct a temporal single-cell RNA-sequencing dataset. The dataset comprised eight samples, including cells treated with an ethanol vehicle control and 100 nM dexamethasone for 1, 2, 4, 8, and 18 hours, with biological replicates at the 4- and 8-hour time points. Raw FASTQ files were processed using the kallisto–bustools workflow [?] to generate spliced and unspliced count matrices. Differential expression analysis was performed using DESeq2 [?] on total gene counts, defined as the sum of spliced and unspliced reads. From this analysis, 151 of the 500 most highly expressed genes with positive log fold change were selected for downstream modeling, representing the most strongly upregulated genes by total expression.

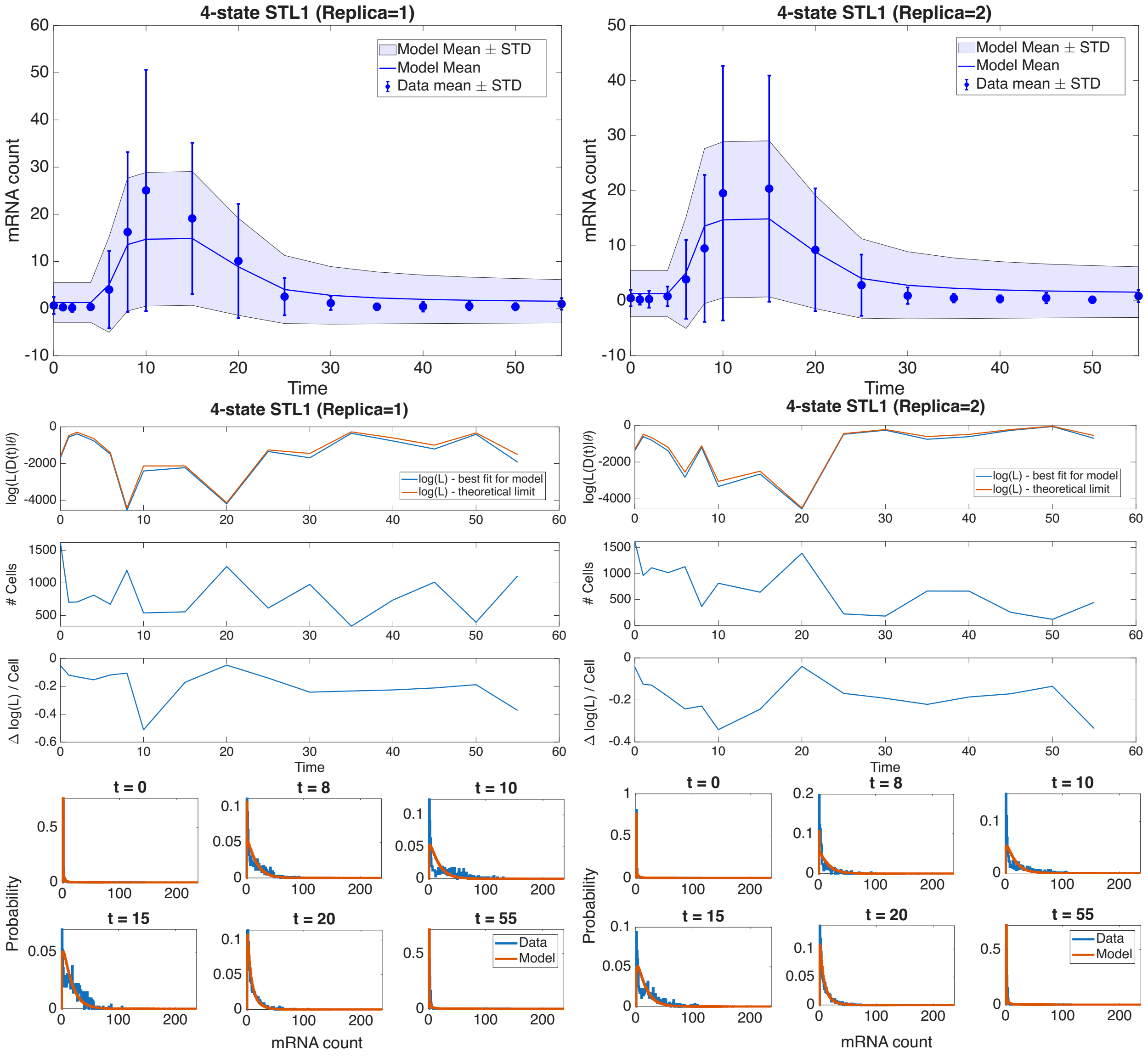


Figure S16: **MultiModel** results from maximum likelihood estimation of 4-state STL1 model (Fig B1 in main text) parameters for (left) "Replica=1" and (right) "Replica=2" experimental data; (top) means and standard deviations (STD) and STL1 data [25], (middle) log-likelihoods ($\log(\mathcal{L}(\mathcal{D}(t)|\theta))$) and number of cells through time, and (bottom) probability distributions by mRNA count, computed using Finite State Projection.

**S1 Table. Properties and functions of SSITMultiModel**

| Property: | Summary: |
|---|---|
| 1. **FIM** | 1. FIM results |
| 2. **SSITModels** | 2. Set of SSIT models and their associated data sets |
| 3. **fspStateSpaces** | 3. Set of FSP state spaces |
| 4. **logLikelihoodFunctions** | 4. Set of functions that compute the log likelihood of the data given each model and provided parameters |
| 5. **parameterConstraints** | 5. Function that applies prior constraints on parameters |
| 6. **parameterIndices** | 6. Set of vectors that map full parameter set to corresponding model |
| 7. **parameters** | 7. Parameter Values |

| Function: | Summary: |
|---|---|
| 1. **addModel** | 1. Add a model or change the definition of the prior |
| 2. **computeFIMs** | 2. This function will compute the individual FIM matrices for |
| 3. **computeTotalLogLikelihood** | 3. Method that computes the total log likelihood of all data and |
| 4. **initializeStateSpaces** | 4. Initialize the FSP state space |
| 5. **maximizeLikelihood** | 5. Search parameter space to determine which sets maximize the likelihood of the data |
| 6. **updateModels** | 6. Updates parameters of the models to provided values and makes |

**S2 Table. MH posterior sample means and standard deviations on the $\log_{10}$ scale, with corresponding moments in linear space assuming $\log_{10}(\theta) \sim \mathcal{N}(\mu, \sigma^2)$ (i.e., $\theta$ is lognormal).**

| Parameter | Mean ($\log_{10}$) | Std ($\log_{10}$) | Mean (lin) | Std (lin) |
|---|---|---|---|---|
| $t_0$ | 0.6643 | 0.0073 | 4.617 | 0.07761 |
| $k_{12}$ | 0.8808 | 0.0313 | 7.62 | 0.5499 |
| $k_{21o}$ | 2.8622 | 0.0390 | 731.1 | 65.78 |
| $k_{21i}$ | 0.5178 | 0.0253 | 3.3 | 0.1924 |
| $k_{23}$ | -0.4147 | 0.0139 | 0.3851 | 0.01233 |
| $k_{34}$ | 0.3439 | 0.0422 | 2.218 | 0.216 |
| $k_{32}$ | 0.7125 | 0.0241 | 5.166 | 0.2869 |
| $k_{43}$ | -0.0642 | 0.0333 | 0.8651 | 0.06643 |
| $d_r$ | -0.6503 | 0.0084 | 0.2238 | 0.004328 |
| $k_{r1}$ | -1.1605 | 0.0155 | 0.06915 | 0.002469 |
| $k_{r2}$ | -0.6631 | 0.0397 | 0.2181 | 0.01998 |
| $k_{r3}$ | 1.9896 | 0.0331 | 97.92 | 7.474 |
| $k_{r4}$ | 1.1843 | 0.0332 | 15.33 | 1.174 |
